# Plaque-associated oligodendrocyte proteostatic failure underlies myelin loss in Alzheimer’s disease

**DOI:** 10.64898/2026.07.08.737318

**Authors:** Jong Shin, Nandita Joshi, Brendan Miller, Shams Vellarikkal, Jianhua Huang, Fan Wu, Yi Cui, Aparna Murali, Joshua Chason, Christopher Scott Campbell, Khoi Chu, Reza Roozafzoon, Mark Dostalík, Jindrich Soukup, Gina Savastano, Xiaolan Shen, Javier Ganz, Shanshan He, Vanessa Peterson, Matthew Kennedy, Iya Khalil, Methodios Ximerakis, Alex M. Tamburino, Rebecca Mathew, Bilal Cakir

## Abstract

Alzheimer’s disease (AD) features amyloid-β plaques and tau pathology, yet the mechanism underlying early and clinically significant myelin loss remains unresolved. Here, we report human iPSC-derived forebrain organoids with doxycycline-inducible expression of SOX10, OLIG2, and NKX6-2 (SON), which generate robust, mature oligodendrocytes and compact myelin in vitro and in vivo. Introducing amyloid precursor protein (APP) pathogenic mutations produces extracellular amyloid-β plaques and phosphorylated tau, accompanied by reduced myelin basic protein (MBP) expression and disrupted myelin ultrastructure. Single-cell and spatial transcriptomics combined with amyloid plaque imaging reveal that oligodendrocytes in plaque-dense regions show pathological changes in calcium signaling, immune activation, lipid remodeling, and protein catabolic pathways. Notably, MBP protein is reduced despite elevated myelin-related transcription, suggesting plaque-induced degradation of myelin proteins. Consistent with this, proteasomal inhibition restores MBP protein levels and improves compact myelin ultrastructure in AD organoids. In human AD brain tissue, protein catabolic pathways are similarly upregulated with increasing plaque density. Together, these findings identify plaque-associated oligodendrocyte proteostatic failure as a candidate mechanism of myelin loss and a potential therapeutic pathway in AD.

---

Alzheimer’s disease (AD) is defined by the accumulation of amyloid-β plaques and neurofibrillary tau tangles, yet myelin loss has emerged as an early and clinically significant feature that has received comparatively little attention. In the AD brain, progressive deterioration of myelin sheaths is detectable across disease stages and correlates with cognitive decline (*1–5*); radiologically, it manifests as white matter hyperintensities (WMH) on MRI, are present in virtually all patients over 60 and frequently precede neuronal atrophy. Despite its prevalence, the cellular mechanism linking amyloid pathology to myelin loss remains poorly understood.

Resolving this requires human model systems that support physiological oligodendrocyte maturation and compact myelination alongside controlled amyloid pathology, a combination not achievable with existing platforms. Two-dimensional cultures lack the 3D architecture required for compact myelination, transgenic mouse models are limited by species-specific differences in glial biology (*6, 7*), and while human brain organoids offer a promising 3D context, existing protocols do not reliably produce mature, myelinating oligodendrocytes at scale (*8–12*).

Here, we address this mechanistic gap by engineering a genetically inducible oligodendrocyte maturation program within human forebrain organoids. Introducing pathogenic APP mutations produces amyloid plaques and recapitulates myelin loss, including reduced MBP protein despite sustained myelination-associated transcription. Using spatial transcriptomics integrated with amyloid plaque imaging, we find that plaque burden reshapes oligodendrocyte transcriptomes in a density-dependent manner, coordinately inducing immune and inflammatory responses, calcium signaling, lipid metabolism, and alterations in proteasomal subunit composition, suggesting a plaque-driven shift in protein catabolism that may contribute to MBP loss. These findings implicate a mechanism by which amyloid pathology drives myelin degeneration through plaque-density-dependent disruption of oligodendrocyte proteostasis.

## Generation of Human Oligodendrocyte-Enriched Forebrain Organoids

To establish a robust platform for studying human oligodendrocytes with myelinating function, we lentivirally engineered XCL-1 hiPSCs with a doxycycline-inducible cassette encoding *SOX10*, *OLIG2*, and *NKX6-2* (SON factors), a combination previously demonstrated to drive stable specification and maturation of oligodendrocytes (*13*) (Fig. 1A, Fig. S1A). We validated the potency of this system by inducing SON expression in suboptimal conditions (embryoid body (EB) and neuronal induction media) lacking specific oligodendrogenic growth factors (Fig. S1B). Within five days, critical lineage markers, such as *MOG* and *PLP1,* were significantly upregulated by approximately 2-fold at the transcription level, confirming the capability of SON factors to drive specific oligodendrocyte identity even in the absence of complex oligodendrocyte growth factor cocktails (Fig. S1C).

**Figure 1.**
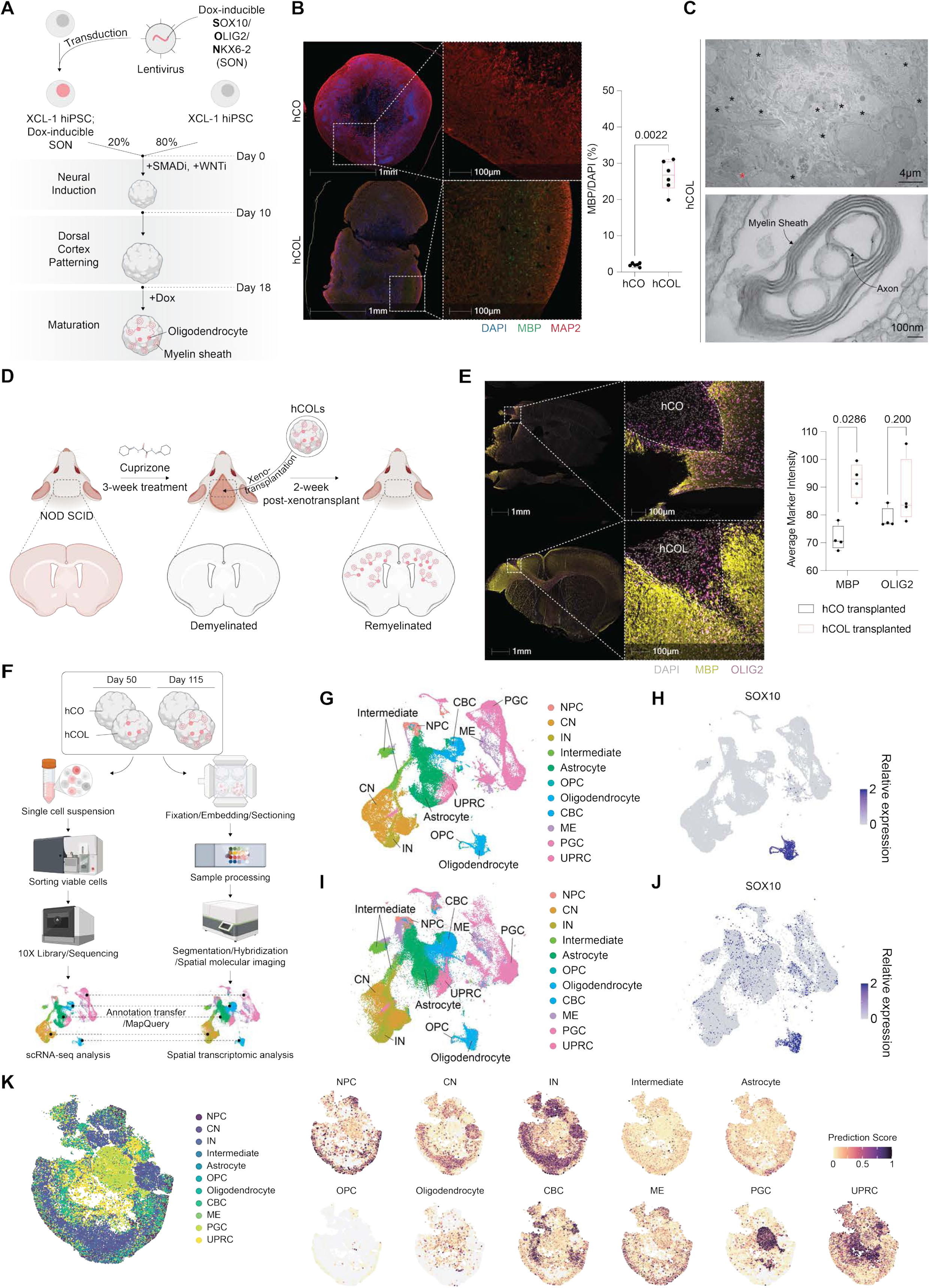
Generation and characterization of oligodendrocyte-enriched human cortical organoids (hCOL). (**A**) Schematic of the protocol to generate oligodendrocyte-enriched human cortical organoids. (**B**) (Left) Immunofluorescence of day 115 human cortical organoids (hCO) and hCOL stained for neurons (MAP2, red) and myelin (MBP, green). Nuclei stained with DAPI (blue). Low-magnification (left, 1 mm) and high-magnification (right, 100 µm). (Right) Box plot of the percentage of MBP+ cells. hCO, 2.0 (mean) ± 0.5 (SD), n=6; hCOL, 26.5 ± 4.2, n=6. Mann-Whitney test. (**C**) Electron microscopy (EM) analysis of hCOL. Low-magnification (top, 4 µm) showing myelinated axons (asterisks) and high-magnification (bottom, 100 nm) showing compact myelin lamellae. hCOL, n=12, from 2 iPSCs from 3 differentiation experiments. (**D**) Schematic experimental design for the xenotransplantation of hCO and hCOL into cuprizone-demyelinated mouse brains. (**E**) (Left) Immunofluorescence of grafted organoids in host mouse brain tissue stained for myelin (MBP, yellow) and oligodendrocyte lineage cells (OLIG2, magenta). hCO (top) and hCOL (bottom) transplants. Low-magnification (left, 1 mm) and high-magnification (right, 100 µm). (Right) Box plots of average marker intensity for MBP and OLIG2. MBP: hCO, 71.6 ± 4.6, n=4; hCOL, 92.4 ± 6.4, n=4. OLIG2: hCO, 78.8 ± 3.7, n=4; hCOL, 87.6 ± 12.3, n=4. Mann-Whitney test. hCO, n=6, and hCOL, n=6, from 3 differentiation experiments. (**F**) Schematic workflow for scRNA-seq and spatial transcriptomic profiling of hCOL. (**G–J**) (**G, H**) scRNA-seq analysis (50 and 115 days of organoid development, n=30,768 cells) and (**I, J**) spatial transcriptomic profiling of hCOL (50 and 115 days, n=31,884 cells). (**G, I**) (Left) UMAP plots of major cell populations and intermediate lineage cells. (**H, J**) (Right) Feature plots showing expression of the pan-oligodendrocyte lineage marker SOX10. NPC, neural progenitor cells; CN, cortical neurons; IN, interneurons; OPC, oligodendrocyte progenitor cells; CBC, cilia-bearing cells; ME, mesenchymal-like cells; PGC, proteoglycan cells; UPRC, unfolded protein responding cells. (**K**) Spatial transcriptomics visualization of day 115 hCOL. (Left) Spatial plot showing the regional distribution of predicted cell types. (Right) Feature plots displaying prediction scores for individual cell types.

We then generated dorsal (human cortical organoid; hCO) and ventral (human medial ganglionic eminence organoid; hMGEO) forebrain organoid models by mixing 20% SON-engineered XCL-1 iPSCs with 80% parental cells, approximating oligodendrocyte abundance (*14*), thereby creating oligodendrocyte-enriched cortical (hCOL) and MGE (hMGEOL) organoids (Fig. S1D). Using a step-wise induction protocol and standard regional patterning, we generated and validated these models as having distinct regional identities: hMGEOLs were enriched for *NKX2-1*, and hCOLs for *PAX6* and vGLUT1 (*SLC17A7*), as reported previously (Fig. S1E) (*13*). Transcriptional and histological analysis further confirmed a robust, time-dependent maturation program. Early lineage markers (*OLIG2*, *PDGFRA*) appeared by day 50, followed by a significant elevation of oligodendrocyte and myelination maturation genes (myelin basic protein *(MBP)*, *MYRF*) and accumulation of OLIG2+ cells by day 115 (Fig. S1F, G). Consistently, SON-induced organoids exhibited robust MBP expression and the formation of compact myelin sheaths wrapping axons, as verified by electron microscopy (EM) (Fig. 1B, C, Fig. S1H, I). Together, these data indicate that SON induction yields mature oligodendrocyte-enriched forebrain organoids that generate compact myelin structures *in vitro*.

To evaluate oligodendrocyte myelinating potential *in vivo*, we used a cuprizone-induced demyelinating paradigm in NOD-SCID immune-deficient mice and transplanted intact hCOLs into the corpus callosum (Fig. 1D). At 14 days post-transplantation, immunostaining for OLIG2 and MBP identified engrafted human oligodendrocyte-lineage cells and increased MBP signals in graft-adjacent regions (Fig. 1E), demonstrating the myelinating potential of hCOL-derived oligodendrocytes *in vivo*.

To comprehensively characterize the cellular ecosystem of our engineered models, we performed single-cell RNA sequencing (scRNA-seq) and spatial transcriptomics (Fig. 1F). In hCOL organoids, scRNA-seq analysis revealed a heterogeneous cellular composition annotated via a previously established hierarchical system and validated by canonical markers (Fig. S2A-E) (*13*). This analysis identified major brain cell types, including excitatory cortical neurons, inhibitory interneurons, and astrocytes, alongside a distinct population of oligodendrocyte lineage cells (OPCs and oligodendrocytes) marked by the exclusive expression of the lineage-specific transcription factor *SOX10* (Fig. 1G, H). Consistent with previous descriptions, we also identified traceable developmental byproduct and stress-associated populations, including cilium-bearing cells (CBC), proteoglycan-expressing cells (PGC), and unfolded protein response cells (UPRC) (Fig. 1G) (*15*). Spatial transcriptomic profiling mapped these populations to the organoid architecture, demonstrating that oligodendrocytes are dispersed throughout the central core, underlying the neuron-rich outer layer, rather than restricted to isolated clusters (Fig. 1I–K, Fig. S3A-D). Parallel analysis of hMGEOLs confirmed the presence of all major cell types, with a notably higher proportion of interneurons compared to hCOL (Fig. S4A-G, S5A-D) (*13, 16, 17*). Electrophysiological recordings confirmed that both models establish active neuronal networks with both excitatory and inhibitory spiking activity (Fig. S1J) (*18*).

Although oligodendrocyte lineage cells were detectable in hMGEOLs as early as day 50, their population was less abundant and sustained compared to hCOLs, potentially due to region-specific developmental constraints in the ventral forebrain context (*19*). We therefore prioritized the hCOL model for subsequent disease modeling to examine how amyloid pathology alters oligodendrocyte maturation programs and myelin integrity.

## Familial APP Mutations Produce Amyloid Pathology and Myelin Deficits in hCOL

Next, to investigate the impact of amyloid pathology on human oligodendrocytes, we generated hCOL and hMGEOL AD organoids by mixing 80% XCL-1 iPSCs carrying knock-in familial APP Swedish/Indiana mutations with 20% SON-engineered XCL-1 iPSCs carrying wild-type APP (hCOL^APP^ and hMGEOL^APP^) (Fig. 2A, Fig. S6A, B). By day 115, both mutant organoid models exhibited AD-like pathology, including extensive accumulation of extracellular amyloid-β plaques (Fig. 2B, Fig. S6C) and intracellular phosphorylated Tau aggregates (Fig. 2C, Fig. S6D). These pathological features were accompanied by a marked reduction in MBP expression compared to isogenic controls, suggesting impaired myelin integrity (Fig. 2D). EM and morphometric analyses indicated increased myelin disruption in hCOL^APP^ (Fig. 2E, Fig. S6F). Consistent with this, Luxol Fast Blue (LFB) staining showed reduced myelin density (Fig. S6E). Notably, APP transcript levels were not elevated in hCOL^APP^ relative to hCOL, rather reduced at day 115 (Fig. 2G), confirming that amyloid accumulation in this model is driven by the pathogenic processing of the Swedish and Indiana mutations rather than transcriptional upregulation of APP (*20–23*). Importantly, the SON-induced oligodendrocytes themselves carry wild-type APP; the myelin damage therefore arise not from cell-intrinsic mutant APP expression but from the aberrant accumulation of mutant APP-derived amyloid-β in the microenvironment. Together, these data demonstrate that these models effectively recapitulate key features of the myelin degeneration observed in clinical AD and murine models (*24–26*).

**Figure 2.**
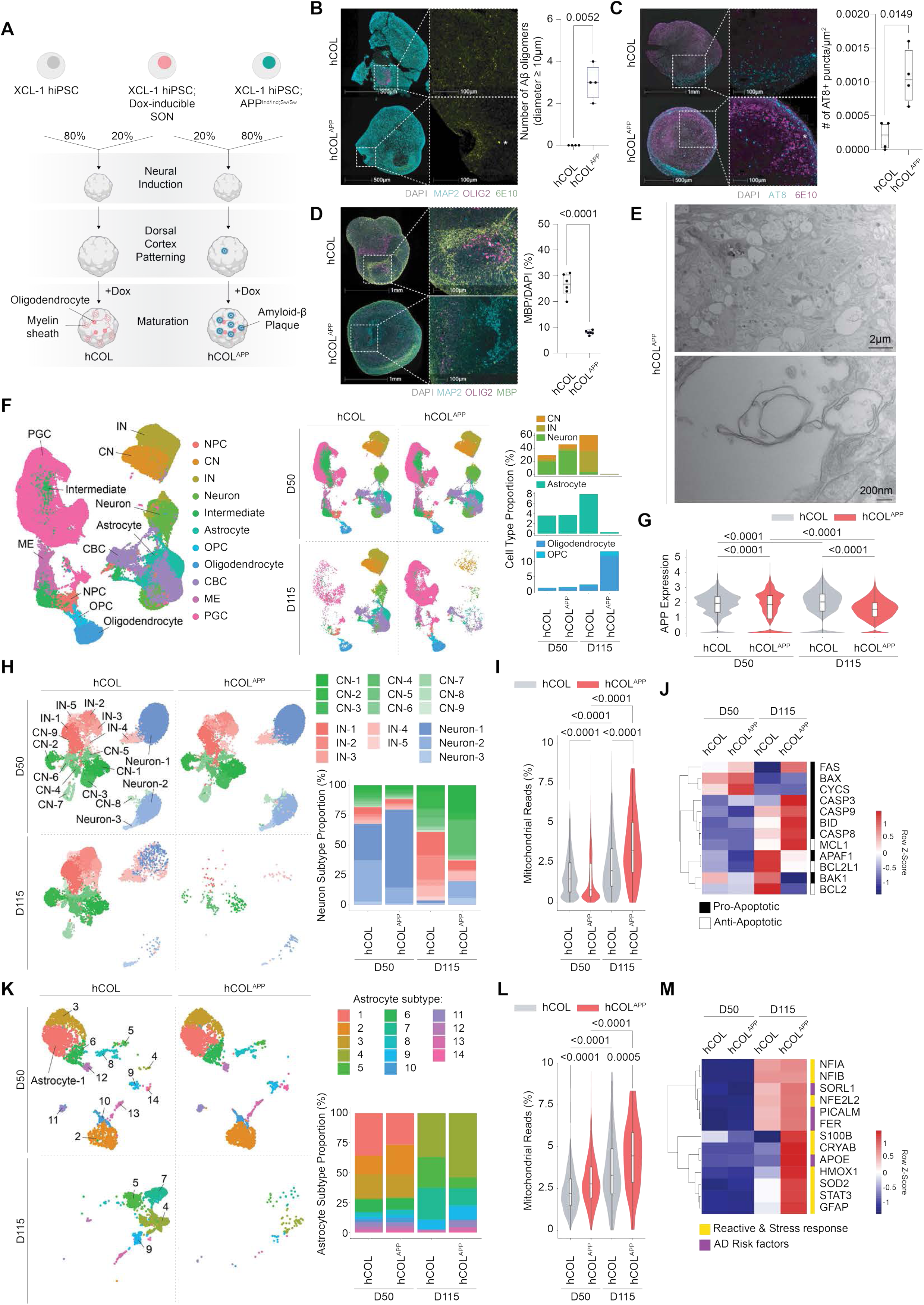
Recapitulation of AD-like pathology and transcriptomic signatures in hCOL^APP^. (**A**) Schematic of the generation of AD oligodendrocyte-enriched human cortical organoid (hCOL^APP^). (**B**) (Left) Immunofluorescence of amyloid-β aggregation. Day 50 hCOL (top) and hCOL^APP^ (bottom) stained for amyloid-β (6E10, green), neurons (MAP2, cyan), and oligodendrocyte lineage cells (OLIG2, magenta). Nuclei stained with DAPI (grey). Low-magnification (left, 500 µm) and high-magnification (right, 100 µm). (Right) Box plot of the number of amyloid-β oligomers (≧10 µm in diameter). hCOL, 0 ± 0 (Not detected), n=4; hCOL^APP^, 3.00 ± 0.82, n=4. Welch’s t-test. (**C**) (Left) Immunofluorescence of tau pathology. Day 120 hCOL (top) and hCOL^APP^ (bottom) stained for phosphorylated tau (AT8, cyan) and amyloid-β (6E10, magenta). Nuclei stained with DAPI (grey). Low-magnification (left, 1 mm for top and 500 µm for bottom) and high-magnification (right, 100 µm). (Right) Box plot of the number of AT8+ puncta per µm^2^ area. hCOL, 0.0002 ± 0.0002, n=4; hCOL^APP^, 0.001 ± 0.0004, n=4. Welch’s t-test. (**D**) (Left) Immunofluorescence of myelin loss. Day 115 hCOL (top) and hCOL^APP^ (bottom) stained for neurons (MAP2, cyan), oligodendrocyte-lineage cells (OLIG2, magenta), and mature oligodendrocyte (MBP, green). Low-magnification (left, 1 mm) and high-magnification (right, 100 µm). (Right) Box plot of the percentage of MBP+ cells. hCOL, 26.50 ± 4.25, n=6; hCOL^APP^, 7.97 ± 0.81, n=6. Welch’s t-test. (**E**) Electron microscopy analysis of hCOL^APP^. Low-magnification (top, 2 µm) and high-magnification (bottom, 100 nm). hCOL^APP^, n=12 organoids from 1 iPSC from 3 differentiation experiments. (**F**) scRNA-seq analysis of hCOL and hCOL^APP^ developed for 50 days (D50) and 115 days (D115). (Left) UMAP of mature and differentiating brain cell types. (Middle) UMAPs of hCOL and hCOL^APP^ at day 50 and day 115, individually (D50 hCOL, n=69,980; D115 hCOL, n=24,623; D50 hCOL^APP^, n=88,425; D115 hCOL^APP^, n=24,204). (Right) Stacked bar plots of cell type proportions of neurons (top), astrocytes (middle), and oligodendrocytes/OPCs (bottom). NPC, neural progenitor cells; CN, cortical neurons; IN, interneurons; OPC, oligodendrocyte progenitor cells; CBC, cilia-bearing cells; ME, mesenchymal-like cells; PGC, proteoglycan cells. (**G**) Violin plots of APP transcript level in hCOL and hCOL^APP^ at day 50 and day 115. Kruskal-Wallis test. (**H-J**) Neuronal transcriptomic analysis of hCOL and hCOL^APP^ at day 50 and day 115. (**H**) (Left) UMAPs of neuron subtypes (D50 hCOL, n=20,898; D115 hCOL, n=14,958; D50 hCOL^APP^, n=41,101; D115 hCOL^APP^, n=320). (Right) Stacked bar plots of neuron subtype proportions. (**I**) Violin plots of mitochondrial read percentages. Kruskal-Wallis test. (**J**) Heatmap of pro- and anti-apoptosis genes (row z-score). (**K-M**) Astrocyte transcriptomic analysis of hCOL and hCOL^APP^. (**K**) (Left) UMAPs of astrocyte subtypes (D50 hCOL, n=2,529; D115 hCOL, n=1,960; D50 hCOL^APP^, n=3,295; D115 hCOL^APP^, n=89). (Right) Stacked bar plots of astrocyte subtype proportions. (**L**) Violin plots of mitochondrial read percentages. Kruskal-Wallis test. (**M**) Heatmap of reactive astrocyte markers, stress response genes, and AD risk factors (row z-score).

To define the cellular and molecular landscape associated with this pathology, we performed scRNA-seq profiling of hCOL and hCOL^APP^ at day 50 and 115, revealing a global shift in cellular states associated with disease progression (Fig. 2F, Fig. S7A-G). While cellular composition was comparable between conditions at day 50, by day 115 we observed a marked reduction in both neuronal and astrocyte proportions in hCOL^APP^, accompanied by a relative increase in the proportion of oligodendrocytes (Fig. 2F). Neuronal subclustering revealed a widespread loss affecting all neuron types, including a notable decrease in interneuron populations. Moreover, surviving neurons exhibited an elevated mitochondrial gene fraction and upregulation of pro-apoptotic genes, indicative of ongoing apoptosis (Fig. 2H–J, Fig. S8A-C). Similarly, astrocytes in hCOL^APP^ transitioned to a reactive state alongside signs of metabolic reprogramming (Fig. 2K–M, Fig. S8E–I). Collectively, these findings establish that the hCOL^APP^ model recapitulates the broad multicellular AD pathology, establishing a relevant context for dissecting oligodendrocyte dysfunction.

While neurons underwent apoptosis and astrocytes became reactive, oligodendrocytes in hCOL^APP^ exhibited a complex, apparently discordant phenotype. Differential expression analysis revealed a profound upregulation of myelin biosynthesis pathways, including fatty acid, cholesterol, ceramide, and galactolipid synthesis genes (Fig. S9A, B, F, G, Table S1), suggesting a putative compensatory myelin biosynthetic drive. However, this transcriptional upregulation of myelin synthesis appeared uncoupled from central energy metabolism and translational capacity. For example, we observed a reduction in central carbon metabolism, including broad reductions in glycolysis and TCA cycle genes, and a broad downregulation of ribosomal and translation machinery (Fig. S9F–I). Instead, the proteostatic network shifted toward catabolism and immune response, evidenced by the upregulation of antigen processing pathways and protein catabolic pathways (Fig. S9H, I). This disparity is consistent with an amyloid-rich environment that drives oligodendrocytes into a metabolically compromised and immune-reactive state in which the transcriptional programs for myelination may not be energetically supported or translated into functional protein.

## Aberrant Oligodendrocyte Maturation in hCOL^APP^

To better understand the oligodendrocyte response and changes in oligodendrocyte states within the AD environment at single-cell resolution, we performed subclustering and identified oligodendrocyte subtypes (Fig. 3A, Fig. S10A–C). While day 50 organoids were primarily composed of early-stage progenitor populations in both conditions, hCOL^APP^ organoids exhibited a dramatic shift in cellular composition by day 115 (Fig. 3B, Fig. S10F). Slingshot pseudotime analysis indicated that hCOL^APP^ oligodendrocytes at day 115 had significantly higher pseudotime scores than controls, placing them at a later position along the inferred trajectory despite impaired myelin integrity (Fig. 3C, Fig. S10D, E). Indeed, differential expression analysis confirmed that the specific subtypes enriched in hCOL^APP^ (e.g., subtype 9) expressed higher levels of oligodendrocyte maturation and myelination genes compared to the subtypes enriched in controls (e.g., subtype 5) (Fig. 3D, E, Fig. S10A, B, Table S2). LOESS trajectory modeling revealed that hCOL^APP^ oligodendrocytes bypassed subtype 5, a WNT-high/GSK3β-low state we termed the “Reserve” subtype, while retaining the developmentally subsequent WNT-low subtype 8 (Fig. S10G, H). This suggests that the amyloid environment is associated with attenuation of WNT-associated programs, leading to reduced occupancy of this developmental state and a shift toward an aberrant differentiation program (Fig. S10I). Detailed molecular characterization of these AD-enriched subtypes revealed a discordant cellular state with upregulation of myelin biosynthetic transcripts despite a suppression of central carbon metabolism and a concurrent shift toward protein catabolism and immune response (Fig. S10J). This transcriptional–metabolic discordance provides a plausible molecular basis for the observed uncoupling: diversion of metabolic and proteostatic resources toward catabolic stress responses prevents successful translation, accumulation, and assembly of myelin proteins into mature sheaths.

**Figure 3.**
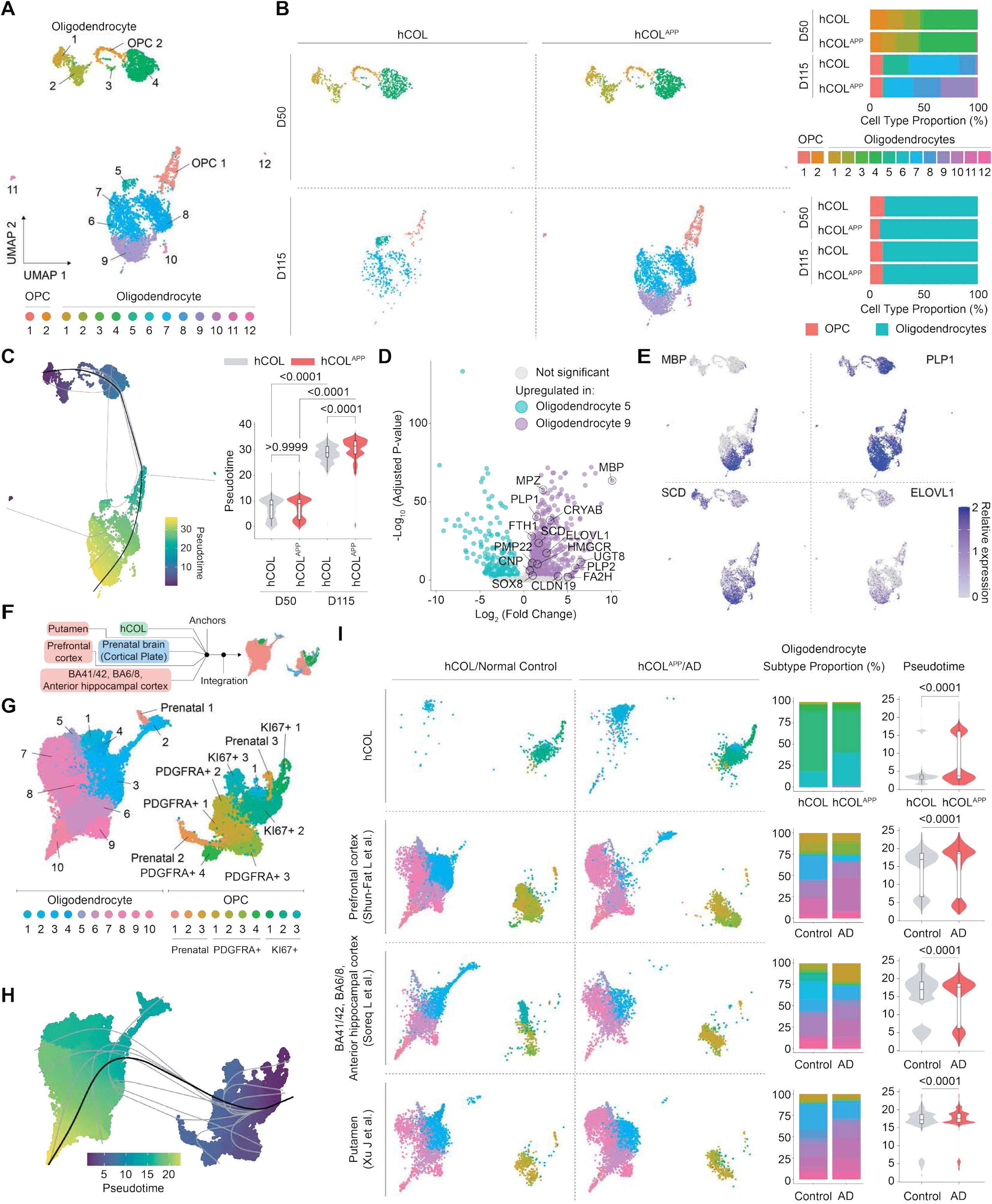
Impaired oligodendrocyte maturation and altered developmental trajectory in hCOL^APP^. (**A**) UMAP visualization of oligodendrocyte and OPC subtypes in hCOL and hCOL^APP^. (**B**) (Left) Individual UMAPs of oligodendrocyte and OPC subtypes in hCOL and hCOL^APP^ at D50 and D115 (D50 hCOL, n=799; D115 hCOL, n=561; D50 hCOL^APP^, n=1,233; D115 hCOL^APP^, n=3,287). (Right) Stacked bar plots showing oligodendrocyte and OPC subtypes (top) and cell type proportions (bottom). (**C**) (Left) Pseudotime trajectory analysis of the oligodendrocyte lineage. (Right) Violin plots showing pseudotime scores of hCOL and hCOL^APP^ at day 50 and day 115. Kruskal-Wallis test. (**D**) Volcano plot of differentially expressed genes highlighting genes upregulated in oligodendrocyte subtype 5 (turquoise, shifted to the left) and subtype 9 (purple, shifted to the right). Oligodendrocyte maturation genes are labeled. (**E**) Feature plots of the oligodendrocyte maturation genes differentially enriched in oligodendrocyte subtype 9. (**F**) Schematic of the integration of hCOL^APP^ scRNA-seq data with human AD and prenatal metadata. (**G**) UMAP of hCOL/hCOL^APP^ oligodendrocyte and OPC subsets integrated with AD and prenatal human datasets, annotated with oligodendrocyte and OPC subtype. (**H**) Pseudotime trajectory analysis of the integrated dataset. (**I**) (Left) Individual UMAPs of oligodendrocyte and OPC subsets comparing control and AD conditions across datasets. Top: hCOL (this study). Middle: Prefrontal cortex (Shun-Fat et al.) and B41/42, BA6/8, Anterior hippocampal cortex (Soreq et al.). Bottom: Putamen (Xu et al.). (Middle) Stacked bar plots of oligodendrocyte and OPC subtype proportions. (Right) Violin plots of pseudotime scores comparing control and AD conditions across datasets. Kolmogorov-Smirnov test.

To validate the clinical relevance of these findings, we integrated our organoid scRNA-seq data with three independent human AD single-nucleus RNA-seq datasets covering distinct brain regions (prefrontal cortex, B41/42-BA6/8-anterior hippocampus, and putamen), alongside human prenatal brain data to provide a comprehensive developmental context and identification (Fig. 3F) (*27–30*). In the integrated oligodendrocyte lineage space, subclustering and pseudotime trajectory analysis revealed a conserved differentiation landscape with distinct segregation of oligodendrocyte subpopulations from OPCs (Fig. 3G–H). Analysis of individual human datasets confirmed a shift in cellular composition characterized by neuronal loss and an expansion of the oligodendrocyte lineage, consistent with the organoid model (Fig. S11A-F). However, unlike the relative astrocyte fractions reduced in hCOL^APP^ organoids, astrocyte populations were expanded in human AD datasets, consistent with well-documented reactive astrogliosis in AD (*31*). Crucially, we observed specific alterations in oligodendrocyte subtype composition and differentiation dynamics in AD, mirroring the shifting pattern seen in the organoid models (Fig. 3B, I). Detailed characterization identified distinct functional states: for instance, subtype 3 (homeostatic mature oligodendrocytes) was depleted, while subtype 7 (a stressed, reactive mature state) was significantly enriched in AD (Fig. S12A–F). Differential expression analysis between these populations further revealed that the AD-enriched subtype 7 displays elevated lipid and myelin biosynthetic transcripts alongside altered proteostasis and central carbon metabolism, particularly mitochondrial respiration, a transcriptional pattern also seen in the organoid models (Fig. S12G-I).

In developing organoids, this manifests as an accelerated but abnormal maturation trajectory; in the adult human brain, where oligodendrocytes are largely mature, the same transcriptional signature may reflect stress-associated maintenance failure rather than increased differentiation.

## Plaque-associated transcriptional reprogramming in hCOL^APP^ implicates proteostatic disruption

Having established that oligodendrocyte subtype composition is shifted in both the hCOL^APP^ organoid model and in previously published human AD single-nucleus datasets, we next asked whether these shifts are directly driven by local amyloid plaque burden. Since the hCOL^APP^ model is defined by familial APP mutations that rapidly accelerate amyloid accumulation, we reasoned that the subtype shifts observed at the transcriptomic level may reflect a plaque-proximal cellular response. To test this, we integrated spatial transcriptomics with immunofluorescence-based plaque imaging (Fig. 4A, Fig. S13A-D). Using an in-house deep-learning semantic segmentation model, we computed plaque density and distance to the nearest plaque for the centroid of each cell and registered these metrics directly to the spatial gene expression profiles of each cell (Fig. 4A, Fig. S14A, B). hCOL^APP^ organoids exhibited markedly higher plaque density than isogenic controls (Fig. 4A), confirming pathogenic amyloid accumulation in the disease model. Furthermore, a substantially larger proportion of oligodendrocyte lineage cells in hCOL^APP^ were located in close proximity to plaques compared to controls (Fig. S14C, D). Consistent with this, plaque density and proximity showed a strong inverse correlation across both conditions (Fig. S14E), validating that these two metrics capture related but complementary aspects of the local amyloid microenvironment. Together, these spatial analyses indicate that plaque density and plaque proximity provide complementary measures of local amyloid burden.

**Figure 4.**
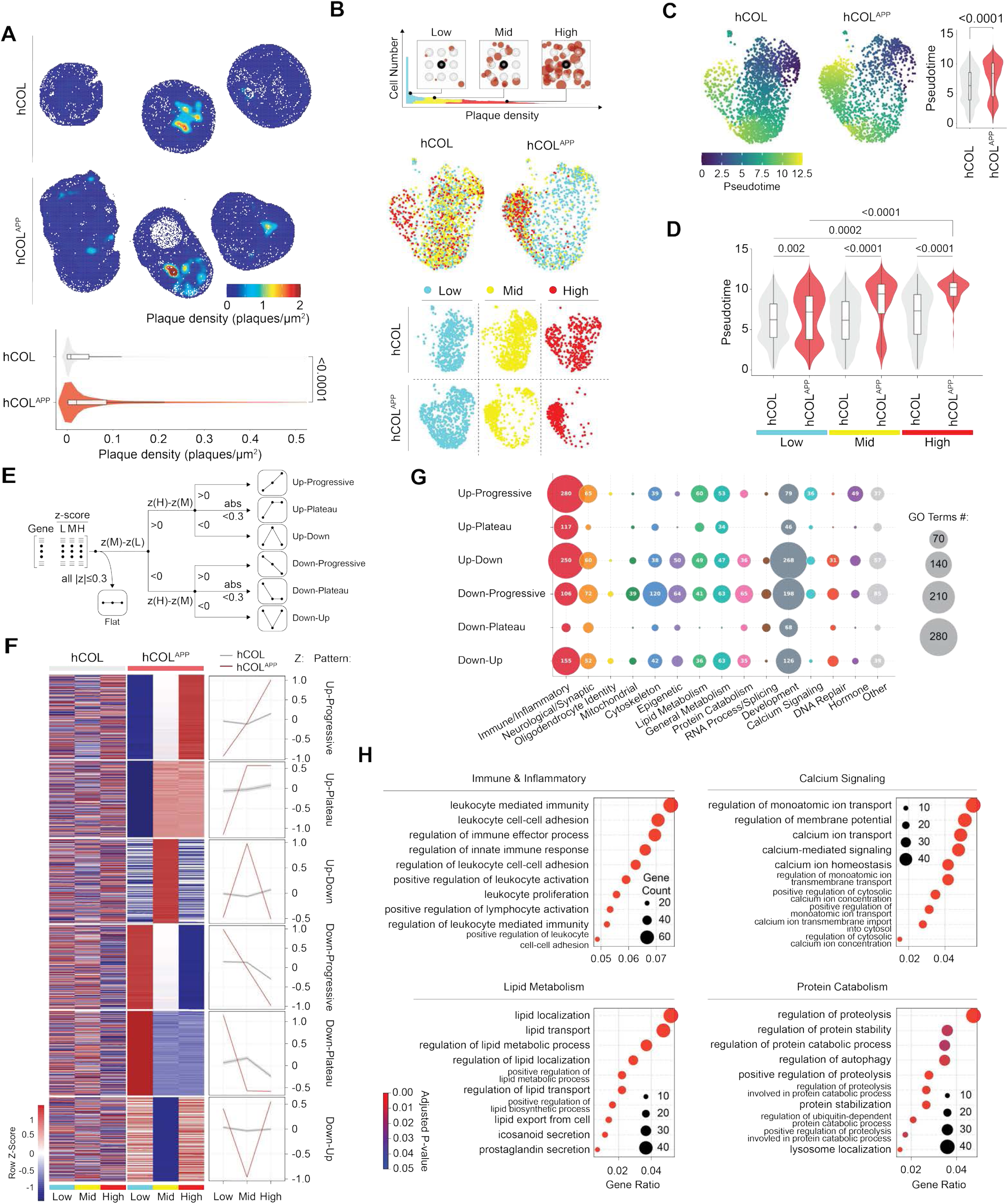
Plaque-associated transcriptional reprogramming in hCOL^APP^ implicates proteostatic disruption. (**A**) (Top) Spatial visualization of amyloid-β plaque density derived from 6E10 immunofluorescence of hCOL and hCOL^APP^. (Bottom) Violin plot of plaque density in hCOL and hCOL^APP^ (zoomed view). hCOL, 0.008282 plaques/μm^2^ (median) with an IQR of 0.0003-0.048; hCOLAPP, 0.02052 plaques/μm^2^ with IQR of 0.001804 and 0.08631. Wilcoxon test. (**B**) (Top) Histogram of all cell types from hCOL and hCOL^APP^ organoids used for spatial transcriptomics, stratified by plaque density tertile (Low, cyan; Mid, yellow; High, red). (Middle) UMAP of subclustered OPC/oligodendrocyte populations colored by plaque density tertile. hCOL, left. hCOL^APP^, right. (Bottom) UMAPs split by both plaque density tertile and group, colored by plaque density tertile. (**C**) (Left) UMAPs of OPC/oligodendrocytes from hCOL and hCOL^APP^ colored by Slingshot pseudotime score. (Right) Violin plot comparing pseudotime scores of OPC/oligodendrocytes between hCOL and hCOL^APP^. Wilcoxon test. (**D**) Violin plots of pseudotime scores stratified by plaque density tertile (Low, Mid, High) in hCOL and hCOL^APP^. Kruskal-Wallis test. (**E**) Decision tree illustrating the classification of genes into six plaque density-dependent trajectory patterns based on z-score expression across tertiles (Low, L; Mid, M; High, H). Only genes from oligodendrocytes. Genes where all |z| ≤ 0.3 are classified as Flat and excluded from downstream analysis. Remaining genes are first split by the sign of z(M)−z(L): genes with z(M)−z(L) > 0 enter the upregulated branch; genes with z(M)−z(L) < 0 enter the downregulated branch. Within each branch, the sign and magnitude of z(H)−z(M) determines the final pattern: if z(H)−z(M) > 0, the gene is Progressive; if |z(H)−z(M)| < 0.3, the gene is Plateau; if z(H)−z(M) < 0, the gene is Up-Down or Down-Up. This yields six mutually exclusive patterns: Up-Progressive, Up-Plateau, Up-Down, Down-Progressive, Down-Plateau, and Down-Up. (**F**) (Left) Heatmap of top 300 differentially expressed genes of oligodendrocytes across plaque density tertile in hCOL and hCOL^APP^ (row z-score)., grouped by trajectory pattern. (Right) Line plots of average z score per pattern across plaque density tertiles in hCOL (grey) and hCOL^APP^ (red). (**G**) Bubble plots of Gene Ontology (GO) biological process enrichment for genes assigned to each of the six trajectory patterns exclusively in hCOL^APP^ oligodendrocytes. (**H**) Dot plots of GO enrichment for the Up-Progressive pattern, showing the top 10 significant pathways per category relevant to oligodendrocyte biology and myelination loss.

To examine whether oligodendrocyte states scale with plaque burden, we stratified all cells into three groups by plaque density tertile (Low, Mid, and High) (Fig. 4B, top) and analyzed composition (Fig. 4B, middle and bottom) and pseudotime (differentiation) scores of OPC/oligodendrocyte subtypes (Fig. 4C). In hCOL^APP^ organoids, OPC and oligodendrocyte subclusters formed distinct groupings across density tertiles, with “mature” subtypes progressively enriched in higher-density regions (Fig. 4B, middle and bottom). Consistent with this, hCOL^APP^ OPC/oligodendrocytes showed significantly higher pseudotime scores than hCOL controls overall, and this elevation was present and progressively increased across all three density tertiles (Fig. 4C, D), mirroring the differentiation advancement documented in the single-cell data (Fig. 3C). Cells in high-density regions were also preferentially located in close proximity to plaques (Fig. S14F, H). However, when pseudotime was stratified by proximity tertile rather than density tertile, the relationship was markedly weaker: the pseudotime difference between hCOL and hCOL^APP^ was non-significant in the distal bin (Fig. S14G), indicating that cumulative local plaque burden, as captured by density, is a stronger predictor of oligodendrocyte transcriptional state than distance to the nearest single plaque. This distinction likely arises because proximity to one plaque fails to account for the overlapping fields of multiple neighboring plaques, which are better reflected by the density metric.

The plaque density-dependent subtype and differentiation shifts suggested the existence of transcriptional programs systematically regulated by local plaque burden. To identify these, we developed a branching classification algorithm that assigns each gene a trajectory pattern based on the direction and magnitude of its z-score change across the three density bins (Fig. 4E). This framework defines six patterns: Up-Progressive (monotonically increasing with density), Up-Plateau (early increase followed by stabilization), Up-Down (transient increase then decline), Down-Progressive (monotonically decreasing), Down-Plateau (early decrease followed by stabilization), and Down-Up (transient decrease then recovery). Applying this framework to hCOL and hCOL^APP^ separately revealed that the conditions diverge substantially across all six pattern categories, with the large majority of density-dependent genes in each pattern showing condition-exclusive trajectories and only a small shared fraction (Fig. 4F, Fig. S15A, Table S3). This divergence was particularly striking for Up-Plateau and Down-Plateau, where fewer than 3% of genes were assigned to the same pattern in both conditions, but was broadly evident across up- and downregulated patterns alike (∼15% overlap). The pervasive condition-specificity of these trajectories indicates that the transcriptional landscape shaped by plaque density is fundamentally distinct in the disease context, reflecting active, disease-specific reprogramming selectively engaged by pathological amyloid levels. The plaque-responsive transcriptional programs in hCOL^APP^ are qualitatively distinct from those in isogenic controls and cannot be explained as an amplification of density-dependent expression patterns already present under non-pathological conditions.

To highlight the disease-specific biological pathways of these plaque-associated gene sets, we performed Gene Ontology (GO) pathway enrichment analysis for each hCOL^APP^-specific gene set across all six pattern categories. Enrichment was broadly distributed across patterns, consistent with a systemic transcriptional reorganization in response to escalating plaque-mediated stress (Fig. 4G, Fig. S15B, Table S4). Examining the pattern-specific distribution of GO categories revealed a notable concentration of immune and inflammatory response, calcium signaling, and lipid metabolism pathways within the Up-Progressive gene set - those most strongly and monotonically induced by plaque density (Fig. 4G, Fig. S15C). Protein catabolic pathways were similarly enriched across all six pattern categories, pointing to a pervasive and pattern-independent disruption of proteostatic balance in plaque-dense regions (Fig. 4G, H).

The pervasive enrichment of protein catabolic pathways across all pattern categories, combined with the density-dependent alteration of proteasomal subunit composition at the transcript level, is notable given the MBP protein deficit observed in hCOL^APP^ despite elevated myelination-associated transcripts (Fig. 2D, Fig. 3D, E). Increased transcription does not guarantee protein accumulation when proteostatic control is disrupted. The co-occurrence of these catabolic programs with calcium signaling, immune and inflammatory responses, and lipid metabolism gene sets, all of which were induced in a plaque density-dependent manner, suggests that plaque-driven cellular stress converges on proteostatic disruption as a mechanism underlying MBP loss.

## Plaque-associated oligodendrocyte proteostatic stress is conserved in human AD brain

To determine whether the plaque-associated oligodendrocyte transcript reprogramming and protein catabolic signature identified in hCOL^APP^ are recapitulated in human AD brain, we obtained formalin-fixed paraffin-embedded (FFPE) tissue blocks of Brodmann area 39 (BA39) from the NIH NeuroBioBank, serially sectioned, and processed them for immunohistochemistry against amyloid-β (6E10) and spatial transcriptomics profiling covering approximately 19,000 genes on the same tissue section (Fig. 5A). The cohort comprised six unaffected normal control donors and seven AD donors spanning the full spectrum of neuropathological severity (Braak stages I-VI, Thal phases 1-3, and CERAD scores 1-3) with similar gender and age distributions across both groups (Fig. S16A, Table S5).

**Figure 5.**
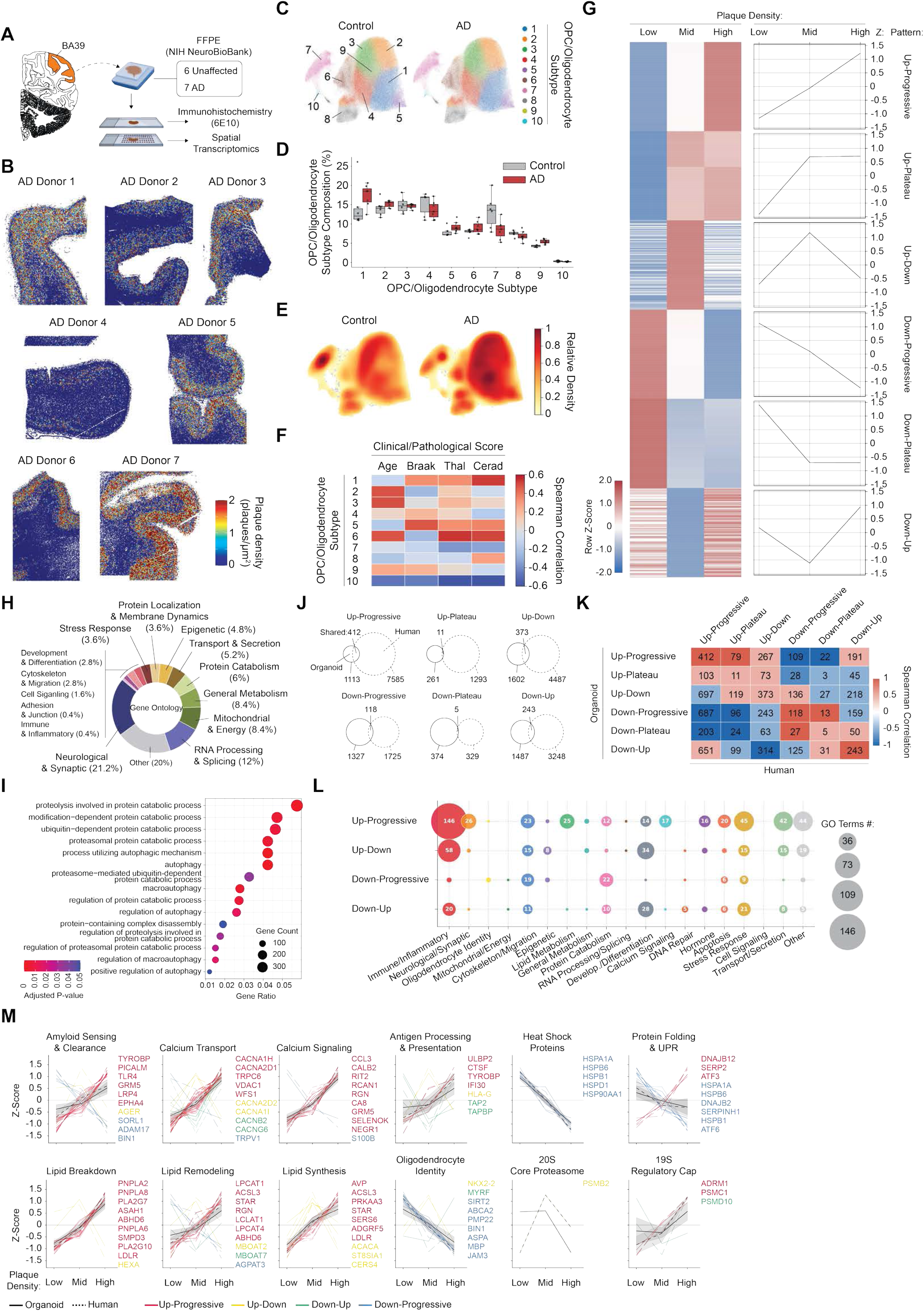
Plaque-associated oligodendrocyte proteostatic stress is conserved in human AD brain. (**A**) Schematic of experimental workflow. FFPE brain tissue from BA39 (NIH NeuroBioBank; 6 unaffected control, 7 AD donors) processed for 6E10 immunohistochemistry and spatial transcriptomics (**B**) Spatial visualization of amyloid-β plaque density across seven AD donor brain sections derived from 6E10 immunofluorescence. (**C**) UMAP of subclustered OPC/oligodendrocyte populations from control and AD human spatial transcriptomics. (**D**) Box plots of OPC/oligodendrocyte subtype composition comparing control (grey) and AD (red) across subtypes (**E**) Kernel Density Estimate (KDE) analysis showing cell density distribution in UMAP comparing control and AD. (**F**) Heatmap of Spearman correlation between OPC/oligodendrocyte subtype proportions and clinical/pathological scores (Age, Braak, Thal, CERAD) across subtypes (**G**) (Left) Heatmap of top 300 differentially expressed oligodendrocyte genes across plaque density tertile (Low, Mid, High) in human AD donor tissues, grouped by trajectory pattern (row Z-score). (Right) Line plots of average z-score per pattern across plaque density tertiles. (**H**) Donut chart of GO biological process categories significantly associated with genes from Up-Progressive pattern (**I**) Dot plot of GO enrichment for protein catabolism-related pathways based on the genes from the Up-Progressive pattern. (**J**) Venn diagrams showing the overlap of plaque-associated genes between organoid and human AD brain tissues for each of six trajectory pattern categories. (**K**) Matrix of Spearman correlations of plaque-associated gene trajectories between organoid and human AD brain tissues across all six pattern category combinations. (**L**) Bubble plots of GO biological process enrichment across pattern categories conserved between organoid and human AD brain tissues. (**M**) Line plots of z-score expression across plaque-associated tertile for canonical gene sets grouped by biological processes including amyloid amyloid sensing, calcium signaling, lipid metabolism, proteostasis, and oligodendrocyte identity. Only genes in overlap between organoid and Human AD brain tissues.

Spatial transcriptomic analysis successfully resolved all major brain cell types across all samples in both groups (Fig. S16B–F, Fig. S17). Cell type compositions were broadly similar between control and AD tissue, with subtle but notable differences: AD samples showed modestly lower neuronal and OPC proportions accompanied by a relative increase in the oligodendrocyte compartment, an observation consistent with the expanded oligodendrocyte lineage seen throughout our organoid and human metadata scRNA-seq analyses (Fig. 2F, Fig. S16B–D, Fig. S11D). Consistent with the organoid findings, MBP transcript levels were substantially elevated in AD oligodendrocytes compared to controls (Fig. S16E), reinforcing the transcript–protein discordance at the center of this study.

Using the same algorithm applied to the organoids, we computed plaque density and proximity and registered these metrics to all cells in the tissues we used for spatial transcriptomics (Fig. 5B, Fig. S18A). Unlike in the organoids, plaques were predominantly concentrated in grey matter cortical layers and more broadly distributed into the underlying white matter, where oligodendrocyte density is high. As in the organoid model, plaque density was strongly inversely correlated with distance to the nearest plaque (Fig. S18B). Stratifying cells into plaque density tertiles (Fig. S18C) and focusing on the oligodendrocyte compartment, subtype compositional shifts in human AD were considerably more modest than in the organoid model or in other scRNA-seq meta-analyses (Fig. 5C). Nevertheless, quantitative subtype composition analysis revealed differential abundances between conditions (Fig. 5D, E), and AD-enriched subpopulations, particularly oligodendrocyte subtypes 1 and 5, showed positive correlations with neuropathological staging scores including Braak stage, Thal phase, and CERAD score, confirming disease relevance of the observed subtype changes (Fig. 5F).

To identify how gene expression responds to local plaque load in human AD oligodendrocytes, we applied the same plaque density tertile-based pattern classification across all six categories (Fig. 5G, Table S6). In contrast to the organoid model, where pathway enrichment was broadly distributed across all six pattern categories, in human AD tissue the most biologically significant pathways were concentrated within the Up-Progressive category (Fig. 5H, Fig. S18D, E, Table S7). Enriched pathways included neurological and synaptic processes, consistent with active remodeling of oligodendrocyte–neuron interactions (*32*); RNA processing and splicing, reflecting broad transcriptional reprogramming that has recently emerged as a key feature of glial stress responses (*33*); mitochondrial metabolism indicative of energetic compromise (*34*); transport and secretion processes suggesting active cellular remodeling and paracrine signaling to neighboring cells (*35*); and protein catabolism (Fig. 5I, Fig. S18E). Each of these programs has been independently implicated in AD, and their convergent enrichment specifically in Up-Progressive oligodendrocytes provides corroboration in human tissue of the plaque-associated stress signature identified in the organoid model.

To systematically compare plaque-associated transcriptional programs between the organoid and human contexts, we identified genes overlapping in both across pattern categories (Fig. 5J, Table S8). While the absolute overlap was moderate, reflecting the inherent differences between model systems as well as technical factors including different spatial transcriptomics platforms and gene panel sizes and different dynamic ranges of plaque density, genes shared between models showed notable directional concordance within matching pattern categories (Fig. 5K). For each shared gene, concordance was quantified as the Spearman correlation of its three-point z-score trajectory across plaque density tertile between organoid and human tissues. Genes assigned to the same pattern category in both models showed highly concordant density-dependent trajectories, while genes in different patterns showed variable correlation. Upregulation patterns retained a subset of positively correlated genes even among mismatches, consistent with inherent sensitivity of discrete classification near threshold boundaries, whereas downregulation patterns showed the strongest anticorrelation, indicating that the classification is most definitive for genes declining with plaque density. Conservation of trajectory shape across model systems was largely confined to Up-Progressive and Up-Plateau genes; however, given the small number of overlapping genes in the Up-Plateau category, Up-Progressive emerges as the most reliable and robustly conserved plaque-associated program across both systems (Fig. 5K, Fig. S19A, B). GO enrichment of these shared Up-Progressive genes confirmed the similar core biological processes identified in the organoid model, such as immune and inflammatory response, calcium signaling, lipid metabolism, and protein catabolism, corroborating their concordance in human AD tissue at single-cell spatial resolution (Fig. 5L, Fig. S19C, D, Table S9).

To characterize the biological processes underlying this concordance, we examined the plaque-associated trajectories of gene sets spanning key processes implicated in oligodendrocyte pathology and myelin loss (Fig. 5M, Fig. S20). Genes involved in amyloid sensing and clearance, calcium transport and signaling, and immune and inflammatory response, including cytokine, interferon, and antigen processing genes, were consistently elevated with increasing plaque density in both models, pointing to a coordinated stress and immune activation response in AD oligodendrocytes. Concurrent elevation of lipid breakdown and lipid synthesis genes was also observed, consistent with simultaneous membrane lipid catabolism and compensatory biosynthesis. In contrast, heat shock protein and unfolded protein response (UPR) genes declined with increasing plaque density, suggesting that the proteostatic stress response is attenuated rather than amplified at high plaque load.

Examination of proteasome-related gene modules revealed that proteasomal subunit composition is altered in both models in response to plaque burden, though the specific patterns differ between systems (Fig. 5M, Fig. S20). In human AD tissue, 11S activator subunits were elevated at high plaque density, while 20S core and 19S regulatory cap subunits showed more variable trajectories; in the organoid model, the patterns were largely inverted or non-monotonic. The alteration of proteasomal subunit composition across both systems, along with induction of immune and inflammatory, calcium signaling, and lipid metabolic programs, points to a global stress response to plaque burden that is qualitatively conserved across model systems despite context-specific differences in individual gene trajectories.

These observations are consistent with a pathogenic environment in which MBP, an intrinsically disordered protein whose stability depends on membrane association, may become a preferential target for altered catabolic activity. This provides a mechanistic link between plaque-associated proteostatic dysregulation, and the transcript-protein disconnect underlying myelin deficiency in AD. To directly test whether altered proteasomal activity contributes to MBP protein deficiency in the AD model, we treated hCOL and hCOL^APP^ organoids with two well-characterized proteasome inhibitors, the reversible inhibitor bortezomib and the irreversible inhibitor lactacystin, both previously reported to modulate MBP protein expression levels in vitro (*36, 37*). Following four weeks of treatment beginning at two months of organoid development, we assessed MBP protein levels and myelin ultrastructure by immunofluorescence and electron microscopy, with OLIG2 used to track oligodendrocyte lineage integrity (Fig. S21A). Bortezomib treatment did not restore MBP protein levels in hCOL^APP^ organoids but was associated with a reduction in OLIG2 expression, indicating disruption of oligodendrocyte lineage integrity that precludes direct interpretation of MBP protein level changes (Fig. S21B, C). In contrast, lactacystin treatment significantly restored MBP protein levels to those comparable to isogenic controls and was accompanied by recovery of compact myelin ultrastructure, evidenced by the restoration of circular, concentrically lamellated myelin sheaths around axons by EM, without altering OLIG2 expression (Fig. S21B, C). Notably, myelinated axons in lactacystin-treated hCOL^APP^ organoids appeared smaller in caliber and more numerous compared to the isogenic control, potentially reflecting preferential remyelination of smaller-diameter axons, or treatment-related changes in axonal composition. Together, these findings support a model in which altered proteasome-mediated catabolic activity contributes to MBP protein loss in the AD environment, providing a mechanistic link between plaque-associated proteostatic dysregulation and the transcript-protein disconnect underlying myelin deficiency in AD.

In this study, we demonstrate that amyloid plaque burden is associated with a reproducible, density-dependent transcriptional reprogramming of oligodendrocytes, characterized by induction of immune activation, calcium signaling, lipid catabolism, and altered proteasomal subunit composition, that is conserved across a human iPSC-derived organoid model and post-mortem AD brain tissue. This program defines a cellular environment incompatible with MBP accumulation, providing a molecular framework for the transcript–protein disconnect and myelin protein insufficiency observed in the amyloid-rich AD brain.

The involvement of white matter degeneration in AD has been recognized for over a century; Alois Alzheimer himself originally described lipid granule accumulation in glia and myelin thinning along with plaques and tangles (*2*). However, for decades, these observations were largely overshadowed by a neuron-centric focus, with myelin loss often viewed as a secondary consequence of axonal degeneration. Our findings provide molecular evidence supporting the possibility that oligodendrocytes adopt distinct plaque-associated states in amyloid-rich tissue that can contribute to myelin vulnerability. Importantly, this model does not exclude axon-dependent mechanisms; rather, it suggests that oligodendrocyte-intrinsic stress programs and local plaque-associated cues can coexist with, and potentially amplify, myelin disruption in degenerative contexts.

The broader consequences of this oligodendrocyte dysfunction may extend beyond the white matter. Recent work demonstrates that focal white matter demyelination drives transient grey matter microgliosis and synapse loss through retrograde signaling from demyelinated axon to neuronal somata, and that failure of remyelination results in chronic grey matter neuroinflammation (*38*). In the AD context, where our data suggest that amyloid-driven proteostatic disruption impairs MBP accumulation and myelin maintenance, the inability of oligodendrocytes to sustain the myelin sheath may similarly trigger maladaptive white matter-grey matter crosstalk, potentially amplifying the neuroinflammatory milieu and contributing to grey matter pathology independently of direct plaque-neuron interactions.

Human brain organoids paired with single-cell and spatial transcriptomics have enabled mapping of human-specific cellular states inaccessible in rodent models (*12, 39–41*), and single-cell profiling has defined oligodendrocyte heterogeneity in demyelinating disorders such as multiple sclerosis (*42, 43*). The present work extends this framework to AD, defining a plaque-associated oligodendrocyte state distinguished by proteostatic remodeling and linking oligodendrocyte heterogeneity to amyloid pathology. Importantly, this organoid platform enables compact myelin formation in a human context, captures human-specific plaque responses, and allows spatial coupling of plaque burden with oligodendrocyte state changes.

Mechanistically, our data point to altered proteasomal degradation as a contributor to myelin loss, supported by the selective restoration of MBP protein and compact myelin ultrastructure by lactacystin, an irreversible 20S proteasome inhibitor in hCOL^APP^ organoids (Fig. S21). Although our data support a model in which proteostatic dysregulation contributes to myelin loss, direct causal linkage will require additional perturbation studies. MBP is an intrinsically disordered protein (IDP) that requires chaperoning and membrane association to maintain its structure (*44–46*). Because MBP depends on membrane association for stability and oligodendrocytes are lipid rich, metabolically demanding cells, they may be particularly vulnerable to plaque-induced stress (*47, 48*). In the context of AD, we propose that the plaque-associated environment imposes severe proteotoxic stress (Fig. S22). The observed alterations in proteasomal subunit composition and the downregulation of cytosolic chaperones suggest that oligodendrocytes shift toward protein catabolism and immune surveillance, thereby limiting the accumulation of structurally necessary myelin proteins (*37, 49*). This is consistent with the known lability of IDPs; when the folding capacity of the cell is exceeded, these proteins are preferentially targeted for degradation, often via ubiquitin-independent mechanisms involving the 20S proteasome core. (*50, 51*). This proteostatic imbalance is likely exacerbated by lipid remodeling and inflammatory signaling, imposing additional membrane and metabolic stress on the cell.

This proteostatic vulnerability is compounded by the intrinsic biology of oligodendrocyte renewal. OPCs differentiate into new oligodendrocytes at a constitutive and constant rate that is independent of myelin demand or loss (*52*). Rather than mounting an accelerated regenerative response to myelin damage, OPC differentiation instead declines with aging and inflammation, two features that are prominent in the AD brain. New oligodendrocytes entering the plaque-rich environment will therefore encounter the same proteostatic stress conditions as their predecessors, rendering constitutive turnover insufficient to compensate for the catabolic bottleneck identified in the present study.

The upstream trigger linking plaque burden to oligodendrocyte proteostatic failure remains to be directly demonstrated. Our spatial data show a strong correlation between plaque density and the induction of calcium transport and signaling genes, pointing to calcium dysregulation as one candidate mechanism (Fig. S22). Intracellular calcium influx is a known downstream consequence of amyloid exposure and a potent trigger for cytoskeletal disassembly, protease activation, mitochondrial dysfunction, and immune activation (*53–56*). We therefore hypothesize that local amyloid exposure may induce calcium-dependent destabilization of the myelin sheath, releasing MBP and membrane lipids into both intracellular and extracellular compartments (*57–61*). This breakdown would not only expose labile proteins to degradation but also generate a pool of free lipids available for extensive remodeling and conversion into bioactive mediators that drive lipid-associated immune signaling (*62, 63*). Direct demonstration of this calcium-dependent mechanism will require targeted perturbation experiments.

We acknowledge several limitations in our study. Organoids model a developing brain-like environment and lack full adult white matter architecture, vasculature, and adaptive immune components; these differences may affect baseline maturation and inflammatory signaling (*64, 65*). Human spatial analyses also show donor-to-donor variability, highlighting heterogeneity in disease stage and regional vulnerability. The pharmacological rescue observed with lactacystin provides initial functional support for the proteasomal mechanism, though dose-response characterization and additional perturbation experiments will be needed to fully establish causality. Finally, our mechanistic conclusions are derived primarily from transcriptomic and pathway-level inference, and motivate future experiments that directly measure proteostasis flux, myelin protein turnover, and lipid mediator dynamics in plaque-associated oligodendrocyte states.

Collectively, this study reframes myelin loss in AD not as a passive consequence of neurodegeneration but as an oligodendrocyte-intrinsic response to the pathogenic amyloid microenvironment, driven by plaque-associated proteostatic disruption. The hCOL^APP^ platform established here provides a tractable human AD 3D model system to investigate oligodendrocyte contributions to AD pathology and serves as a foundation for testing interventions targeting disrupted oligodendrocyte proteostasis. By defining the pathologically relevant biological pathways and demonstrating their conservation across model systems, this work nominates oligodendrocyte proteostasis as a candidate therapeutic target to enhance glial resilience and attenuate white matter degeneration in AD.

## Acknowledgments

This work is fully funded by Merck Sharp & Dohme LLC, a subsidiary of Merck & Co., Inc., Rahway, NJ, USA.

## Author contributions

Conceptualization: J.S., R.M., B.C.

Surgery: J.H

Electrophysiology analysis: F.W., E.S.

Paraffin embedding and Bond Staining and analyses: K.C., J.C., A.M., C.S.C.

scRNA-seq library generation and analysis: S.V., B.C., J.S

Conceptualization of human AD spatial transcriptomics: A.T.

Human and organoid spatial transcriptomics data generation and analysis: Y.C., S.H, M.D., N.J., B.M., R.R., M.X., J.S

Funding acquisition: V.P., I.K.

Writing: J.S., B.C. (with input from all authors)

## Data, code, and materials availability

Single-cell and spatial transcriptomic data will be available in the Gene Expression Omnibus under accession code (will be set).

## Supplementary Materials

### Materials and Methods

#### Ethical Compliance

This study complies with all relevant ethical regulations approved by Merck & Co., Inc., Rahway, NJ, USA (MRL). All experiments involving hiPSCs were approved by the MRL Stem Cell for Drug Discovery (SCDD) Committee. All animal experiments described in this study were approved by the Institutional Animal Care & Use Committee (IACUC) of MRL. Post-mortem human brain tissues assessed in this study were obtained from NIH NeuroBioBank in accordance with applicable regulations.

#### Animals

The NOD SCID mice were purchased from Charles River Laboratories. Animals were housed at room temperature (68–79°F) with 40–60% humidity.

#### Cuprizone-Induced Demyelination and Xenotransplantation

Adult NOD-SCID mice (age and sex matched; supplier) were subjected to cuprizone diet to create a focal demyelination paradigm prior to organoid transplantation. Mice were fed an Envigo diet containing 0.2% cuprizone (TD.140804, Envigo) mixed with red food dye ad libitum for 3 weeks. Control animals received isocaloric control diet (TD.00588, Envigo) for the same duration. Food intake and body weight were monitored twice weekly throughout the treatment period. All procedures were performed in accordance with institutional animal care and use committee (IACUC) guidelines and approved protocols.

At the end of the 3-week cuprizone feeding period, mice received subcutaneous dexamethasone (2.5 mg/kg) to minimize edema and were then anesthetized with isoflurane. Each animal was positioned in a stereotaxic frame, and a small scalp incision was made to expose the skull. A 2-mm diameter craniotomy was performed by drilling at the following coordinates relative to bregma: anteroposterior (AP) +1.0 mm; mediolateral (ML) +1.4 mm. The dura mater was carefully removed, and a cortical cavity was generated by aspiration using a blunt-tip needle connected to a vacuum line. This unilateral aspirative lesion was made in the anterior motor cortex by removing the cortical tissue overlying the corpus callosum. Intact hCO and hCOL organoids grown for 60 days (size ∼1.6-1.8 mm) embedded with Matrigel as shown previously were then placed into the cavity (*1*). The organoid was covered and stabilized in situ and the craniotomy was sealed with Kwik-Sil silicone adhesive (World Precision Instruments). The incision was closed with sutures, and mice were allowed to recover in a temperature-controlled cage before being returned to their home cages. Animals were returned to control diet for the planned post-transplant survival period. At the indicated endpoints, mice were euthanized and transcardially perfused with PBS followed by 4% PFA. Brains were post-fixed overnight at 4°C and sectioned for immunohistochemistry. Engraftment and remyelination were assessed by immunostaining for MBP and human-specific OLIG2.

#### hiPSCs Culture

XCL-1, XCL-1_APP Ind (SwHomo)-3G05-E01, and BC1 hiPSCs were cultured on Matrigel (BD Biosciences) coated cell culture dishes with mTeSR1 Plus media (Stem Cell Technologies). hiPSCs were passaged every week by treatment with Dispase (0.83 U/ml, Stem Cell Technologies).

#### Generation of BC1 hiPSCs

As previously detailed (*1*), we constructed a cassette containing the doxycycline-inducible *SOX10*-P2A-*OLIG2*-T2A-*NKX6.2*-IRES-RFP and rTTA, and integrated it into the *AAVS1* locus of XCL-1. Briefly, 2 million XCL-1 cells were electroporated with 8 µg donor plasmid, 1 µg AAVS1 TALEN-L, and 1 µg AAVS1 TALEN-R using the Amaxa Nucleofector device (AAB-1001, Lonza) and then cultured in mTeSR1 Plus media supplemented with Y27632 (10 µM). G-418 (Thermo Fisher Scientific) was administered for 7 days (400 µg/ml for the first 3 days and 300 µg/ml for the following 4 days) to establish stable colonies. A single isogenic colony was selected and expanded for quality control analysis.

#### Generation of Ventral and Dorsal Forebrain Organoid Variants

To produce forebrain organoids with oligodendrocytes (hCO and hMGEO), a lentivirus containing the three transcription-factors (SOX10, OLIG2, NXK6.2; hereafter ‘3-TF’) (purchased from VectorBuilder) was first generated and transduced for 7 days without dox. As described earlier (*1, 2*), we generated hCOs and hMGEOs by combining 20% 3-TF-infected BC1-iPSCs with 80% non-infected parental XCL-1 hiPSCs. Briefly, after dissociating cells via Accutase, a total of 9,000 cells comprising 1,800 3-TF-infected and 7,200 XCL-1 hiPSCs were plated into a well of a U-bottom ultra-low-attachment 96-well plate in neural induction medium (DMEM-F12, 15% (v/v) KSR, 5% (v/v) heat-inactivated Fetal Bovine Serum (FBS) (Life Technologies),1% (v/v) Glutamax, 1% (v/v) MEM-NEAA, 100 µM β-Mercaptoethanol) supplemented with 10 µM SB-431542, 100 nM LDN-193189, 2 µM XAV-939, and 50 µM Y27632).

Basal activation of 3-TF was initiated on day 2 by adding 0.5 µM dox, and FBS and Y27632 were removed from day 2 and 4, respectively. The medium was replenished every other day until day 10, when organoids were transferred to the ultra-low-attachment six-well plate.

- hCO (Dorsal): Cultured in spinning dorsal patterning medium (day 10 to day 18) without vitamin A (1:1 mixture of DMEM-F12 and Neurobasal media, 0.5% (v/v) N2 supplement, 1% (v/v) B27 supplement without vitamin A, 0.5% (v/v) MEM-NEAA, 1% (v/v) Glutamax, 50 µM β-Mercaptoethanol, 1% (v/v) Penicillin/Streptomycin, and 0.025% Insulin).
- hMGEO (Ventral): Cultured in ventral patterning media (day 10 to day 18) containing DMEM-F12, 0.15% (w/v) Dextrose, 100 μM β-Mercaptoethanol, 1% (v/v) N2 supplement, 2% B27 supplement without vitamin A, 100 ng/ml recombinant SHH, and 1 μM purmorphamine.

The patterning medium was replenished every other day until day 18. Typically, media was then switched to maturation media (hCO medium with vitamin A, composition as above but with B27 containing vitamin A) supplemented with 20 ng/ml BDNF and 200 µM ascorbic acid. Maturation medium was changed every 3 days. Activation of 3-TF was performed beginning on day 18 by adding 2 µM dox continuously.

#### Generation of AD Forebrain Variants

Dorsal and ventral forebrain organoids were developed using XCL-1_APP Ind (SwHomo)-3G05-E01 and 3-TF-expressing BC1-iPSCs. AD control variants utilized 100% XCL-1_APP^Ind(SwHomo)-3G05-E01^. AD variants with OL-lineage cells (hCOL^APP^, hMGEOL^APP^) were created by mixing 20% BC1-iPSCs (driving OL-lineage) and 80% non-infected XCL-1_APP ^Ind(SwHomo)-3G05-E01^, followed by dorsal or ventral patterning and maturation as described above.

#### Live Imaging

Live images were captured in hCOL, hMGEOL, hCOL^APP^, and hMGEOL^APP^ at day 50 to visualize the RFP-expressing cells. The Phenix microscope, equipped with a controlled cell chamber maintained 37 °C temperature and 5% CO2, was used to generate z-stack images. 3D reconstruction of images was attained by using Leica Las-X software.

#### RT-qPCR analysis

Whole organoids were homogenized and total RNA was extracted using the RNeasy Mini Kit (Qiagen) according to the manufacturer’s instructions. One microgram of purified RNA was reverse-transcribed to cDNA with the iScript Select cDNA Synthesis Kit (Bio-Rad). Gene expression was measured by quantitative PCR on a CFX96 Real-Time System (Bio-Rad) using SsoFast EvaGreen Supermix (Bio-Rad). Thermocycling parameters were an initial denaturation at 95°C for 15 min, followed by 40 cycles of 94°C for 10 s and 60°C for 45 s. Primer sequences are:

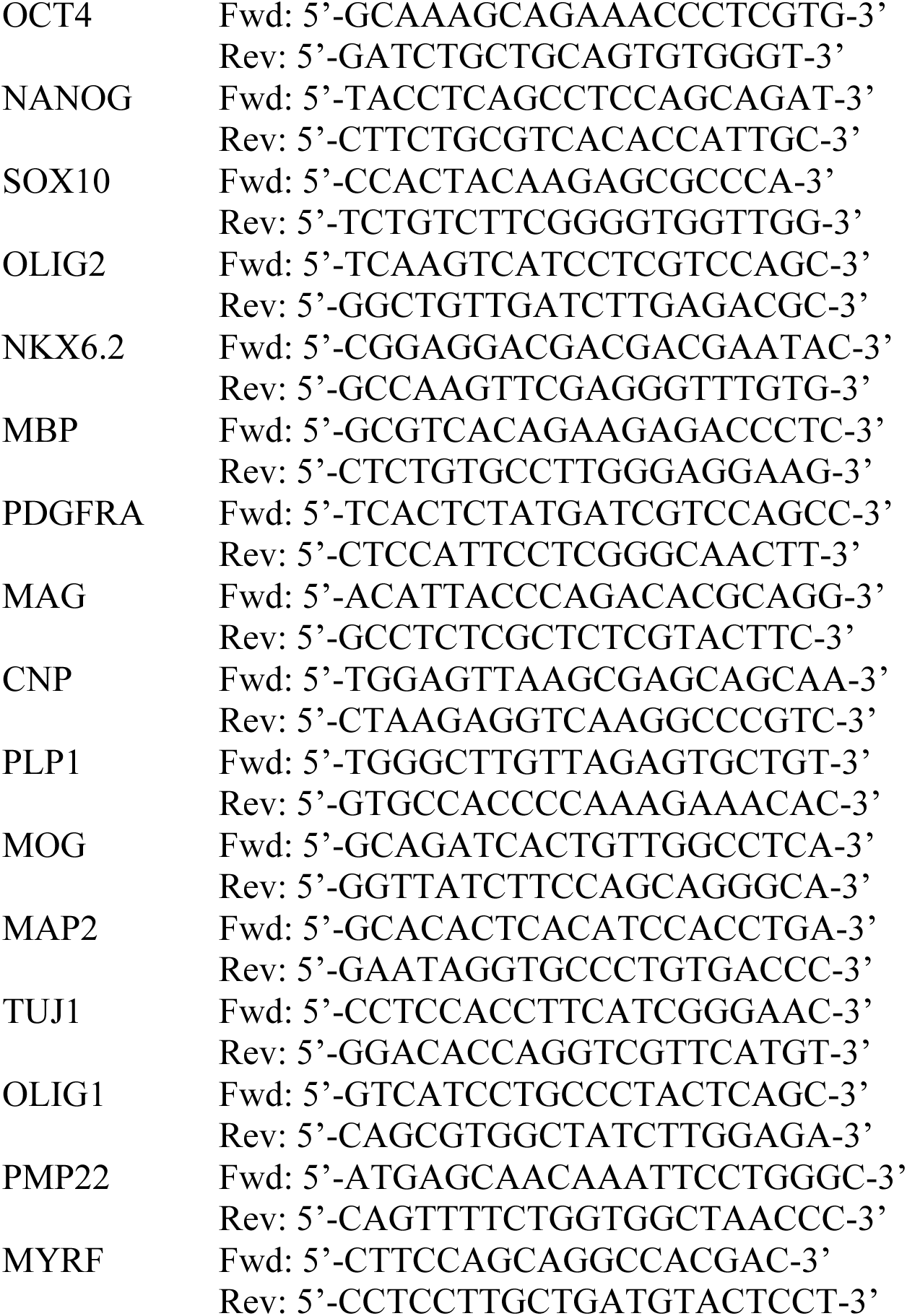

#### Electrophysiological Characterization of Oligodendrocyte-Enriched Brain Organoids

Multisite electrophysiological recordings were performed on intact human brain organoids using SomaFocus (Diagnostic Biochips Inc., Glen Burnie, MD), a high-throughput platform that automatically drives a 64-channel microfabricated silicon probe into a stationary organoid positioned at the bottom of a custom-designed well. Representative wideband traces from hCOL and hMGEOL recordings reveal spontaneous spiking activity across 16 channels spanning a 0.3 mm vertical depth, with both organoid types exhibiting diverse spiking patterns indicative of heterogeneous cellular composition and network activity (Fig. S1J).

#### Electron Microscopy

Transmission electron microscopy (TEM) was carried out in the Electron Microscopy Core Facility at UTSW. The forebrain organoid variants, hCOL, hMGEOL, hCOL^APP^, and hMGEOL^APP^ at day 90 and day 120, were fixed using 2.5% glutaraldehyde and 4% PFA in 0.1 M sodium cacodylate buffer for 1 hour at room temperature, followed by shipping overnight at 4 °C. After five rinses in 0.1 M sodium cacodylate buffer, they were post-fixed in 1% osmium tetroxide with 1.5 % K_3_[Fe(CN)_6_] in 0.1 M sodium cacodylate buffer for 1 h at room temperature. Cells were rinsed with water and en bloc stained with 0.5% aqueous uranyl acetate in 25% methanol overnight at 4 °C. After five rinses with water, specimens were stained with 0.02 M lead nitrate in 0.03M L-aspartate for 30 minutes at 60 °C. Samples were dehydrated with increasing concentration of ethanol, followed by propylene oxide, and infiltrated with Embed-812 resin and polymerized in a 60 °C oven overnight. Blocks were sectioned with a diamond knife (Diatome) on a Leica Ultracut UCT (7) ultramicrotome (Leica Microsystems) and collected onto copper grids. Images were acquired on a JEOL JEM-1400 Plus transmission electron microscope equipped with a LaB6 source operated at 120 kV using an AMT-BioSprint 16M CCD camera. To measure the percentage of morphological changes in myelinated axons in the organoid variants, approximately 100 axons per organoid were assessed. Morphological characteristics were manually identified, and NIH ImageJ was utilized to measure the relevant dimensions.

#### Automated multiplex immunofluorescence staining

Organoids were fixed, processed, and stained using an automated multiplex immunofluorescence pipeline. Whole organoids were collected into 6-well plates and fixed in 4% paraformaldehyde (PFA) at 4°C for 24 hours. After fixation, individual organoids were placed into separate tissue-processing cassettes and embedded in paraffin using a Leica HistoCore PEGASUS tissue processor (Leica Biosystems). To position organoids in the embedding mold, the warmed paraffin was manipulated with the tip of a tube and organoids were transferred into molten wax via Leica HistoCore Arcadia (Leica Biosystems). Paraffin blocks were configured to include three organoids per experimental condition alongside Alzheimer’s disease patient brain tissue controls. Blocks were sectioned at 5 µm on a Leica rotary microtome (Leica RM2235) onto charged glass slides to yield serial sections representing nine organoids across three conditions.

For immunofluorescence, slides were baked at 60°C for 1 hour, deparaffinized using the BOND Dewax protocol, and subjected to heat-induced epitope retrieval with BOND Epitope Retrieval Solution 2 for 20 minutes. Multiplex staining was performed on a Leica BOND RX autostainer using ready-to-use primary antibodies (mouse or rabbit monoclonals). An Opal-style sequential chromogenic/fluorophore amplification workflow was executed on-board, employing enzymatic amplification with BOND Polymer Refine Detection (HRP) and BOND Polymer Refine Red Detection (AP) and automated on-instrument mixing of chromogens. After staining, slides were scanned at 20X to 40X magnification on a Vectra Polaris and multispectral images were unmixed for downstream analysis via HALO (Indica Labs, v3.6) image analysis platform. High-plex immunofluorescence was analyzed with the Highplex FL module (v4.2.14) and corresponding marker-specific thresholds were applied uniformly across samples. Tissue coverage and signal intensity were quantified using the Area Quantification FL module (v2.3.4). Primary antibody incubations and dilutions used in this study were: anti-AT8 (Invitrogen, MN1020, 1:100), anti-MAP2 (Millipore Sigma, MAB3418, 1:100), anti-6E10 (Biolegend, 803002, 1:100), anti-hNuc (Millipore Sigma, MAB1281, 1:200), anti-OLIG2 (Cell Signaling, 65915S, 1:200), and anti-MBP (Cell Signaling, 78896S, 1:1000).

#### Luxol Fast Blue (LFB) staining

LFB staining for sectioned organoids was performed using the Newcomer Supply LFB–Cresyl Violet kit (Part 9155) following validated workflow. Paraffin sections (5 µm) were baked, deparaffinized in xylene, and rehydrated through graded alcohols to 95% ethanol. Sections were stained in Luxol Fast Blue solution (0.1% alcoholic) using a microwave-accelerated protocol (10 min at ∼70°C) and then rapidly rinsed in 95% ethanol and distilled water. Differential differentiation was achieved by brief dips (10–15 s) in 0.05% lithium carbonate followed by controlled differentiation in 70% ethanol until white and gray matter contrasted; slides were checked microscopically and further differentiated if needed. Sections were then counterstained in heated Cresyl Violet working solution (6 min at 57°C), rinsed, dehydrated through 95% and 100% ethanol, cleared in xylene, and coverslipped with mounting medium. All solutions and modifications follow the kit instructions; differentiation steps were performed carefully to avoid over-differentiation and to preserve myelin (blue) versus Nissl (purple) contrast. Slides were scanned at 40X magnification on a Vectra Polaris and multispectral images were unmixed for downstream analysis via HALO image analysis platform.

#### Proteasome Inhibitor Treatment

To assess the contribution of proteasomal activity to MBP protein deficiency in the hCOL^APP^ organoid model, hCOL and hCOL^APP^ organoids were prepared under standard aforementioned protocol until day 60, at which point mature myelinating oligodendrocytes are well established. Beginning at day 60, organoids were treated with either vehicle (0.1% DMSO), 5 nM bortezomib (Selleckchem, S1013), or 50 nM lactacystin (Cayman, 70980) for 30 days (n=3 organoids per condition). Treatment media was refreshed every 3-4 days. Organoids were harvested at day 90 for downstream analysis. MBP protein levels and oligodendrocyte lineage integrity were assessed by immunofluorescence staining for MBP and OLIG2, and myelin ultrastructure was evaluated by electron microscopy as described above.

#### Library Preparation of scRNA-seq

Each forebrain organoid variant at 50- and 115-days old was randomly collected from 3 different culture dishes, with 6-10 organoids pooled together. As previously described, the organoids were initially dissociated using the papain dissociation system according to the manufacturer’s instructions. Subsequently, after washing with HBSS, the organoids were dissected into small pieces in papain solution and oxygenated with 95% O2:5% CO2 for 5 minutes, and then incubated at 37 °C for 1 hour. Following the generation of a single-cell suspension via trituration, the single cells were suspended in 1% BSA/PBS supplemented with 10 μM Y27632 and stained for Aqua Dead Cell stain (Cat.no. 50-112-1525). FACS sorted live cells or RFP^+^ live cells were re-suspended in 0.04% BSA/PBS (128 cell/μl) and used to generate cDNA libraries by utilizing the Single Cell 3’ Reagent Kits. In brief, the cells were partitioned into nanoliter-scale Gel Bead-In-Emulsions (GEMs), and microfluidic cells were streamed at limiting dilution into a stream of Single Cell 3’ Gel Beads and then a stream of oil. After cell lysis, primers, an Illumina P7 and R2 sequence, a 14 bp 10xBarcode, a 10 bp randomer, and a poly-dT primer were released and mixed with the cell lysate and a bead-derived Master mix. Full-length cDNA from poly-adenylated mRNA was generated in each individual bead. Subsequently, after individual droplets were broken down, homogenized, and cleaned from the remaining non-cDNA, the libraries were size-selected, followed by the addition of R2, P5, P7 sequences to each selected cDNA. Finally, each library was sequenced using the Illumina HiSeq4000 2×150 bp in Rapid Run Mode.

#### scRNA-seq Data Analysis

Single-cell RNA-seq libraries were processed using Cell Ranger (v3.0.2). Reads were aligned to the hg19 human reference and quantified against Ensembl gene annotations using the standard Cell Ranger processing workflow with default parameters. Libraries were then depth-normalized and combined using the Cell Ranger aggregation workflow with default parameters. Prior to downstream single-cell analysis, doublet frequency was estimated by quantifying the fraction of cells co-expressing TBR1 and GFAP, which are typically restricted to cortical neurons and astrocytes, respectively; doublet frequency was low (0.82 ± 0.28%).

#### Per-sample quality control (five-step filtering)

After generating per-sample Seurat objects, QC was performed per sample using five sequential filters based on robust, distribution-derived thresholds. (1) Sequencing-depth filter: cells were retained only if total UMI counts (nCount_RNA) fell within the 10th–90th percentiles of the per-sample distribution. (2) Mitochondrial outlier filter: mitochondrial fraction (percent of counts mapping to genes matching MT-) was required to be below a robust cutoff defined as median(percent.mt) + 3×MAD(percent.mt) within each sample. (3) Ribosomal outlier filter: ribosomal fraction (percent of counts mapping to genes matching RPS and RPL) was required to be below the 99th percentile of the per-sample distribution. (4) Depth-adjusted complexity filter: to remove cells with low gene detection relative to sequencing depth, we fit a per-sample linear model log10(nFeature_RNA + 1) ∼ log10(nCount_RNA + 1) and excluded cells with standardized residuals ≤ −2. (5) Doublet filter: putative doublets were identified per sample using the scDblFinder workflow and excluded by retaining only cells classified as singlets.

#### Integration, dimensionality reduction, and clustering

QC-passed singlets from all samples were merged into a single Seurat object while preserving sample identity. The merged dataset was log-normalized, highly variable genes were selected, and scaled expression values were used for PCA. To mitigate sample- and batch-associated technical variation, integration was performed using the Harmony workflow with sample identity (and/or batch) as the grouping variable. Downstream neighborhood graph construction, UMAP embedding, and graph-based clustering were performed in the Harmony latent space using a dataset-appropriate number of components (typically ∼15–30 Harmony dimensions, selected based on standard variance/separation diagnostics and cluster stability). Clustering resolution was tuned per dataset to capture biologically meaningful granularity (typically ∼0.4–0.8 for parental datasets, and ∼0.5-1 for subclustering analysis), and final settings were chosen to balance separation of major lineages with avoidance of over-partitioning.

#### Cell Type Annotation

Cell-type identities were assigned at the cluster level using a modular, marker- and Gene Ontology (GO)–guided framework applied in a dataset-adaptive sequence to accommodate differences in lineage representation and state composition across organoid preparations. Cluster-wise average expression was computed from the RNA assay (Seurat AverageExpression). Marker-based assignments were made using predefined cluster-average expression thresholds specified in the analysis scripts for each marker module. GO-guided assignments were performed by testing significant cluster marker genes (adjusted P < 0.05) for over-representation of GO-defined gene sets using a one-sided Fisher’s exact test (alternative = “greater”), followed by Benjamini–Hochberg correction across clusters; GO calls used the stated FDR thresholds. Major lineages were resolved using canonical marker modules, including neuronal programs (e.g., STMN2/GAP43/DCX, with progenitor-like contamination excluded using VIM/HES1), inhibitory neuronal identity (GAD1/GAD2), and excitatory/cortical neuronal identity (e.g., SLC17A6). Progenitor/glial modules were detected using VIM/HES1 and further partitioned into cycling versus non-cycling states by enrichment for mitotic cell-cycle programs (GO:0000278; FDR < 0.05). Oligodendrocyte lineage states were identified using OPC markers (OLIG2/SOX10/PDGFRA) and GO enrichment for oligodendrocyte differentiation (GO:0048709; FDR < 0.01), whereas astrocytic identity was assigned using GFAP/AQP4. Mesenchymal/proteoglycan-expressing clusters were identified using extracellular matrix–associated markers (e.g., BGN/DCN) and, where relevant, mesenchymal-like signatures (e.g., COL1A1/FOXC1). Additional cell clusters were annotated using GO enrichment for cilium-associated genes (GO:0060271; FDR < 0.05) and unfolded protein response genes (GO:0006986; FDR < 0.05). Any residual clusters not meeting criteria for these modules were retained as unspecified. Final labels were stored as metadata and used for visualization and compositional analyses.

#### Subtype Analysis

To resolve finer cellular states within major organoid lineages, we performed lineage-restricted subtyping by subsetting the integrated Seurat object to a lineage of interest (e.g., neuronal, astrocytic, oligodendrocytes) based on the cluster-level annotation metadata. For each lineage subset, the analysis was re-run in a within-lineage manner to improve separation of closely related states. Specifically, cells were subset, log-normalized, and variable features were identified before scaling and PCA. Where required to reduce condition-driven technical structure, subsets were integrated using the Harmony workflow with the relevant grouping as the integration covariate. Neighbor graph construction, UMAP embedding, and graph-based clustering were then performed in the Harmony latent space using a dataset-appropriate number of dimensions and clustering resolution chosen to capture stable within-lineage substructure while avoiding over-partitioning. Subtype identities were stored as a lineage-specific clustering and used for downstream marker discovery and compositional comparisons across conditions.

#### Pseudotime trajectory analysis

Pseudotime trajectory inference was performed on lineage-restricted subsets exhibiting continuous state progression. Trajectories were inferred using the Slingshot workflow applied to a reduced-dimensional embedding together with graph-based cluster labels. To orient trajectories in a biologically meaningful direction, we typically defined a starting (root) cluster representing the earliest/most progenitor-like state in the subset (e.g. OPC cluster) and provided this root to the trajectory inference workflow. Slingshot was then used to fit lineage curves through cluster centroids, generating per-cell pseudotime values for each inferred lineage. Pseudotime values were stored in cell metadata and summarized across clusters to order states and compare pseudotime distributions and state occupancy across experimental conditions.

#### Differential gene expression (DEG) analysis

Differential gene expression was performed using Seurat within lineage-restricted subsets and/or within defined subtypes to identify transcriptional changes associated with experimental condition. For each analysis, cells were subset based on metadata, and group comparisons were conducted either (i) within a given subtype/cluster or (ii) between subtypes within a given condition when required. Differential expression testing was performed using Seurat FindMarkers with the Wilcoxon rank-sum test (test.use = “wilcox”). Genes were required to be detected in at least 25% of cells in either comparison group (min.pct = 0.25), and ta minimum effect-size threshold was typically applied (logfc.threshold = 0.25). P values were adjusted for multiple testing using the Benjamini–Hochberg method, and genes were designated as differentially expressed based on adjusted P value (FDR) < 0.05. DEGs were reported with estimated log fold-change and adjusted P value, and were used as inputs for downstream functional enrichment analyses (e.g., GO Biological Process over-representation) and visualization (e.g., volcano plots, heatmaps, and dot plots) as appropriate.

#### Gene Ontology (GO) Pathway Enrichment analysis

GO pathway enrichment analyses were performed to interpret biological programs associated with condition- and subtype-associated transcriptional changes. Significant gene sets (e.g., DEGs passing adjusted P value/FDR < 0.05) were used as input for GO over-representation analysis and, where appropriate, were stratified by direction of effect based on the sign of the estimated log fold-change. For GO enrichment of plaque density-dependent pattern gene sets, the background gene set differed between model systems: for organoid analyses, the entire human genome was used as background; for human AD spatial transcriptomics analyses, all genes detected in the dataset were used as background. Enrichment was computed with clusterProfiler (enrichGO) using the Biological Process ontology (ont = “BP”), with human gene symbols provided as input (keyType = “SYMBOL”) and annotation against org.Hs.eg.db. Enrichment P values were adjusted using the Benjamini–Hochberg method (pAdjustMethod = “BH”), and GO terms were considered enriched typically using pvalueCutoff = 0.05 and qvalueCutoff = 0.2. Enriched terms were summarized and visualized (e.g., donut charts or dot plots of top categories) to contextualize lineage- and state-specific programs.

#### Human metadata integrated analysis

For human AD scRNA-seq datasets, we obtained per-sample GEO (Gene Expression Omnibus; https://www.ncbi.nlm.nih.gov/geo/) GSM accession IDs and used these identifiers to link cells to sample-level metadata (e.g., control vs AD and other available covariates). Count matrices were imported into R to generate Seurat objects, and each dataset was processed using the same QC filtering, normalization, integration, dimensionality reduction, and clustering workflows described above. The human datasets analyzed included:

Shun-Fat L et al. (GSM4775580, GSM4775578, GSM4775577, GSM4775576, GSM4775572, GSM4775569, GSM4775567, GSM4775561),

Soreq L et al. (GSM5348374, GSM5348375, GSM5348376, GSM5348377),

Xu J et al. (GSM4888887, GSM4888888, GSM4888889, GSM4888890, GSM4888891, GSM4888892, GSM4888893, GSM4888894),

Ramos S.I. et al. (GSM6720852, GSM6720854, GSM6720856, GSM6720858, GSM6720860, GSM6720862, GSM6720864, GSM6720866, GSM6720868, GSM6720870, GSM6720872, GSM6720874, GSM6720876, GSM6720878, GSM6720880, GSM6720882, GSM6720884, GSM6720886).

Cell types were assigned using canonical marker genes, and integrated objects were subset to the oligodendrocyte lineage (OPC/oligodendrocyte populations) for downstream analyses.

After subsetting each dataset to the oligodendrocyte lineage (OPCs/oligodendrocytes), oligodendrocyte-lineage cells from brain organoids (hCOL and hCOL^APP^) and human datasets were integrated into a shared space using Seurat’s anchor-based integration framework. Briefly, oligodendrocyte-lineage subsets from each source were treated as separate integration inputs and processed in parallel with log-normalization and variable-feature selection. A common set of integration features was selected (SelectIntegrationFeatures), anchors between datasets were identified (FindIntegrationAnchors), and datasets were merged into an integrated expression space (IntegrateData) to reduce study-specific technical differences while preserving shared biological structure. The integrated object was subsequently used for downstream dimensionality reduction, neighborhood graph construction, UMAP visualization, clustering, marker discovery, subtype definition, differential expression, and enrichment analyses using the procedures described above.

#### Plaque density and proximity tertile definition

Plaque density values and nearest-plaque distances, computed as described in Plaque Density and Distance Measurement, were discretized into tertiles to enable stratified analyses. Tertile thresholds were defined using the full set of cells in the spatial transcriptomic dataset (all cell types), rather than the oligodendrocyte-lineage subset, to avoid lineage-restricted distributions biasing the binning. Plaque density was divided into three groups using the 33rd and 67th percentiles of its distribution across all cells: cells at or below the 33rd percentile were classified as low density, those between the 33rd and 67th percentiles as mid density, and those above the 67th percentile as high density. Nearest-plaque distance was similarly divided into tertiles: cells at or below the 33rd percentile were classified as proximal, those between the 33rd and 67th percentiles as intermediate, and those above the 67th percentile as distal. Tertile assignments were stored as metadata and used for downstream comparisons within oligodendrocyte-lineage cells. Per-sample visualization confirmed that density-expression trends were directionally consistent across individual samples (Fig. S14, Fig. S18), supporting the use of globally defined tertile boundaries.

#### Plaque-density–dependent expression patterns and response class assignment

Plaque density tertile labels (Low, L; Mid, M; High, H), defined on the full spatial dataset as described in Plaque density and proximity tertile definition, were retained when subsetting to oligodendrocyte-lineage cells for downstream analysis. For each condition, we computed the mean expression of each gene within each density bin in the oligodendrocyte-lineage subset. To compare trends across bins independently of baseline expression, we converted each gene’s three-bin mean profile to a row-wise z-score: for each gene, the mean of the three bin means was subtracted and the result divided by their standard deviation, yielding a standardized three-point trajectory z(L), z(M), z(H) per gene. No minimum detection filter was applied prior to pattern classification; genes with near-zero or stochastic expression across all tertiles were effectively excluded by the flat gene filter described below.

Genes with insufficient dynamic range were excluded as flat: a gene was classified as flat if all three z-scores had absolute value ≤ 0.30. This threshold was chosen as a conservative criterion to exclude genes with minimal variation across tertiles, as values within 0.30 standard deviations of zero represent negligible expression change relative to the dynamic range captured by the z-score normalization. For non-flat genes, overall direction was determined by the sign of z(M)−z(L): genes with z(M)−z(L) > 0 entered the upward branch; those with z(M)−z(L) < 0 entered the downward branch. Within each branch, the sign and magnitude of z(H)−z(M) determined the final pattern assignment: z(H)−z(M) > 0 yielded Progressive; |z(H)−z(M)| < 0.30 yielded Plateau; z(H)−z(M) < 0 yielded the reversal pattern (Up-Down or Down-Up). This yields six mutually exclusive pattern classes: Up-Progressive, Up-Plateau, Up-Down, Down-Progressive, Down-Plateau, and Down-Up. Pattern labels were computed separately within each condition and used for downstream visualization and functional interpretation.

Pattern assignments are deterministic and based solely on tertile means, without a formal statistical test of whether differences between tertile means exceed noise. For genes with high cell-to-cell variability or low cell counts per tertile, mean estimates may be imprecise and assignments near classification boundaries should be interpreted with caution. Cross-model conservation of pattern assignments between organoid and human AD tissue (Fig. 5K, Fig. S19) provides empirical support for the reliability of the classification, as independent biological systems are unlikely to consistently reproduce noise-driven assignments.

#### Spatial Transcriptomics (Organoid)

To generate the single-cell spatial omics dataset for CRISPR-edited brain organoids, organoid samples harvested at different time points were embedded to formalin-fixed paraffin block and sectioned to 5 μm serial slices using a microtome. Sample processing, staining, and imaging followed established protocols. In brief, tissue sections were mounted on VWR Superfrost Plus Micro Slides (Cat# 48311-703) to ensure optimal adherence, air-dried at 37°C for 16 hours, and subjected to deparaffinization with xylene (2 × 5 min, for RNA assay) or CitriSolv (for protein assay) followed by ethanol rehydration. Heat-induced antigen retrieval was performed in citrate buffer (pH 6.0) at 95°C for 8 min, followed by proteinase K-mediated permeabilization (1 μg/mL, 37°C, 15 min, only for RNA assay) as adapted from the CPA-processing procedure in MAN-10184-05 (https://university.nanostring.com/cosmx-smi-manual-slide-prep-for-rna-assays).

For RNA readout, RNA in situ hybridization (RNA-ISH) probes (1 nM), designed to detect 6175 protein-encoding mRNAs (with ∼4000 brain specific targets), were hybridized at 37°C for 18 hours in a humidified chamber. Post hybridization, slides underwent stringent washing (2× SSC with 50% formamide, 37°C, 2 × 25 min) to remove unbound probes. A flow cell was assembled atop each slide, and cyclic RNA readout was conducted using the CosMx Spatial Molecular Imager (SMI) with 108-bit combinatorial encoding strategy (27 imaging rounds, 4 color channels per round) for transcript identification.

Prior to RNA detection cycles, morphology markers were applied for visualization and cell segmentation: GFAP antibody (Alexa Fluor-647) for glial staining, rRNA oligo probes (Dyomics Dy-605) for cytoplasmic staining, Histone H3 antibody (Alexa Fluor-488) and DAPI for enhanced nuclear staining. 10-25 fields of view (FOVs, 0.509 mm × 0.509 mm) per organoid were placed to cover all intact specimens on the slide, determined via pre-scan imaging. Transcript data collection was performed using the commercial CosMx platform, an epifluorescence microscope with a customized water-immersion objective (magnification 22.7×, NA 1.1). Widefield illumination was provided by a combination of lasers and light-emitting diodes (385 nm, 750 mW; 488 nm, 1 W; 530 nm, 1 W; 590 nm, 150 mW; 647 nm, 1 W), enabling excitation of DAPI, Alexa Fluor-488, Atto-532, Dyomics Dy-605, and Alexa Fluor-647, while photocleavable dye components were cleaved post-imaging. Images were captured with a Lucid Atlas10 ATX sCMOS sensor (pixel size 120 nm, 10 MP resolution). For each FOV, a quasi-3D multichannel image stack (8 z-planes, 0.8 μm step size) was acquired per imaging round, totaling 27 rounds to decode the 6k-discovery panel.

For protein readout, the tissue section was first blocked with CosMx reagent buffer-W, followed by overnight antibody mix incubation. Brain specific markers were applied for visualization and cell segmentation: Gfap antibody (Alexa Fluor-488) for glial staining, NeuN (Atto-532) and Map2 (Dyomics Dy-605) antibodies for neuron staining, Iba1 antibody (Alexa-647) for microglia staining, and DAPI for nuclear staining. The protein targets were sequentially imaged on CosMx (16 imaging rounds) without combinatorial barcoding.

Image processing involved registration, feature extraction, background subtraction, transcript localization, and decoding as previously described (*3*). For initial data quality control, machine learning-based cell segmentation, adapted from Cellpose (*4*), was applied using a pre-trained model fine-tuned on morphology marker channels (membrane plus nucleus). The final dataset mapped each decoded transcript or protein signals to individual cells and subcellular compartments within the registered image stack, enabling spatial analysis of multiple biomolecules at subcellular resolution.

#### Transfer of scRNA-seq cell-type labels to spatial transcriptomics

To annotate spatial transcriptomics profiles using scRNA-seq–defined cell types, we performed reference-to-query mapping in Seurat using an anchor-based label transfer workflow. Briefly, a reference Seurat object containing scRNA-seq cell-type annotations (metadata field annotation) and an integrated low-dimensional representation was used as the atlas, and a query Seurat object containing spatial transcriptomics expression profiles (spots) was used as the target for label projection.

The reference object was confirmed to contain a PCA embedding; if PCA was not present, the reference was preprocessed by log-normalization, variable feature selection, scaling, and PCA (30 components). To enable projection of query data into the reference manifold, a UMAP model was computed on the reference using the integrated latent space (UMAP run on the reference integration reduction; 20 dimensions) with return.model = TRUE, generating a reusable UMAP transformation for query projection.

Cross-modality correspondences between reference cells and query spots were then learned using Seurat’s transfer anchors. Specifically, anchors were computed with FindTransferAnchors using PCA projection (reduction = “pcaproject”) to map the query onto the reference low-dimensional space, while explicitly specifying the reference integrated reduction used for alignment (reference.reduction set to the reference integration reduction). Label transfer and projection were then performed with MapQuery, supplying the anchor set and using the reference annotation as the label source (refdata = list(annotation = “annotation”)). This procedure generated, for each spatial spot, a predicted cell-type label (stored as predicted.annotation) and associated prediction scores, and additionally projected the query into the reference UMAP space (stored as ref.umap) using the learned UMAP model. Predicted labels were subsequently visualized in the projected embedding and used for downstream spatial analyses. All downstream analyses of the mapped objects (visualization, subsetting, marker inspection, and group comparisons) followed the same general procedures described for scRNA-seq analyses above.

#### Plaque Density and Distance Measurement

To assess amyloid- β plaque density, the antibody ab-1-40 was applied to adjacent sections of hCOL/hMGEOL and hCOL/hMGEOL^APP^. Signal intensity was background-subtracted, and plaque puncta were inferred using an in-house deep-learning semantic segmentation model. To ensure data quality, masks representing puncta with an area smaller than 20 µm² were excluded from further analysis.

The remaining masks were registered to the RNA layer using an in-house deep-learning image registration algorithm that includes the conventional affine transformation as well as a nonlinear elastic transformation that locally fine-tunes the alignment between the two images. From these registered masks, two types of spatial transforms were computed:

1. <u>Nearest-plaque distance</u>: This was calculated as the standard Euclidean distance transform of the plaque binary mask, scaled by the physical pixel size to yield physical distance in micrometers. This metric represents the shortest distance from the background (e.g. a cell centroid) to the nearest plaque.
2. <u>Plaque Density</u>: To capture the cumulative local plaque burden and to account for plaque size—factors not reflected in the standard distance transform—we developed an original, physically motivated, in-house algorithm to compute a more robust density metric. The method convolves the binary plaque mask with a kernel corresponding to the fundamental solution of the heat equation. This approach can be conceptually understood in two ways: as modeling the diffusion of heat from concentrated heat sources placed at each plaque pixel, or as a scaled kernel density estimation of the plaque pixels distribution. It provides a continuous density value (in µm^-2^) at each pixel that reflects the influence of all nearby plaques, (nonlinearly) weighted by their distance. The kernel parameters were scaled by the physical pixel size to ensure the analysis was invariant to image resolution. Implementation-wise, the fundamental solution to the heat equation is the normal distribution so Gaussian blur—a standard image processing operation—of the plaque mask was performed. For computational efficiency, spatial separability of the Gaussian kernel and convolution in the Fourier domain were exploited.

To associate plaque density or distance with individual cells, the centroids of the cells in the RNA image were utilized to extract the corresponding plaque density or distance values.

#### Human postmortem brain tissue acquisition and processing

Formalin fixed paraffin embedded (FFPE) human brain tissue blocks were obtained from the NIH NeuroBioBank under approved institutional agreements and in accordance with all ethical and regulatory requirements.

Thirteen donors were selected for this study, consisting of six neurotypical controls and seven individuals presenting with Alzheimer’s disease neuropathologic change (ADNC) according to NIA-AA criteria (*5*). All tissue blocks originated from Brodmann area 39 (BA39) within the temporal-parietal cortex. Donor demographic information and sample specific metadata are provided in Supplementary Table 5 (Table S5). To ensure adequate RNA preservation in archival FFPE material, all blocks underwent a standardized pre screening process. RNA integrities were assessed using both the RNAscope LS Multiplex Fluorescent Reagent Kit (ACD Bio #322800) and DV200 evaluation. Following quality assessment, each block was sectioned in an RNase free environment. The first five 5 µm sections from each block were discarded to eliminate tissue exposed to prolonged oxidation or surface degradation. Subsequent 5 µm serial sections were collected onto Leica Bond slides (S21.2113.A) and reserved for histology, and spatial transcriptomics profiling as detailed below. Slides were stored in RNase-free containers under controlled temperature and humidity to maintain RNA integrity prior to processing.

#### Histopathology

The first collected 5 µm section from each block was processed for standard H&E staining on the Leica Autostainer XL ST5010. Whole slide images were captured at 40x magnification on a Leica Aperio AT2 digital slide scanner. These images were evaluated by expert neuropathologists to assess overall tissue quality and annotate the cortical layers and white matter region. The second 5 µm serial section for each block was subjected to duplex immunohistochemistry on the Biocare ONCORE Pro X using 6E10 (BioLegend #803001) to identify amyloid-β plaques and AT8 (Thermo Fisher Scientific #MN1020) to detect Ser202/Thr205 phosphorylated tau (neurofibrillary tangles). Detection was achieved using enzyme conjugated polymer secondary systems (HRP for DAB and AP for RED chromogen development), followed by a hematoxylin counterstain for nuclear visualization. Imaging was performed on a Leica Aperio AT2 digital slide scanner at 40x resolution. Neuropathologists reviewed all stained sections and annotated regions containing amyloid-β plaques and neurofibrillary tangle–rich pathology.

#### Spatial Transcriptome (human postmortem brain tissue)

Spatial transcriptomic profiling was performed on the third collected 5 µm section for each block using the CosMx Spatial Molecular Imager (Bruker) and the CosMx Human Whole Transcriptome (WTx) assay, enabling subcellular resolution detection of approximately 19,000 human genes. All steps followed the manufacturer’s protocol: CosMx SMI Manual Slide Preparation for RNA Assays (Bruker MAN 10184 06). In brief, slides were baked at 60 °C overnight and equilibrated to RT for 3–5 min prior to deparaffinization using xylene and ethanol washes. Slides were then air dried at 60 °C for 5 min and immersed in pre heated 1x Target Retrieval Solution, followed by pressure cooking at 100 °C for 15 min without pressure. Slides were transferred to water for 15 s, washed in 70% ethanol for 2 h, then washed in 100% ethanol for 3 min, and air dried at RT for 30–60 min. Incubation frames were applied, and slides were treated with pre warmed Proteinase K (3 µg/mL in 1x PBS + 0.5% Tween) for 30 min at 40 °C. After two water rinses, 0.0005% fiducials were added for 5 min at RT, protected from light. Slides were washed in 1x PBS (1 min), fixed in 10% NBF (1 min), washed twice in NBF stop buffer (5 min each), and rinsed in PBS (5 min). NHS acetate (100 mM) was applied for 15 min at RT, followed by two 5 min washes in 2x SSC. The CosMx Human WTx (19K-plex) RNA probe mix panel and rRNA segmentation marker was denatured (95 °C, 2 min), cooled on ice (1 min), mixed with RNase inhibitor, Buffer R, and nuclease free water, applied to slides, and hybridized for 18 h at 37 °C. Post hybridization, slides were washed twice in 50% formamide/2x SSC for 25 min each in a 37°C water bath, then twice in 2x SSC for 2 min. DAPI (1:40 in blocking buffer) was applied for 15 min at RT, followed by a 5 min PBS wash. Slides were incubated for 1 h at RT with histone H3, GFAP, segmentation markers, washed three times in PBS (5 min each), and stored in 2x SSC. Pre bleaching used configuration B; segmentation imaging used configuration B. High resolution imaging (tiling + Z stacks) was performed on the CosMx platform, and Bruker’s SMI pipeline was used for stitching, registration, transcript decoding, and initial cell segmentation.

#### Statistical Analysis

Statistical analyses were performed in R/Python for transcriptomic and computational analyses and in GraphPad Prism for biochemical/imaging quantifications. Unless otherwise indicated, data are presented as mean ± SD for normally distributed measurements and as median with interquartile range (IQR) for non-normally distributed measurements. Two-group comparisons were performed using two-sided Mann–Whitney U tests for nonparametric data and two-sided Welch’s t-tests when unequal variances were expected. For comparisons involving more than two groups, we used Kruskal–Wallis one-way ANOVA (nonparametric), with post hoc multiple-comparison testing in Prism where applicable. For distribution-level comparisons of continuous single-cell measures (e.g., pseudotime distributions), we used the two-sample Kolmogorov–Smirnov test; for comparisons of central tendency of single-cell measures between two groups, we used the Wilcoxon rank-sum test as indicated in figure legends. Correlation analyses were performed using Pearson correlation, and where nonlinear relationships were expected (e.g., density as a function of distance), trend lines were fit using a generalized additive model (GAM). Multiple testing for transcriptomic differential expression and functional enrichment analyses was controlled using Benjamini–Hochberg false discovery rate (FDR) correction, with adjusted P values reported. Exact sample sizes (n), the number of independent experiments or donors, and the specific statistical tests used for each panel are reported in the figure legends.

---

**Figure S1.**
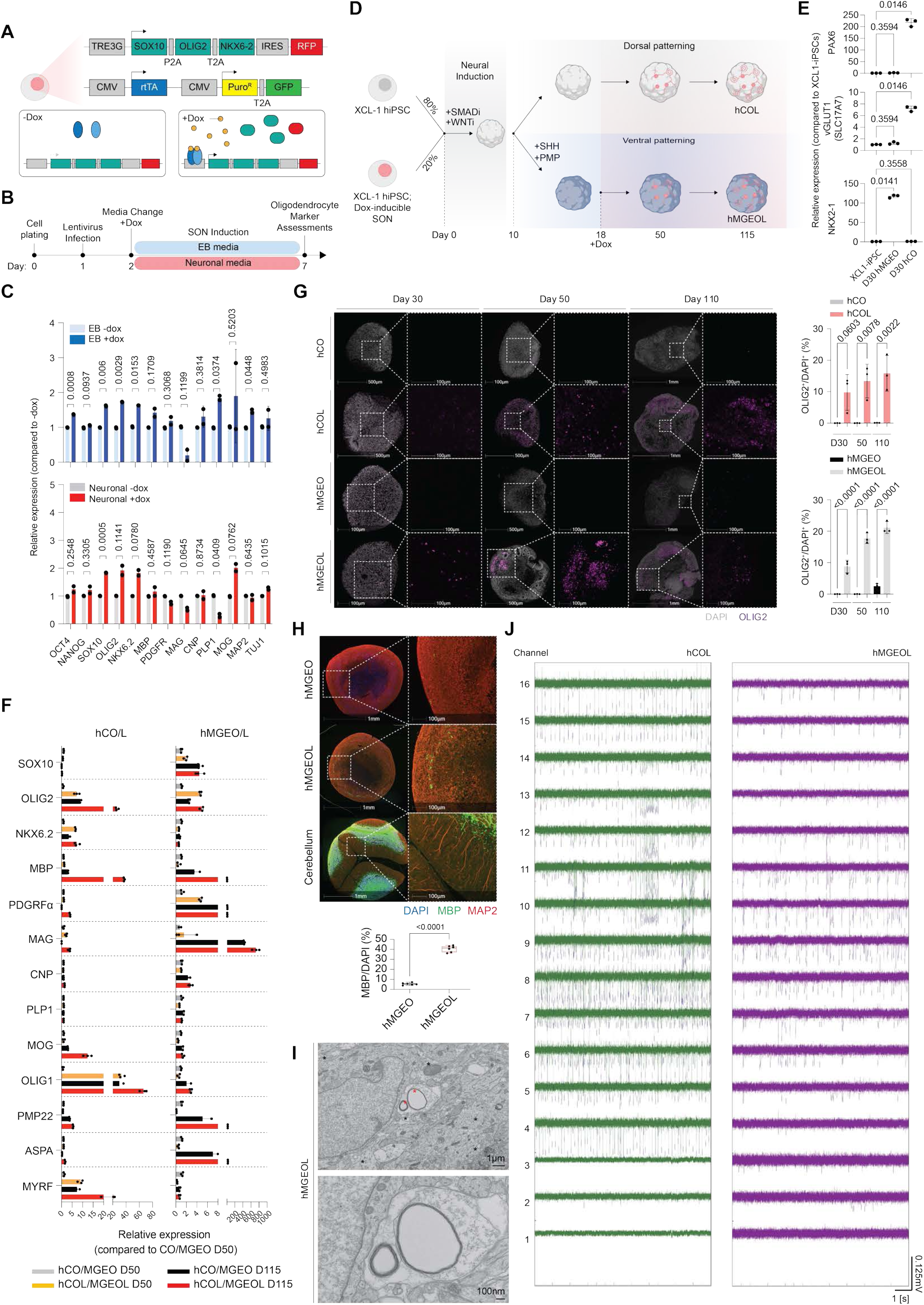
Generation and validation of oligodendrocyte-enriched dorsal and ventral forebrain organoids, hCOL and hMGEOL. (**A**) Schematic of the lentiviral vectors designed for the inducible expression of the transcription factors *SOX10*, *OLIG2*, and *NKX6.2* (SON factors) under the tetracycline-responsive element 3G (TRE3G) promoter, and the constitutive expression of reverse tetracycline-controlled transactivator (rTTA). (**B**) Experimental timeline for the validation of the SON system in iPSCs cultured in Neuron induction or EB induction media. (**C**) RT-qPCR quantification of pluripotency (*OCT4*, *NANOG*) and oligodendrocyte lineage markers, such as *SOX10*, *OLIG2*, *MBP*, in SON-infected XCL-iPSCs treated with or without doxycycline (dox, 5 µM). 2 iPSC lines from 3 differentiation experiments. Mean±SD. Welch’s t-test. (**D**) Schematic protocol for generating dorsal forebrain-patterned (hCOL) and ventral forebrain-patterned (hMGEOL) oligodendrocyte-enriched organoids. (**E**) RT-qPCR quantification of regional forebrain identity markers (*PAX6*, vGLUT1 (*SLC17A7*) for dorsal; *NKX2-1* for ventral) at day 30. Mean±SD. Šidák multiple comparison test. (**F**) RT-qPCR analysis of oligodendrocyte lineage markers in hCO/hCOL (left) and hMGEO/hMGEOL (right) at day 50 and day 115. Mean±SD. (**G**) (Left) Representative immunofluorescence images of hCO/hCOL and hMGEO/hMGEOL at days 30, 50, and 110 stained for OLIG2 (magenta). (Right) Quantification of the percentage of OLIG2+ cells normalized to DAPI. hCO, n=9; hCOL, n=9; hMGEO, n=8; hMGEOL, n=8; organoids from 2 iPSCs from 3 differentiation experiments. Mean±SD. Šidák multiple comparison test. (**H**) (Top) Immunofluorescence of hMGEO and hMGEOL stained for neurons (MAP2, red) and oligodendrocytes (MBP, green). Human cerebellum is included as a positive control. (Bottom) Quantification of MBP+ cells in hMGEO and hMGEOL. Mean±SD. Welch’s t-test. (**I**) Electron microscopy (EM) analysis of hMGEOL. (Top) Low-magnification (1 µm) view showing myelinated axons (red/black arrowhead). (Bottom) High-magnification (100 nm) view showing compact myelin ultrastructure. hMGEOL, n=12, from 2 iPSCs from 3 differentiation experiments. (**J**) Electrophysiology recording from the interior of intact oligodendrocyte-enriched brain organoids. Wideband, 10-second recording windows from (Left) hCOL and from (Right) hMGEOL showing spontaneous spiking activity across one probe shank with 0.33 mm depth coverage.

**Figure S2.**
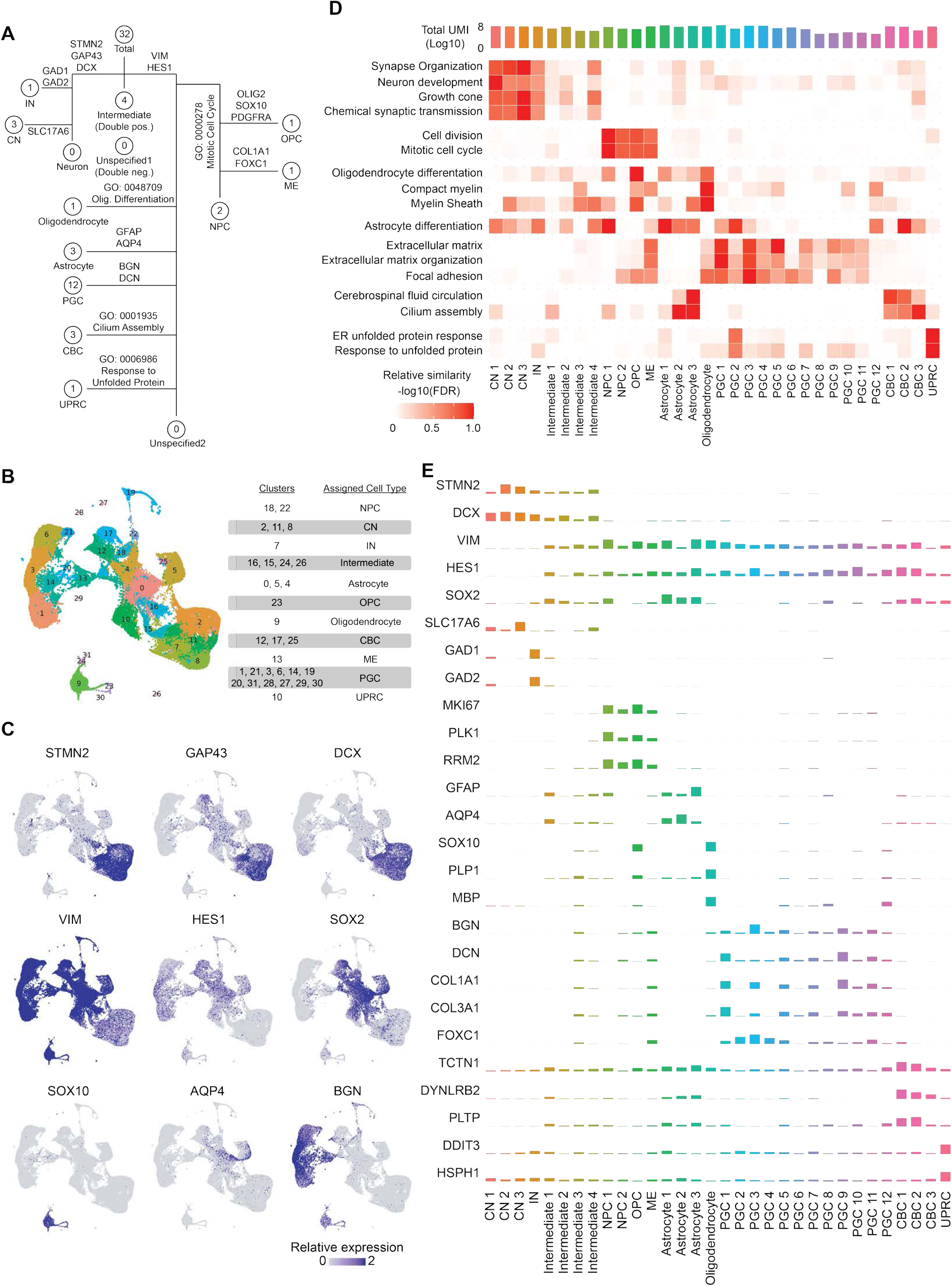
scRNA-seq characterization and cell type annotation of hCOL. (**A**) Hierarchical cell type annotation decision tree. The number within each circle represents the number of clusters at that annotation step. Branching decisions are made by filtering cell clusters based on specific marker genes or Gene Ontology (GO) terms. (**B**) UMAP visualization of the major cell populations identified in hCOL. (**C**) Feature plots showing the expression of key cell type markers. (**D**) Heatmap displaying the False Discovery Rate (FDR) for selected GO terms used to annotate the clusters. (**E**) Bar plots showing the expression levels of the indicated genes for each cell type.

**Figure S3.**
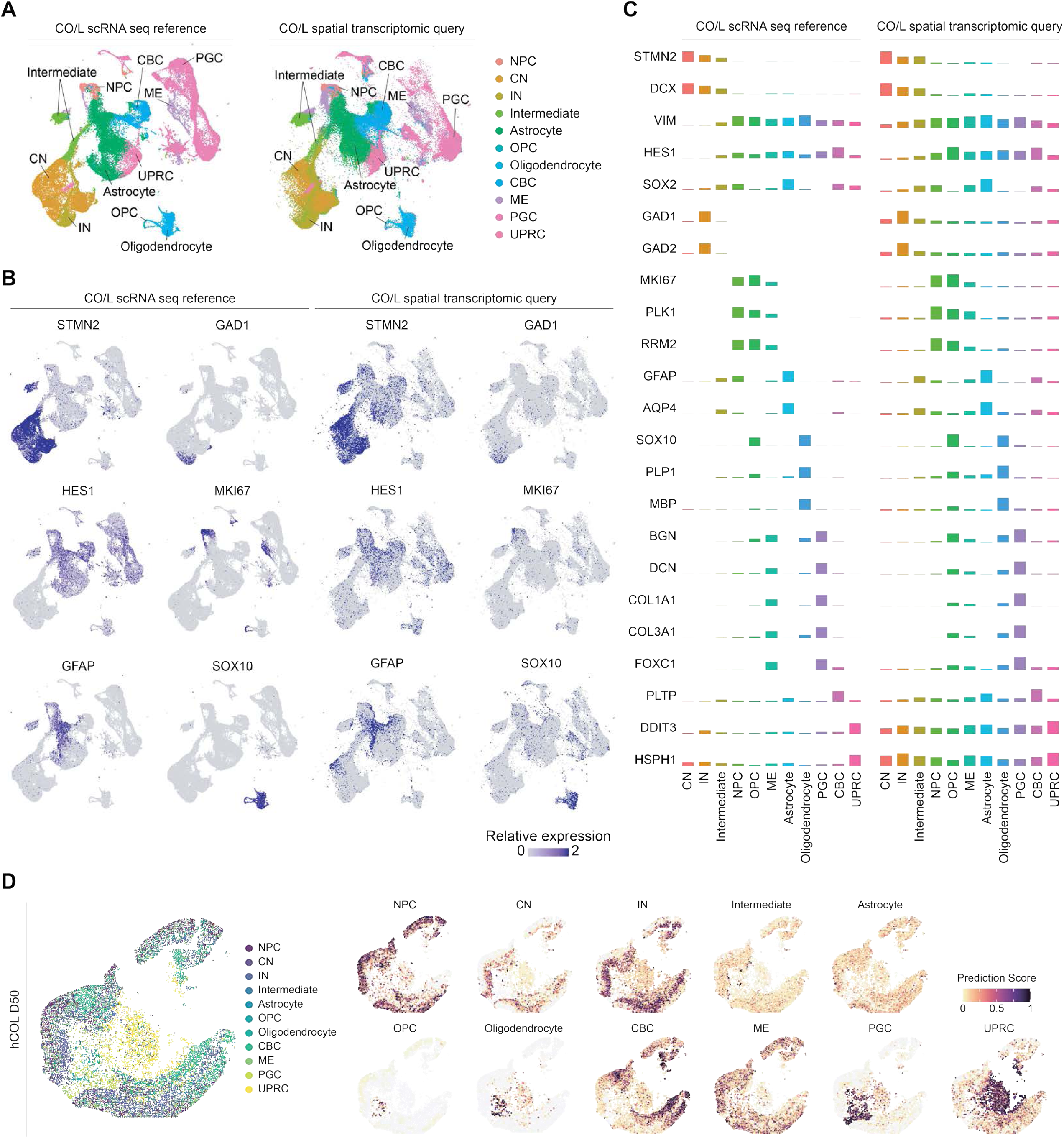
Spatial transcriptomics characterization of hCOL. (**A**) UMAP showing cell type labels transferred and projected using QueryMap. (Left) scRNA-seq (reference). (Right) spatial transcriptomics (query). (**B**) Feature plots displaying the expression of key cell type markers. (Left) scRNA-seq, reference. (Right) spatial transcriptomics, query. (**C**) Bar plots showing the expression levels of the indicated genes for each cell type. (Left) scRNA-seq, reference. (Right) spatial transcriptomics, query. (**D**) Spatial transcriptomics visualization of day 50 hCOL. (Left) Spatial plot showing regional distribution of predicted cell types. (Right) Feature plots displaying prediction scores for individual cell types.

**Figure S4.**
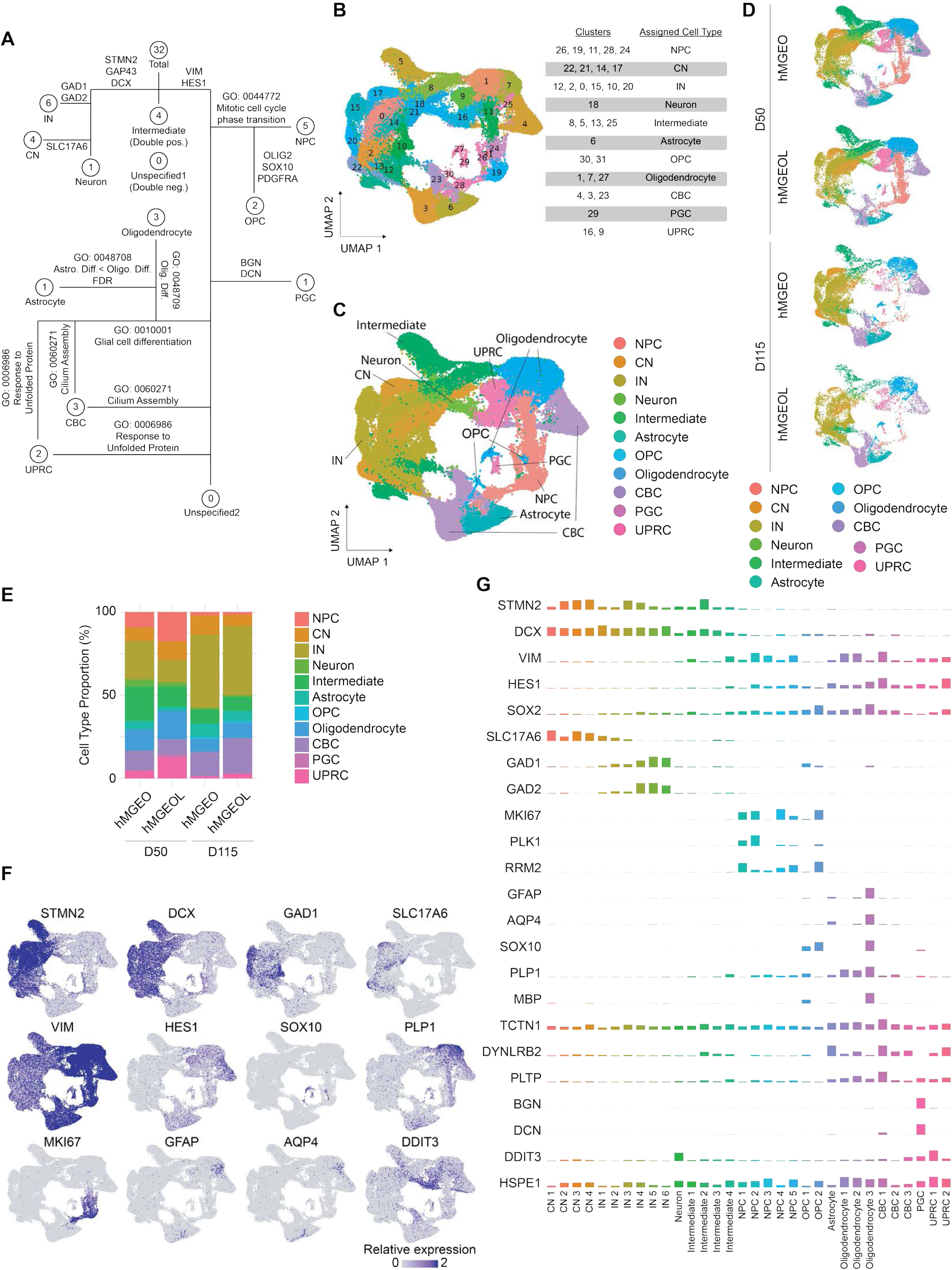
scRNA-seq characterization of hMGEOL. (**A**) Hierarchical cell type annotation decision tree used to classify hMGEOL clusters. (**B**) UMAP visualization of hMGEOL scRNA-seq data colored by cluster identity. (**C**) UMAP visualization colored by cell type annotation. (**D**) Individual UMAPs of hMGEO and hMGEOL at day 50 and day 115. (**E**) Bar plots showing the cell type composition of hMGEO and hMGEOL at day 50 and day 115. (**F**) Feature plots showing the expression of key cell type markers. (**G**) Bar plots showing the expression levels of the indicated genes for each cell type.

**Figure S5.**
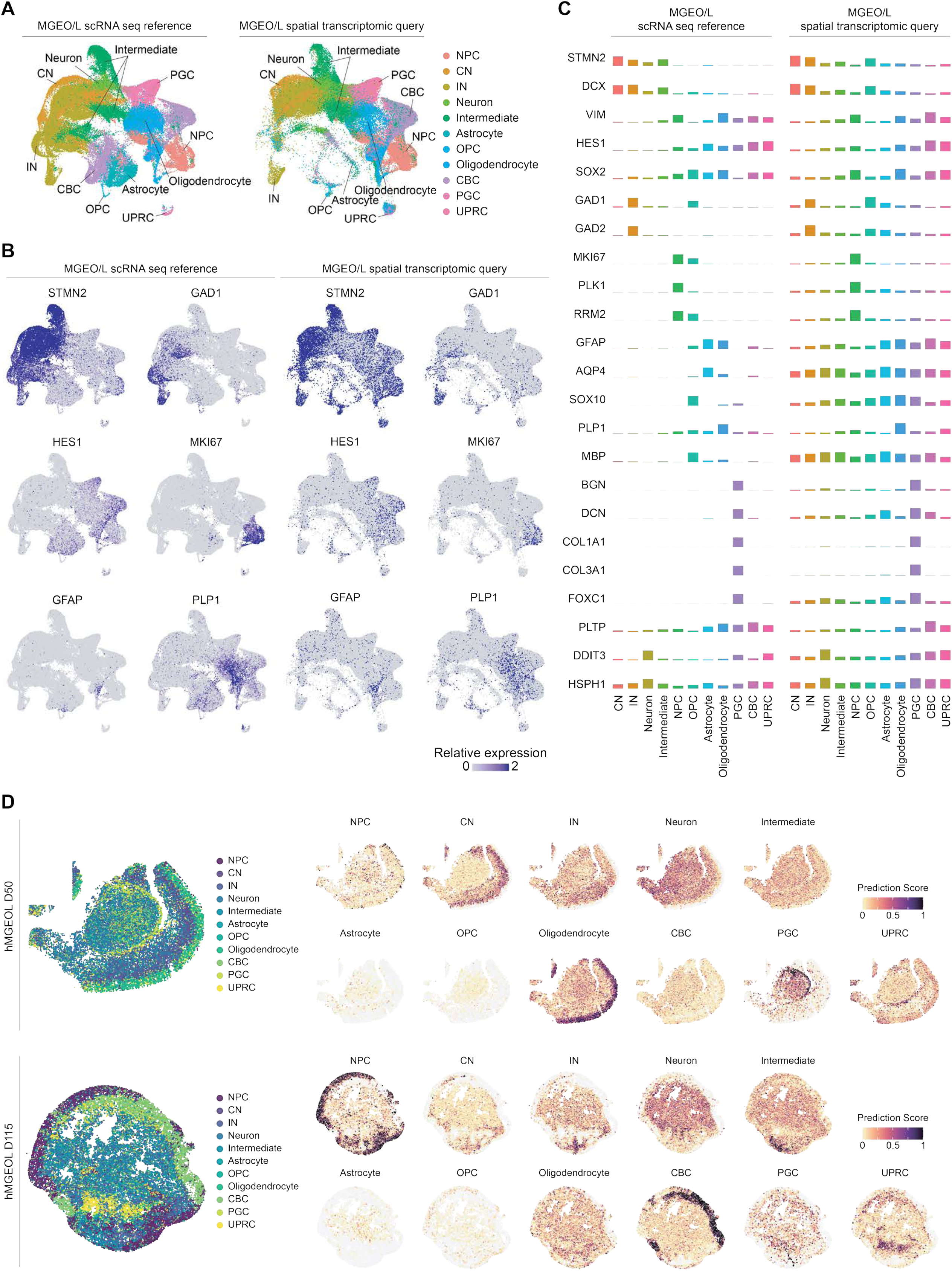
Spatial transcriptomics characterization of hMGEOL. (**A**) UMAP visualization showing the cell type annotation labels transferred and projected using QueryMap. (Left) scRNA-seq (reference). (Right) spatial transcriptomics (query). (**B**) Feature plots displaying the expression of key cell type markers. (Left) scRNA-seq, reference. (Right) spatial transcriptomics, query. (**C**) Bar plots showing the expression levels of the indicated genes for each cell type. (Left) scRNA-seq, reference. (Right) spatial transcriptomics, query. (**D**) Spatial transcriptomics visualization of hMGEOL at day 50 (top) and day 115 (bottom). (Left) Spatial plot showing regional distribution of predicted cell types. (Right) Feature plots displaying prediction scores for individual cell types.

**Figure S6.**
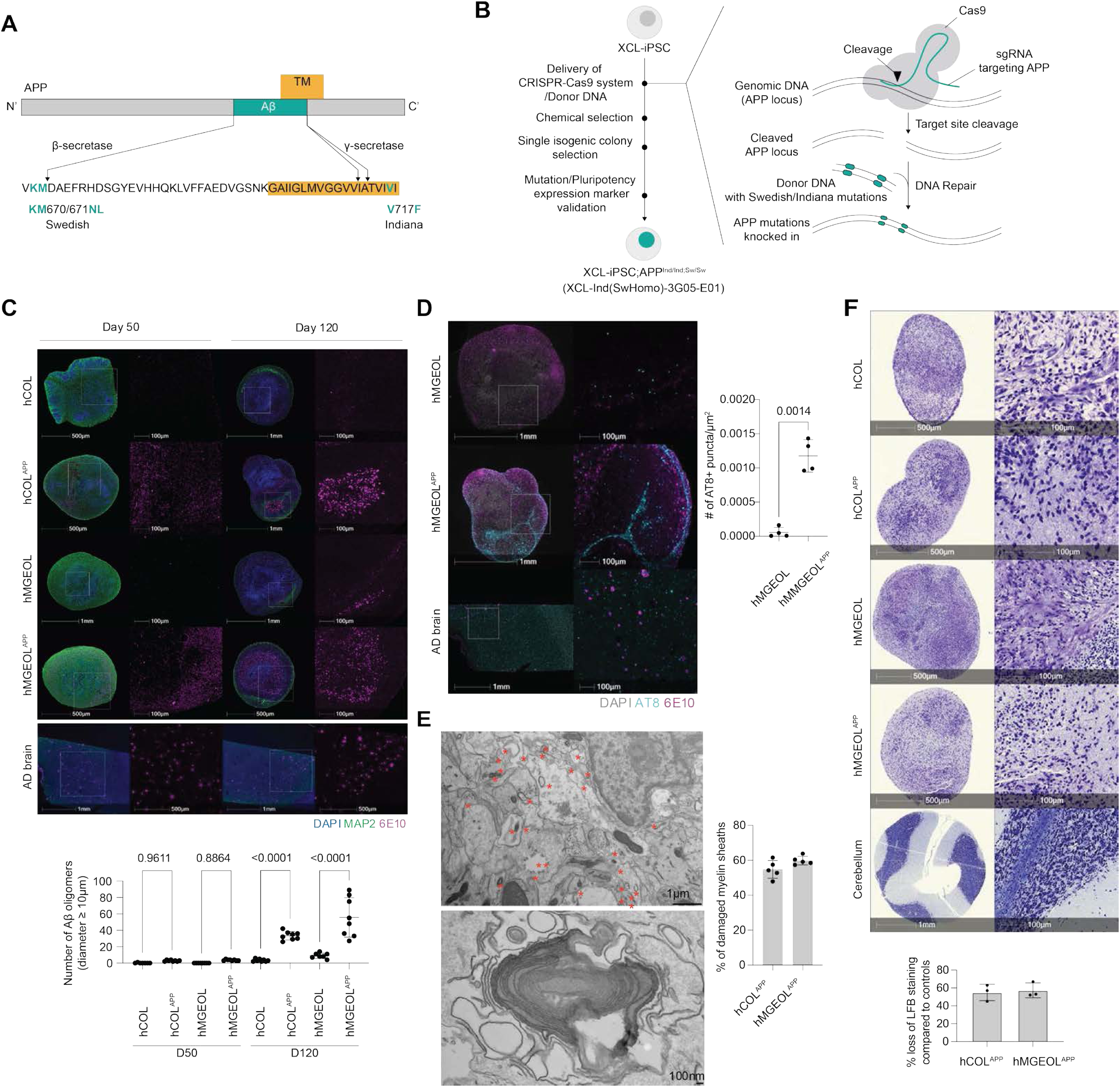
Generation and characterization of hCOL^APP^ and hMGEOL^APP^ models. (**A**) Amino acid sequence of the pathogenic familial amyloid precursor protein (APP) mutations. (**B**) Schematic diagram illustrating the generation of APP mutant XCL-iPSC line. (**C**) (Top) Immunofluorescence of amyloid-β plaques in hCOL^APP^ and hMGEOL^APP^ at day 50 and day 120, compared to human AD brain tissue. (Bottom) Quantification of Aβ oligomers (≥10 µm in diameter). N=12 organoids per group from 2 iPSC lines across 3 differentiation experiments. Mean±SD. Šidák multiple comparison test. (**D**) (Left) Immunofluorescence of tau pathology. Day 120 hMGEOL (top) and hMGEOL^APP^ (bottom) stained for phosphorylated tau (AT8, cyan) and amyloid-β (6E10, magenta), compared to human AD brain tissue. Nuclei stained with DAPI (grey). Low-magnification (left, 1 mm (top) and 500 µm (bottom)) and high-magnification (right, 100 µm). (Right) Quantification of the number of AT8+ cells per mm^2^ area. hMGEOL, 0.00006 ± 0.00007, n=4; hMGEOL^APP^, 0.001 ± 0.0002, n=4. Mean±SD. Welch’s t-test. (**E**) (Left) Electron microscopy analysis of hMGEOL^APP^. Low-magnification (top, 1 µm) and high-magnification (bottom, 100 nm). (Right) Quantification of damaged myelin sheaths (%) in hCOL^APP^ (Figure 2E) and hMGEOL^APP^. hMGEOL^APP^, n=12 organoids from 1 iPSC from 3 differentiation experiments. Mean±SD. (**F**) (Top) Luxol Fast Blue (LFB) staining of day 120 hCOL, hCOL^APP^, hMGEOL, and hMGEOL^APP^, compared to human cerebellum tissue. (Bottom) Quantification of LFB staining intensity reduction in APP-mutant organoids, compared to their matched controls. Mean±SD.

**Figure S7.**
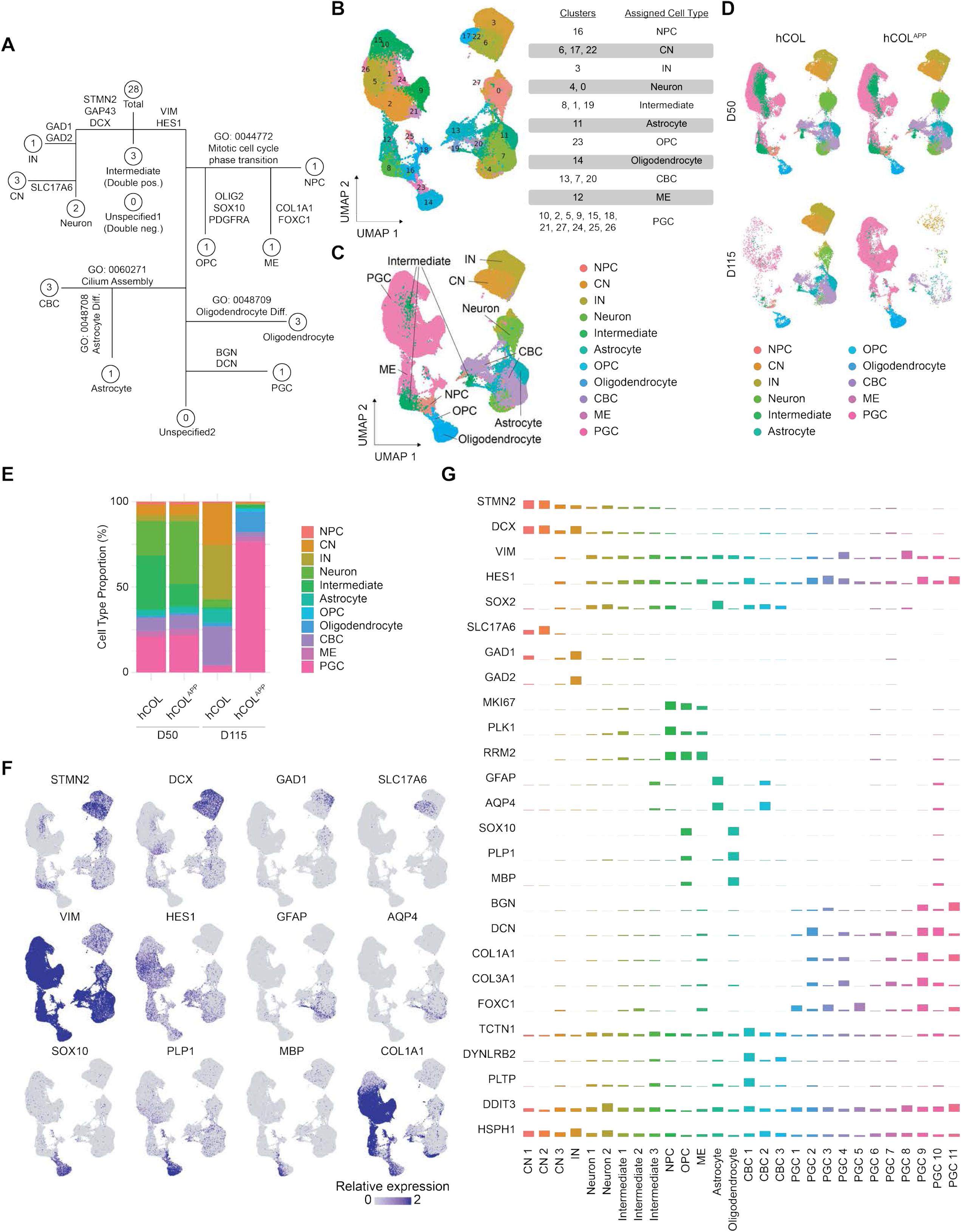
scRNA-seq analysis and cell type annotation of hCOL and hCOL^APP^. (**A**) Hierarchical cell type annotation decision tree used to classify clusters in the hCOL and hCOL^APP^ dataset. (**B**) UMAP visualization of hCOL and hCOL^APP^ colored by cell clusters. (**C**) UMAP visualization of hCOL and hCOL^APP^ colored by cell type annotation. (**D**) Individual UMAPs of hCOL and hCOL^APP^ at day 50 and day 115. (**E**) Stacked bar plots of cell type composition of hCOL and hCOL^APP^ at day 50 and day 115. (**F**) Feature plots of canonical cell type markers. (**G**) Bar plots showing the expression levels of selected genes for each cell type.

**Figure S8.**
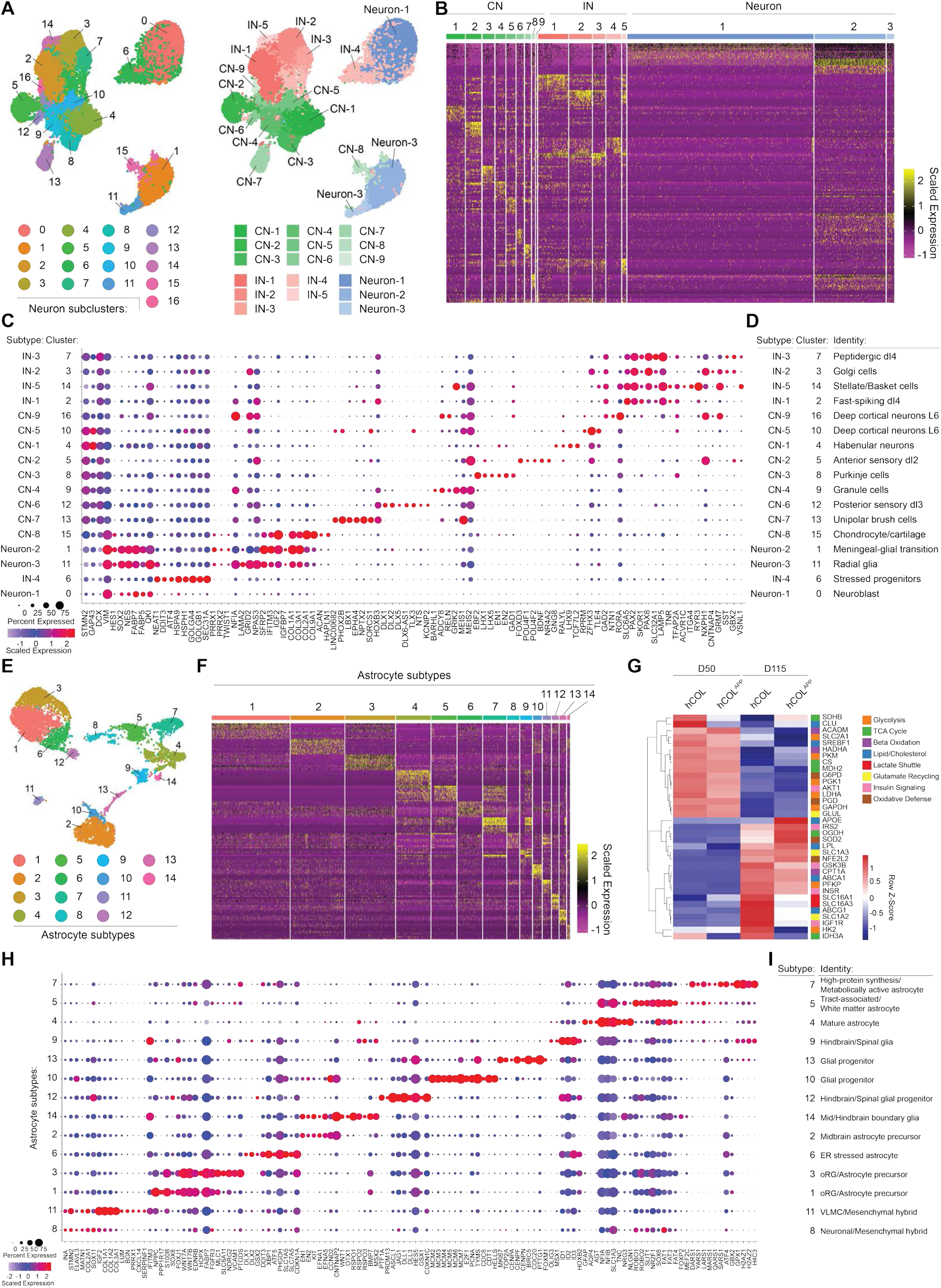
Subclustering and detailed characterization of neuron and astrocyte subtypes in hCOL and hCOL^APP^. (**A**) UMAP of hCOL and hCOL^APP^ neuronal subtypes colored by (Left) subclusters and (Right) parental cell labels, with clusters numerically annotated. (**B**) Heatmap of top 10 differentially expressed genes for each neuronal subtype. (**C**) Dot plot showing the differential expression of key genes used to identify neuronal subtype characteristics. (**D**) Annotation of neuronal subtype identity based on the differential gene expression profiles from (**B**) and (**C**). (**E**) UMAP of hCOL and hCOL^APP^ astrocyte subtypes colored by subclusters. (**F**) Heatmap of top 10 differentially expressed genes for each astrocyte subtype. (**G**) Heatmap displaying the expression of key metabolic genes across hCOL and hCOL^APP^ astrocyte subtypes at day 50 and day 115. (**H**) Dot plot showing the differential expression of key genes used to identify astrocyte subtype features. (**I**) Annotation of astrocyte subtype identity based on the differential gene expression profiles from (**F**) and (**H**).

**Figure S9.**
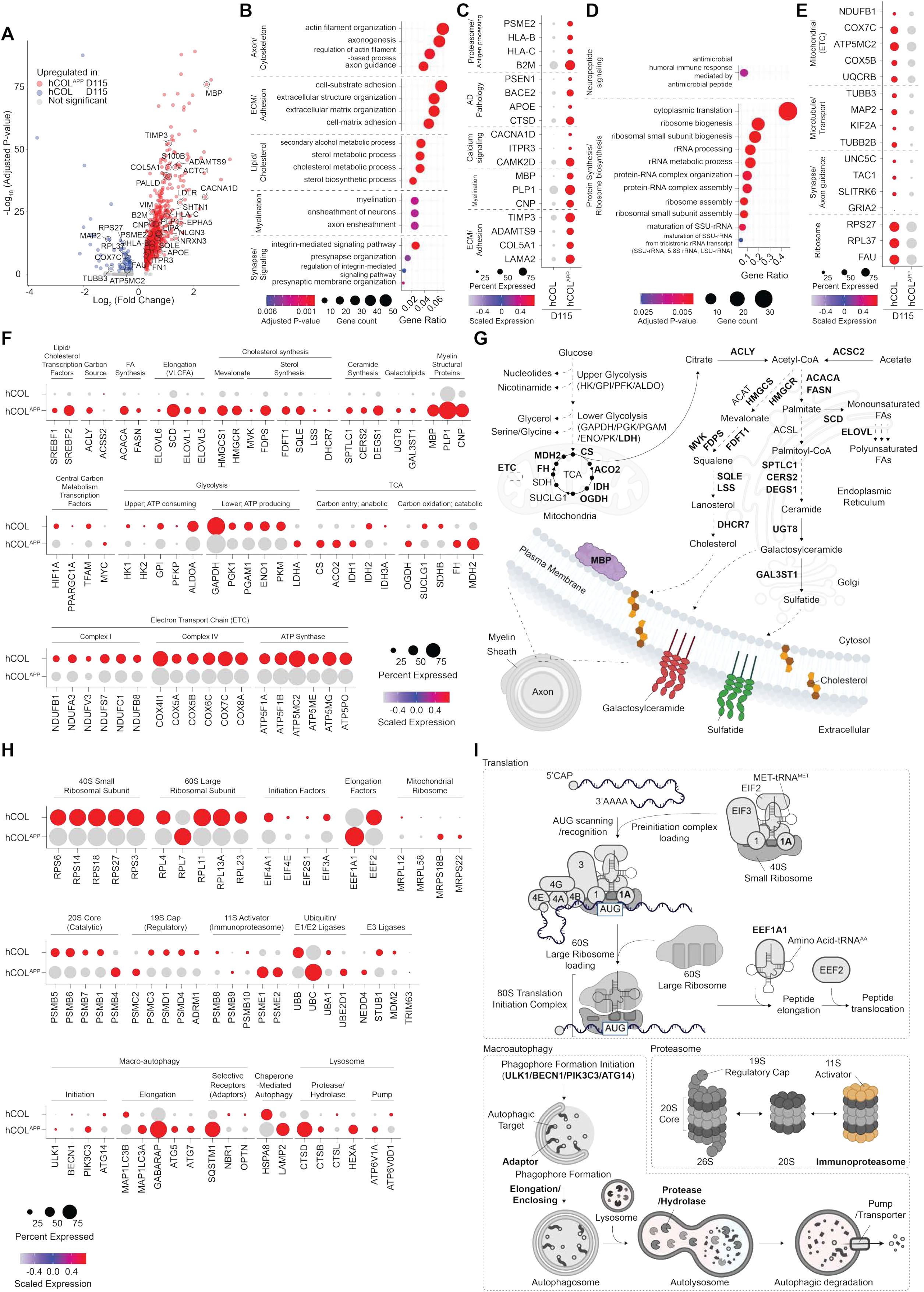
Altered transcript levels of myelin biosynthesis, central carbon metabolism, protein translation, and protein catabolism pathways in day 115 hCOL^APP^ oligodendrocytes. (**A**) Volcano plot of differentially expressed genes comparing day 115 hCOL and hCOL^APP^. Genes highlighting upregulated (**B**) and downregulated (**C**) pathways are annotated. (**B**) Gene Ontology (GO) enrichment analysis for genes upregulated in hCOL^APP^ oligodendrocytes. (**C**) Dot plot of select upregulated genes associated with annotated cellular activities (**B**) and other relevant pathways. (**D**) GO enrichment analysis for genes downregulated in hCOL^APP^ oligodendrocytes. (**E**) Dot plot of select downregulated genes involved in the annotated cellular activities (d). (**F**) Dot plot showing the expression of key genes involved in lipid/myelin biosynthesis and central carbon metabolism (glycolysis, tricarboxylic acid (TCA) cycle, electron transport chain (ETC)), comparing D115 hCOL and hCOL^APP^. (**G**) Schematic diagram showing the metabolic pathways transcriptionally altered in hCOL^APP^ oligodendrocytes (**F**). Genes shown in (**F**) are bolded. (**H**) Dot plot showing the expression of key genes involved in ribosome biogenesis/translation and protein catabolic pathways including proteasomal and autophagic degradation. (**I**) Schematic diagram showing the translation and protein catabolic pathways transcriptionally altered in hCOL^APP^ oligodendrocytes (**H**). Genes shown in (**H**) are bolded.

**Figure S10.**
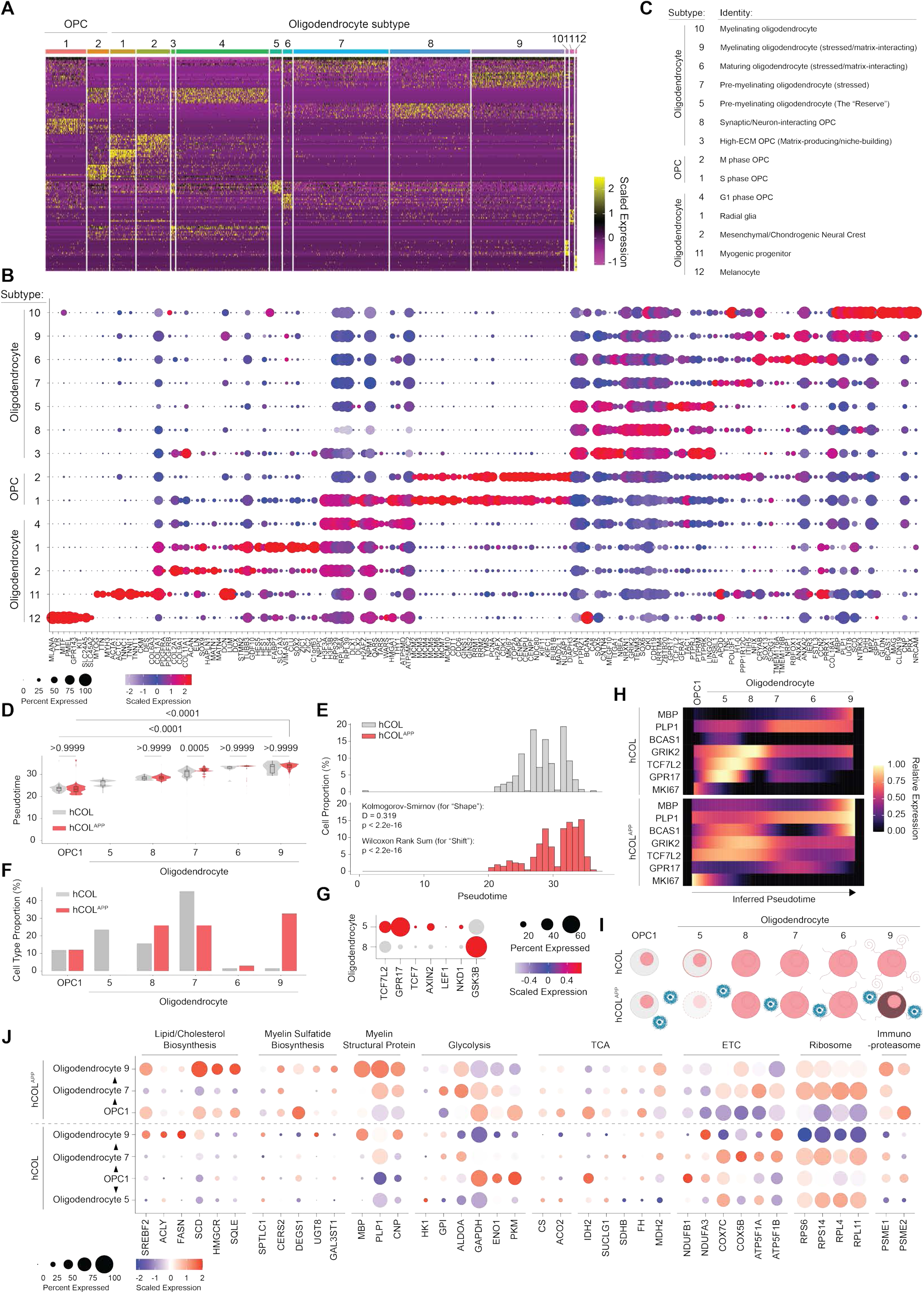
Detailed characterization of oligodendrocyte subtypes and maturation dynamics in hCOL and hCOL^APP^. (**A**) Heatmap of top 10 differentially expressed genes for each oligodendrocyte and OPC subtype. (**B**) Dot plot showing the differential expression of key genes used to identify oligodendrocyte subtype characteristics. (**C**) Annotation of oligodendrocyte subtype identity based on the differential gene expression profiles from (**A**) and (**B**). (**D**) Pseudotime trajectory analysis comparing the differentiation of day 115 hCOL and hCOL^APP^ oligodendrocyte subtypes. Median with interquartile range (IQR). Kruskal–Wallis one-way ANOVA (nonparametric). (**E**) Histogram showing the distribution of D115 hCOL and hCOL^APP^ cells along the pseudotime trajectory. Kolmogorov-Smirnov test and Wilcoxon Rank Sum test. (**F**) Bar plots quantifying the proportion of OPC and oligodendrocyte subtypes in D115 hCOL and hCOL^APP^. (**G**) Dot plot comparing canonical WNT signaling target and GSK3β gene expression in oligodendrocyte subtypes 5 and 8. (**H**) LOESS trajectory smoothing of key oligodendrocyte differentiation and maturation marker genes ordered by pseudotime. (**I**) Schematic diagram illustrating the differential differentiation trajectories of oligodendrocytes in hCOL versus hCOL^APP^. (**J**) Dot plot showing the expression of genes involved in lipid/myelin synthesis, glycolysis, TCA cycle, ETC, ribosome biogenesis, and the immunoproteasome across oligodendrocyte subclusters in day 115 hCOL and hCOL^APP^.

**Figure S11.**
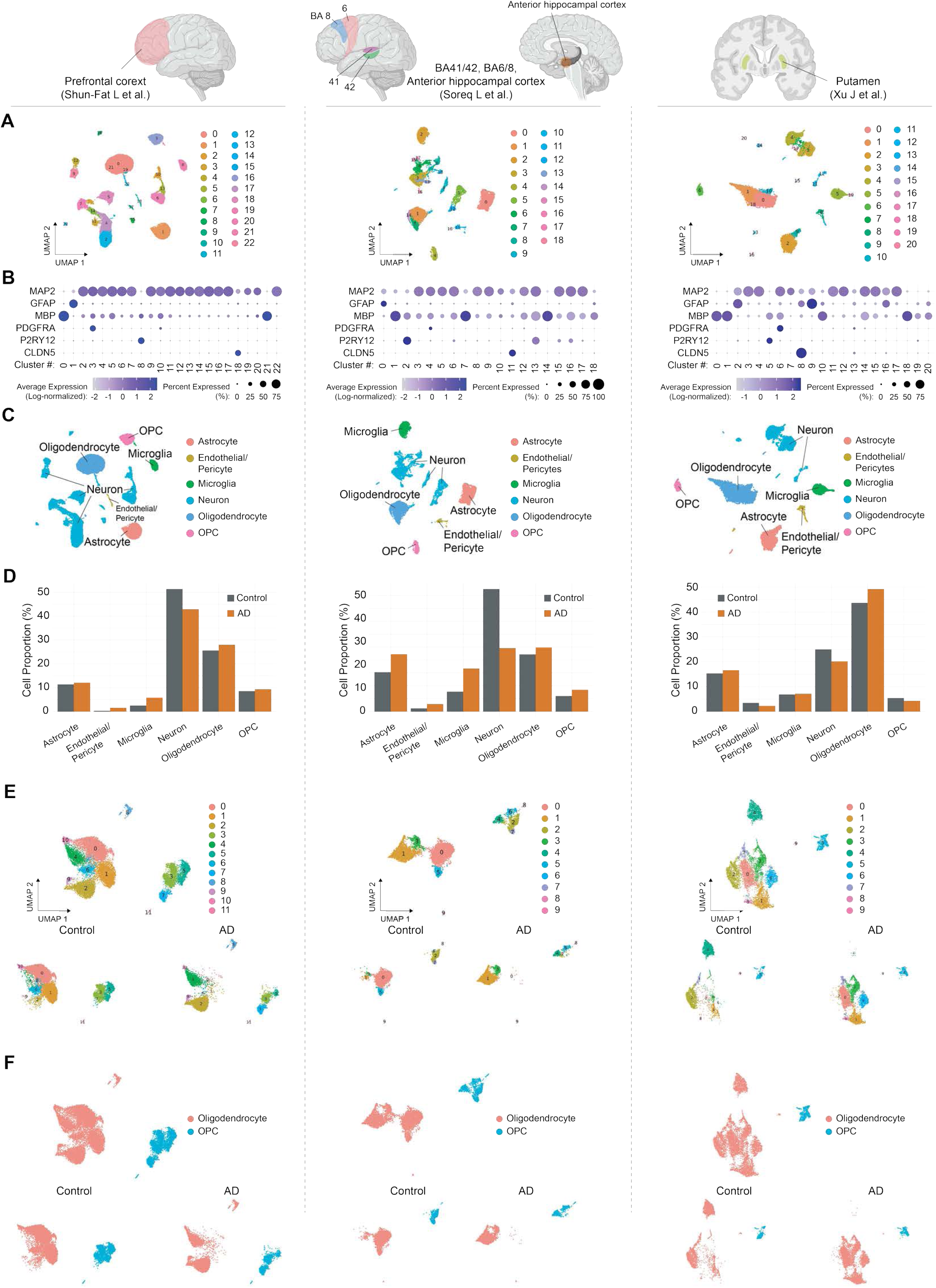
Analysis of human reference datasets from prefrontal cortex, BA41/42, BA6/8, anterior hippocampal cortex, and putamen. (**A**) UMAP visualizations of integrated scRNA-seq datasets from three human brain regions: (Left) Prefrontal cortex (Shun-Fat L et al.). (Middle) BA41/42, BA6/8, and Anterior hippocampal cortex (Soreq L et al.). (Right) Putamen (Xu J et al.). Anatomical locations of the brain regions are highlighted. (**B**) Dot plot showing the expression of key cell type markers used for cell annotation. (**C**) UMAP colored by cell annotation based upon the canonical cell marker expression in (**B**). (**D**) Bar plots quantifying the proportion of cell types in Control and AD. (**E**) (Top) UMAP of oligodendrocyte and OPC subclusters colored by clusters. (Bottom) UMAP split by condition (Control, left; AD, right). (**F**) (Top) UMAP of oligodendrocyte and OPCs colored by cell annotation. (Bottom) UMAP split by condition (Control, left; AD, right).

**Figure S12.**
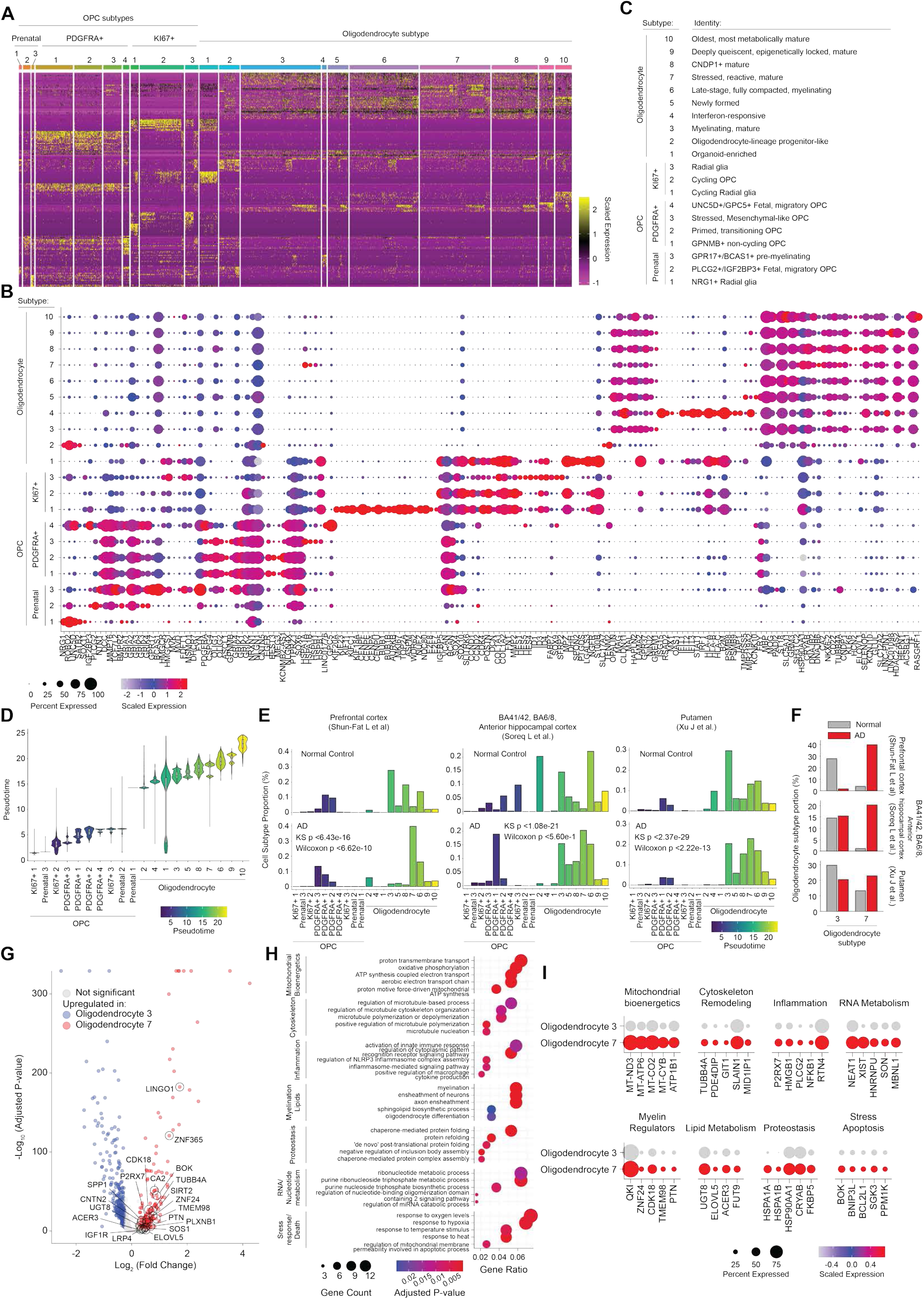
Integrated analysis of oligodendrocytes in hCOL/hCOL^APP^ and human reference datasets. (**A**) Heatmap of top 10 differentially expressed genes for each oligodendrocyte and OPC subtypes from an integrated dataset comprising hCOL, hCOL^APP^, 3 different human Control/AD brain regions (Fig. S11), and human prenatal brain (Ramos SI et al.). (**B**) Dot plot showing the differential expression of key genes used to identify oligodendrocyte OPC subtype characteristics. (**C**) Annotation of oligodendrocyte subtype identity based on the differential gene expression profiles from (**A**) and (**B**). (**D**) Violin plots of pseudotime scores for the oligodendrocyte lineage across the integrated dataset. Median with interquartile range (IQR). (**E**) Histograms showing the distribution of oligodendrocytes and OPCs along the pseudotime trajectory for Prefrontal cortex (Shun-Fat et al., left), BA41/42, BA6/8, Anterior hippocampal cortex (Soreq et al., middle), and Putamen (Xu et al., right), split by Normal Control (top) and AD (bottom). Kolmogorov-Smirnov test and Wilcoxon Rank Sum test. (**F**) Bar plots quantifying the proportion of oligodendrocyte subtypes 3 and 7 in Normal Control and AD conditions across the three human brain regions. (**G**) Volcano plot comparing gene expression between oligodendrocyte subtype 3 and subtype 7, with highlighted genes involved with oligodendrocyte differentiation. (**H**) Dot plot of Gene Ontology enrichment analysis for genes upregulated in oligodendrocyte subtype 7, which is enriched in the three human AD brain regions. (**I**) Dot plot showing the expression of example genes upregulated in oligodendrocyte subtype 7, corresponding to the pathways highlighted in (**H**).

**Figure S13.**
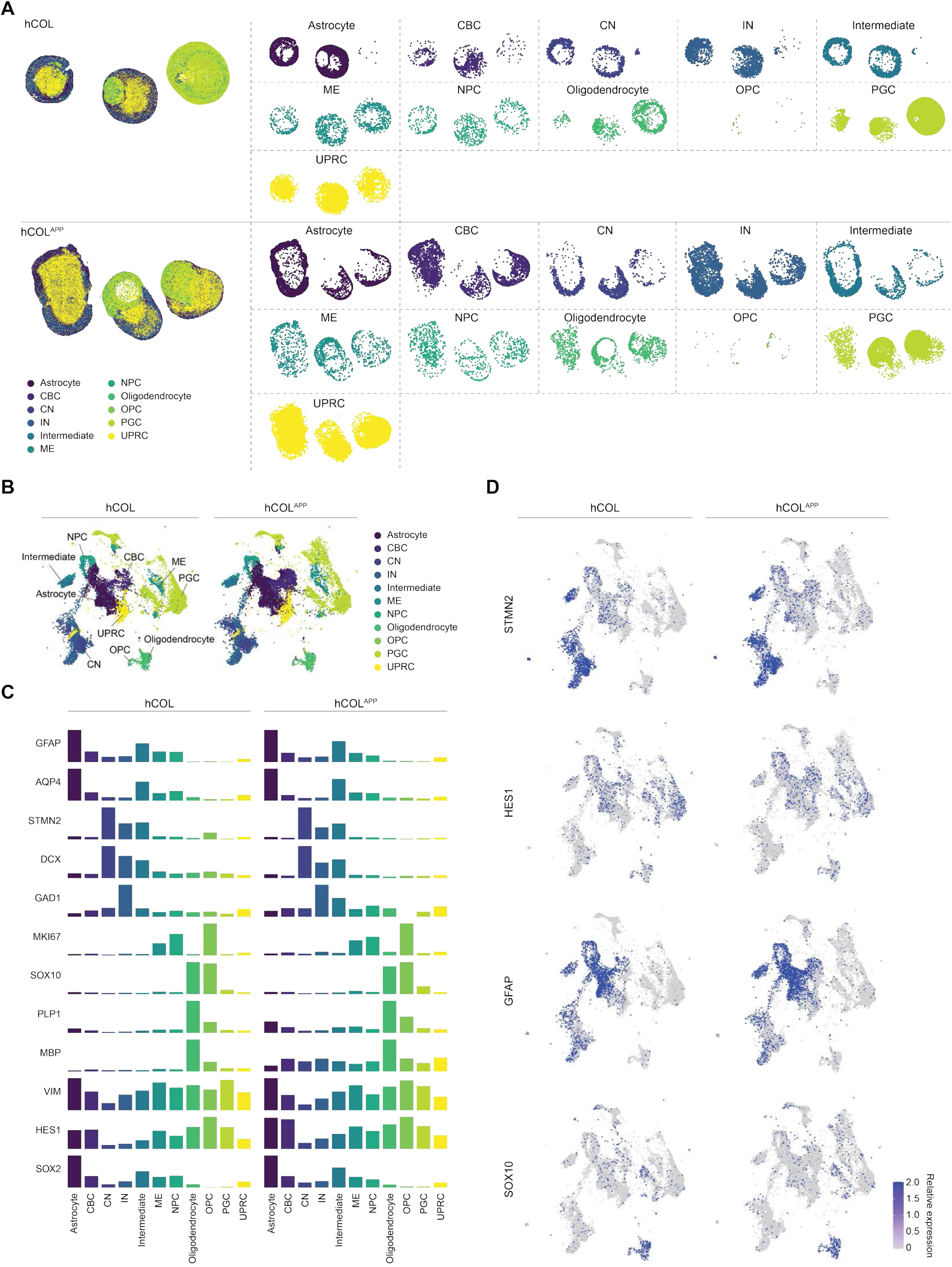
Spatial transcriptomics analysis of hCOL and hCOL^APP^ at day 115. (**A**) Spatial visualization showing the regional distribution of all cell types (left) and individual cell types (right) in day 115 hCOL (top) and hCOL^APP^ (bottom) sections. (**B**) Individual UMAPs of day 115 hCOL and hCOL^APP^ spatial transcriptomics, colored by cell type annotation transferred from scRNA-seq using MapQuery. (**C**) Bar plots showing the expression levels of selected genes for each cell type of day 115 hCOL (left) and hCOL^APP^ (right) spatial transcriptomics. (**D**) Feature plots displaying, in UMAP, the expression of key cell type markers of day 115 hCOL (left) and hCOL^APP^ (right) spatial transcriptomics.

**Figure S14.**
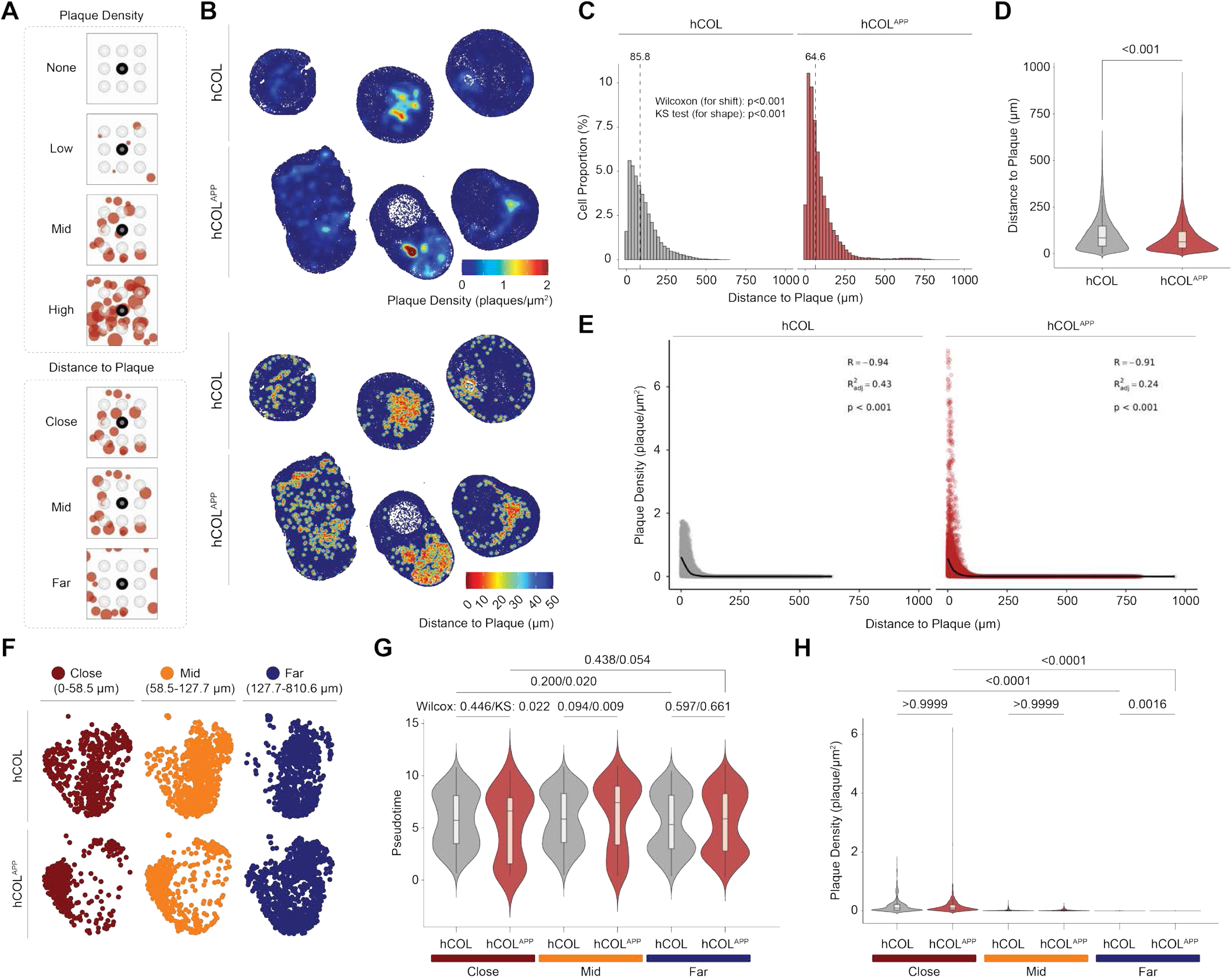
Detailed analysis of plaque density and proximity in spatial transcriptomics. (**A**) Schematic diagram defining parameters for plaque density (top) and distance to plaque (bottom) used in the spatial analysis. Plaque density is computed by convolving the plaque binary mask with a Gaussian kernel, yielding a continuous density value (µm⁻²) at each cell centroid that reflects the cumulative influence of all nearby plaques weighted by distance. Distance to plaque is computed as the Euclidean distance transform of the plaque binary mask, representing the shortest physical distance from each cell centroid to the nearest plaque. Both metrics are subsequently discretized into tertiles (Low, Mid, High for density; Close, Mid, Far for distance) using thresholds defined on the full spatial dataset. (**B**) Spatial visualization mapping plaque density (top) and distance to the nearest plaque (bottom) in day 115 hCOL and hCOL^APP^. (**C**) Histogram showing the distribution of cells relative to their distance to the nearest plaque. hCOL, left. hCOL^APP^, right. Kolmogorov-Smirnov test and Wilcoxon Rank Sum test. (**D**) Violin plot of the distance to plaque for cells in hCOL and hCOL^APP^. Median with interquartile range (IQR). Mann-Whitney test. (**E**) Dot plot showing cell density as a function of distance to plaque, with a Generalized Additive Model (GAM) trend line (black). Pearson correlation (p-value). (**F**) UMAP of hCOL and hCOL^APP^ oligodendrocytes/OPCs, split by distance tertiles (close, mid, far). (**G**) Violin plot comparing pseudotime scores of oligodendrocytes and OPCs in hCOL and hCOL^APP^ across distance tertiles. Median with interquartile range (IQR). Kolmogorov-Smirnov test (left) and Wilcoxon Rank Sum test (right). (**H**) Violin plot comparing plaque density across distance tertiles in hCOL and hCOL^APP^. Median with interquartile range (IQR). Kruskal–Wallis one-way ANOVA (nonparametric).

**Figure S15.**
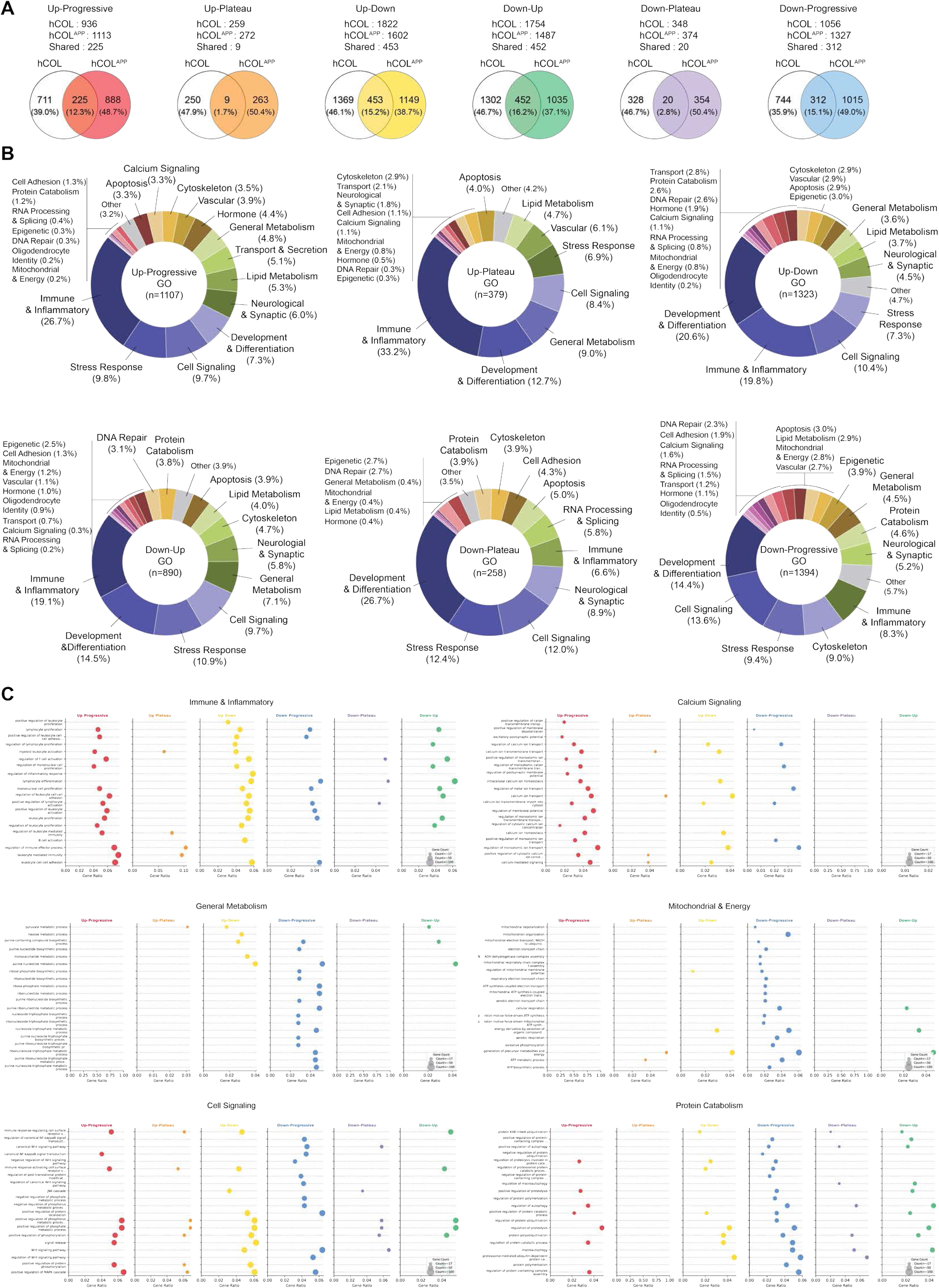
Gene Ontology biological process enrichment analysis of plaque-associated trajectory patterns exclusively in hCOL^APP^ oligodendrocytes. (**A**) Venn diagrams showing the overlap of genes assigned to each of the six trajectory patterns between hCOL and hCOL^APP^ oligodendrocytes. Gene counts and percentages are shown for hCOL-specific, shared, and hCOL^APP^-specific genes for each pattern: Up-Progressive, Up-Plateau, Up-Down, Down-Up, Down-Plateau, and Down-Progressive. (**B**) Donut charts showing the distribution of GO biological process categories significantly enriched among genes assigned to each of the six trajectory patterns exclusively in hCOL^APP^ oligodendrocytes. Each chart summarizes the proportion of enriched GO terms belonging to major biological process categories including immune and inflammatory response, stress response, cell signaling, development and differentiation, neurological and synaptic processes, lipid metabolism, transport and secretion, general metabolism, calcium signaling, protein catabolism, and mitochondrial and energy metabolism, among others. (**C**) Dot plots of GO biological process enrichment across all six trajectory patterns in hCOL^APP^ oligodendrocytes, shown for six pathway categories of interest: Immune & Inflammatory, Calcium Signaling, General Metabolism, Mitochondrial & Energy, Cell Signaling, and Protein Catabolism.

**Figure S16.**
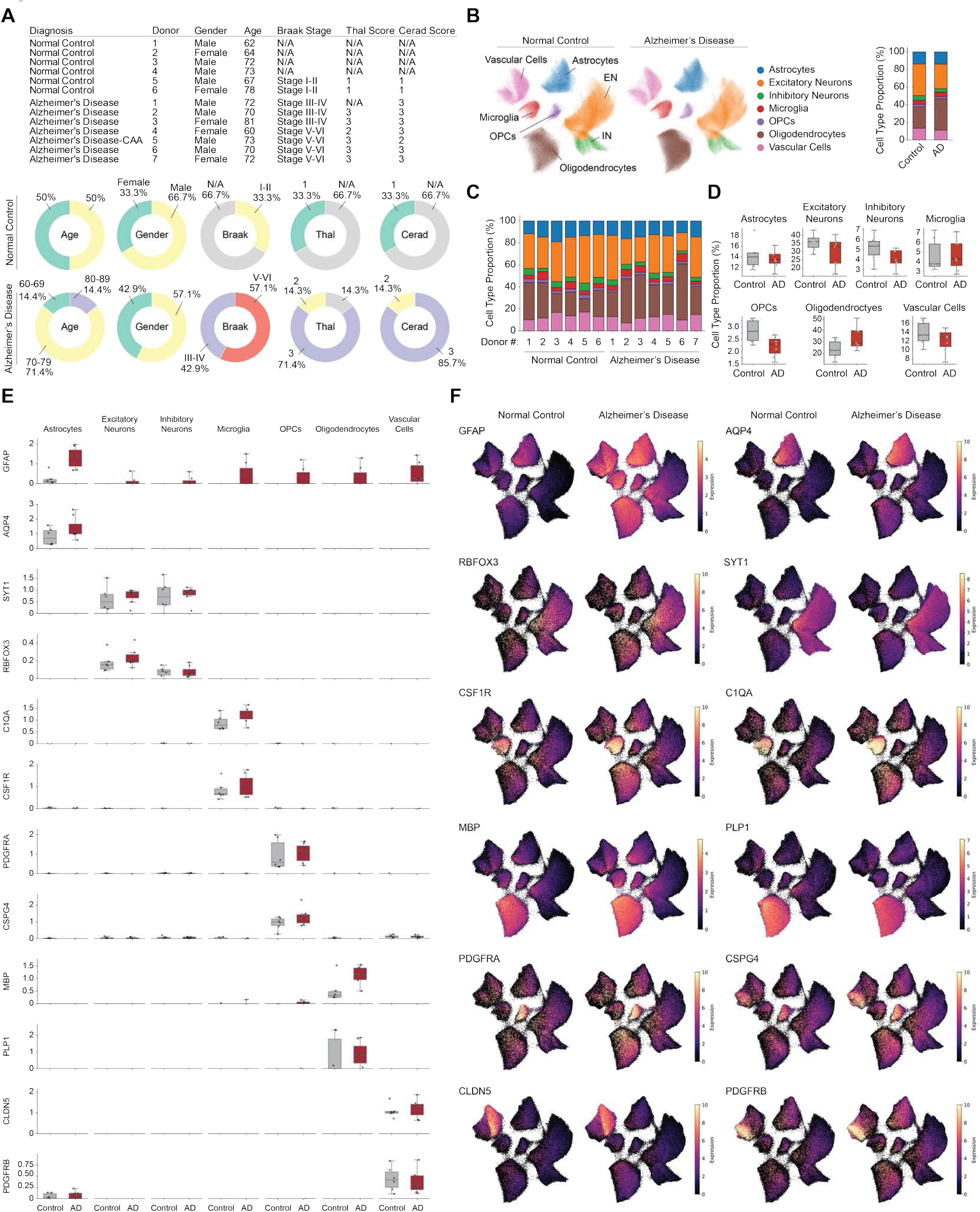
Clinical and spatial transcriptomic cell type profiles of human unaffected control and AD donors. (**A**) (Top) Table summarizing clinical profiles of 6 unaffected control and 7 AD donors, including study-assigned donor identification number, gender, age, Braak stage, Thal score, and CERAD score. (Bottom) Donut charts summarizing the distribution of age, gender, Braak stage, Thal score, and CERAD score across control and AD donors. (**B**) (Left) UMAP of all cell types identified from human control and AD spatial transcriptomics, colored by cell type annotation. (Right) Stacked bar plot showing cell type proportions in control and AD. (**C**) Stacked bar plots showing cell type proportions per individual donor across the 6 control and 7 AD donors. (**D**) Box plots comparing the proportion of each cell type between control and AD donors. Each dot represents one donor. Mean±SD. (**E**) Box plots showing expression of canonical cell type marker genes across cell types in control (grey) and AD (red): GFAP and AQP4 (astrocytes), SYT1 and RBFOX3 (neurons), C1QA and CSF1R (microglia), PDGFRA and CSPC4 (OPCs), MBP and PLP1 (oligodendrocytes), CLDN5 and PDGFRB (vascular cells). Each dot represents one donor. Mean±SD. (**F**) Feature plots showing expression of canonical cell type markers on UMAPs split by diagnosis.

**Figure S17.**
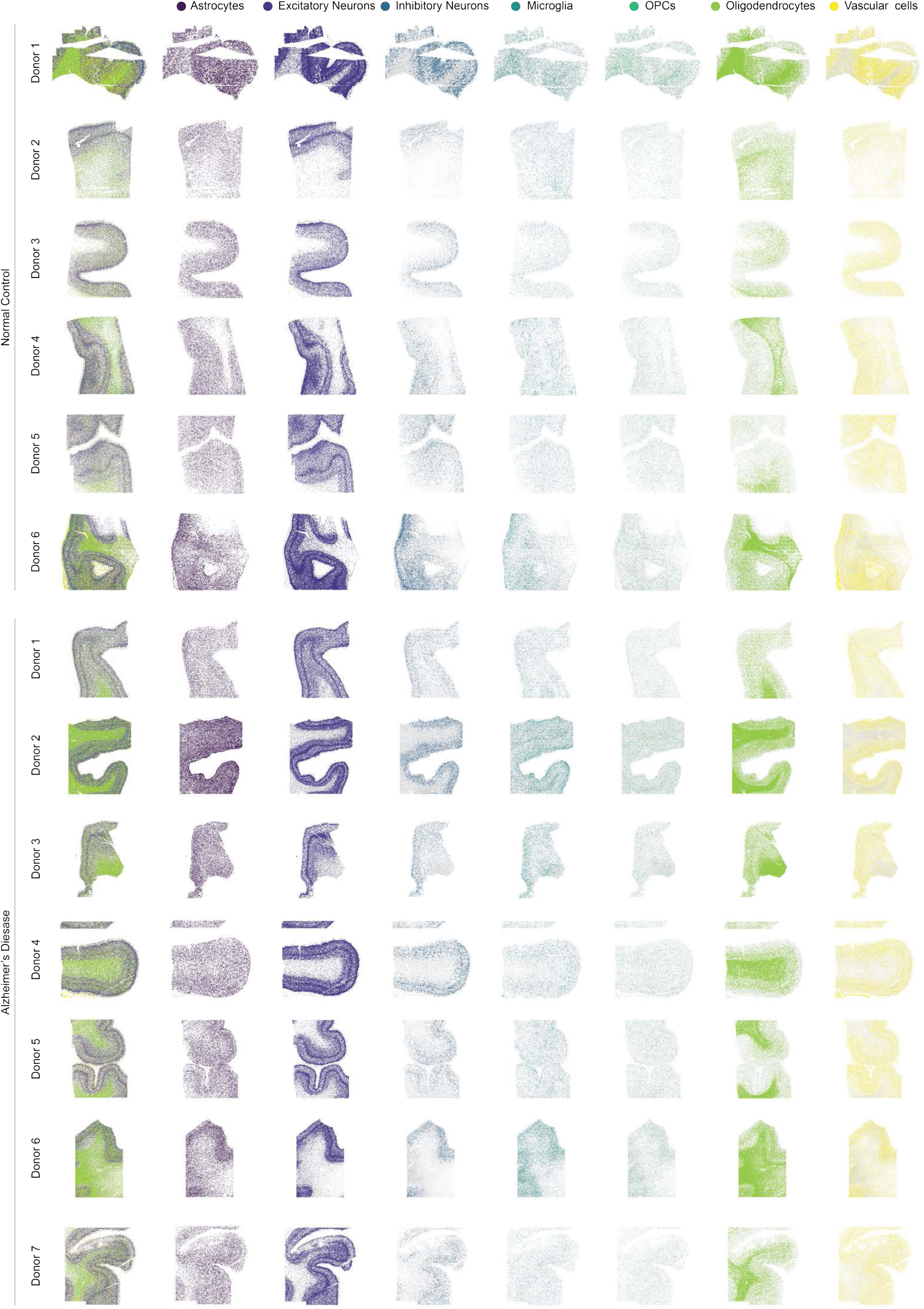
Spatial distribution of cell types across all human control and AD donor brain sections. Spatial visualization showing the distribution of each annotated cell type across individual tissue sections from all 13 donors (6 unaffected control and 7 AD donors). Each row represents one donor and each column represents one cell type, except for the first column, showing all cell types overlaid. Each cell type shown in a distinct color.

**Figure S18.**
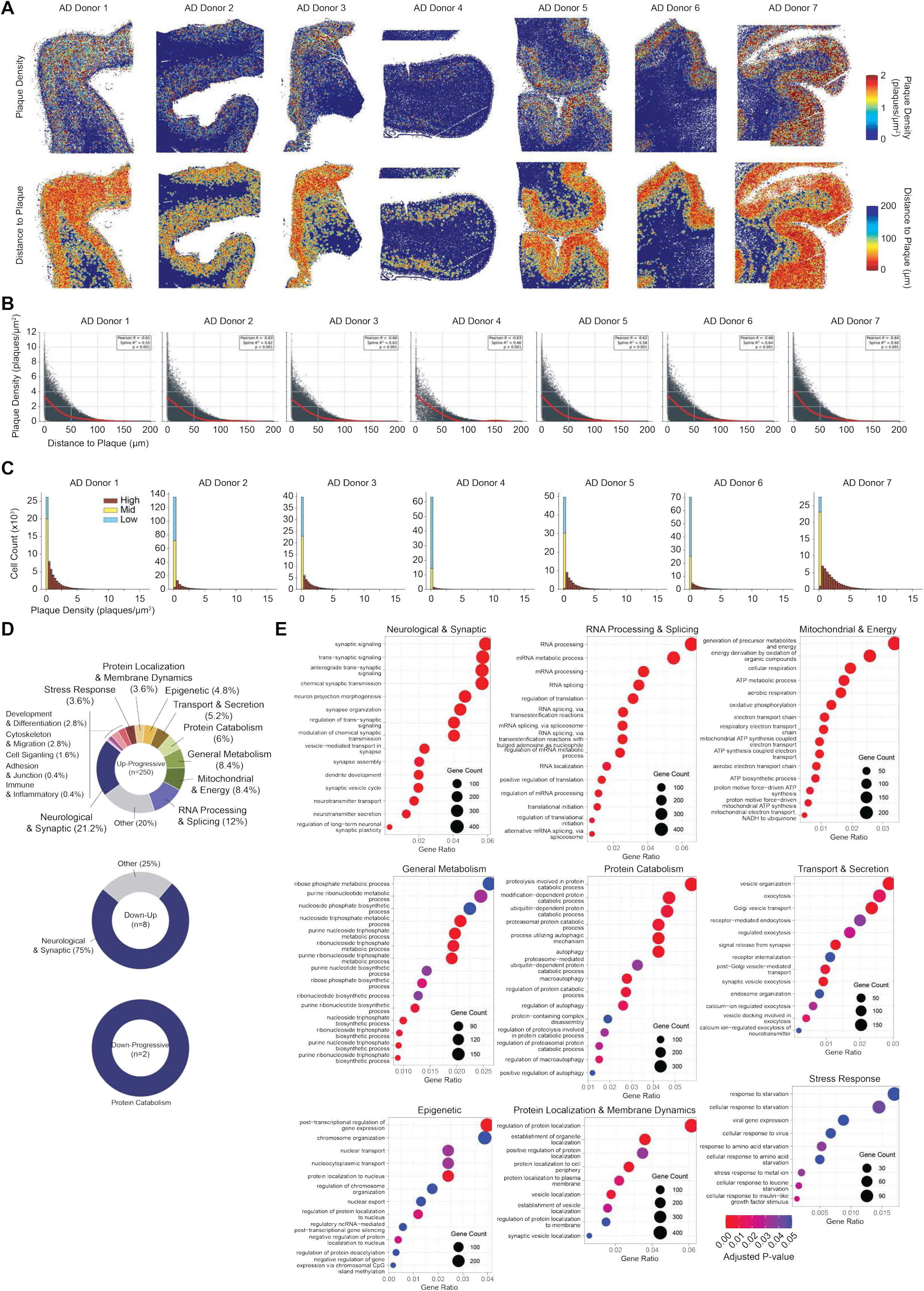
Plaque density and proximity analysis, and GO biological process enrichment analysis in human AD donors. (**A**) Spatial visualization of plaque density (top row) and distance to nearest plaque (bottom row) across tissue sections from all 7 AD donors. (**B**) Dot plot showing cell density as a function of distance to plaque for each of the 7 AD donors, with a Generalized Additive Model (GAM) trend line (black). Pearson correlation (p-value). (**C**) Histograms showing the distribution of cells across plaque density stratified by plaque density tertile for each of the 7 AD donors. (**D**) Donut charts showing GO biological process categories enriched among plaque-associated genes for trajectory patterns with statistically significant enrichment in human AD oligodendrocytes. (**E**) Dot plots of GO biological process enrichment for Up-Progressive genes in human AD oligodendrocytes, organized by pathway categories.

**Figure S19.**
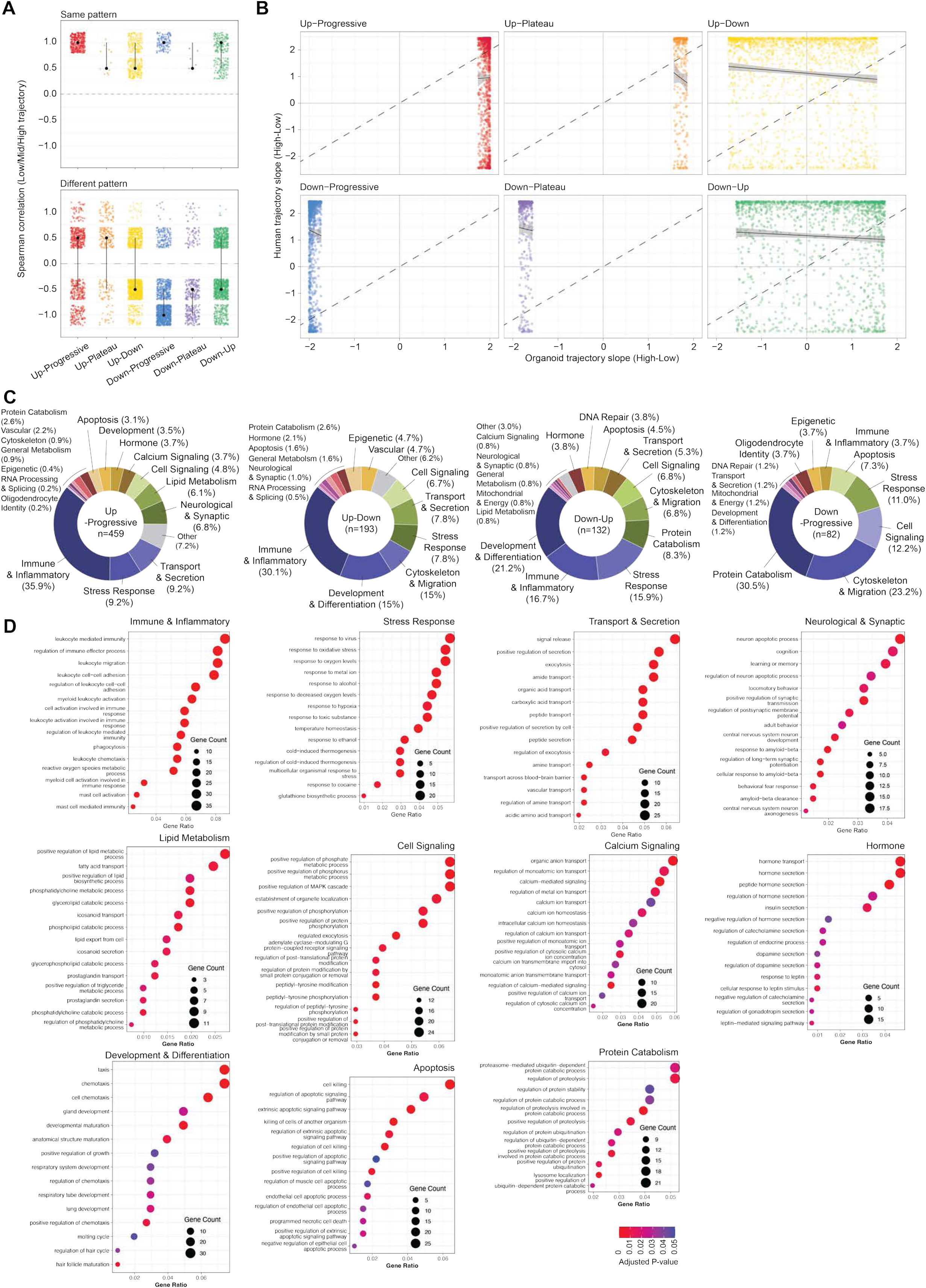
Cross-model conservation of plaque-associated transcriptional programs between organoid and human AD oligodendrocytes. (**A**) (Top) Raincloud plots of Spearman correlations of plaque density-dependent gene trajectories between organoid and human AD oligodendrocytes for genes assigned to the same pattern category in both models. (Bottom) Raincloud plots for genes assigned to different pattern categories between the two models. Each color represents one of the six trajectory patterns. Crossbar indicates median (black dot) with IQR. (**B**) Scatter plots of organoid versus human AD trajectory slopes (High-Low z-score) for genes within each of the six pattern categories. Each dot represents one gene colored by its pattern assignment. Dashed diagonal indicates perfect concordance. Linear regression trend line shown with 95% confidence interval. (**C**) Donut chart showing GO biological process categories enriched among shared plaque-associated genes for each of the four pattern categories showing significant conservation between organoid and human AD oligodendrocytes: Up-Progressive, Up-Down, Down-Up, and Down-Progressive (**D**) Dot plots of GO biological process enrichment for shared Up-Progressive genes conserved between organoid and human AD oligodendrocytes, organized by biological process category: Immune & Inflammatory, Stress Response, Transport & Secretion, Neurological & Synaptic, Lipid Metabolism, Cell Signaling, Calcium Signaling, Hormone, Development & Differentiation, Apoptosis, and Protein Catabolism.

**Figure S20.**
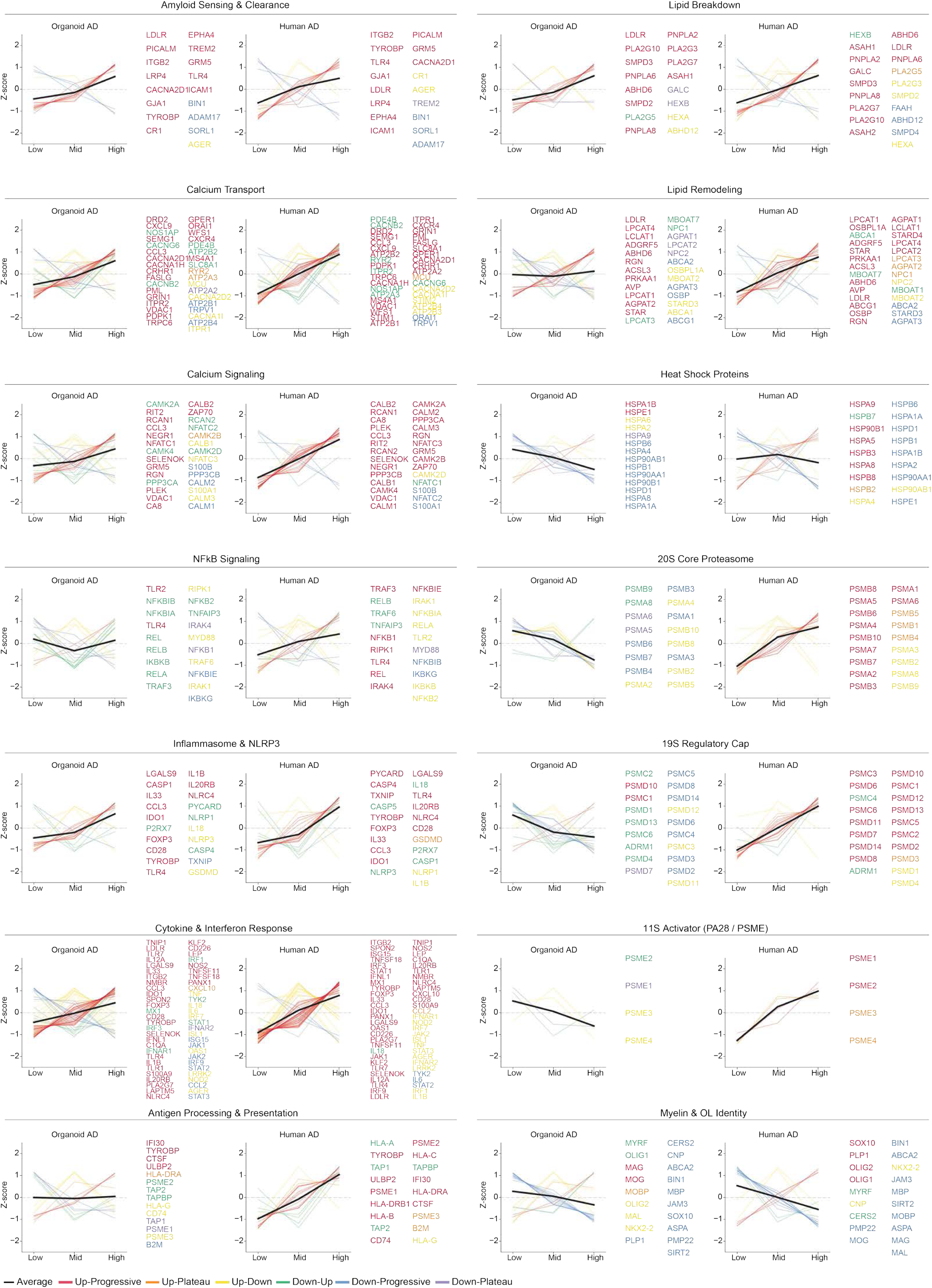
Plaque-associated transcriptional trajectories of canonical gene sets in organoid and human AD oligodendrocytes. Line plots of z-score expression across plaque density tertiles (Low, Mid, High) for curated gene sets representing known biological processes, shown separately for organoid AD (left) and human AD (right) oligodendrocytes. Unlike Figure 5M, which shows only genes with concordant pattern assignments across both models, all genes belonging to each gene set are shown here regardless of their assigned trajectory pattern. Individual gene trajectories are colored by pattern assignment (Up-Progressive, red; Up-Plateau, orange; Up-Down, yellow; Down-Up, green; Down-Progressive, blue; Down-Plateau, purple). The black line indicates the average trajectory across all genes in the gene set. Biological process categories shown: Amyloid Sensing & Clearance, Lipid Breakdown, Calcium Transport, Lipid Remodeling, Calcium Signaling, Heat Shock Proteins, NFkB Signaling, 20S Core Proteasome, Inflammasome & NLRP3, 19S Regulatory Cap, Cytokine & Interferon Response, 11S Activator (PA28/PSME), Antigen Processing & Presentation, and Myelin & Oligodendrocyte Identity.

**Figure S21.**
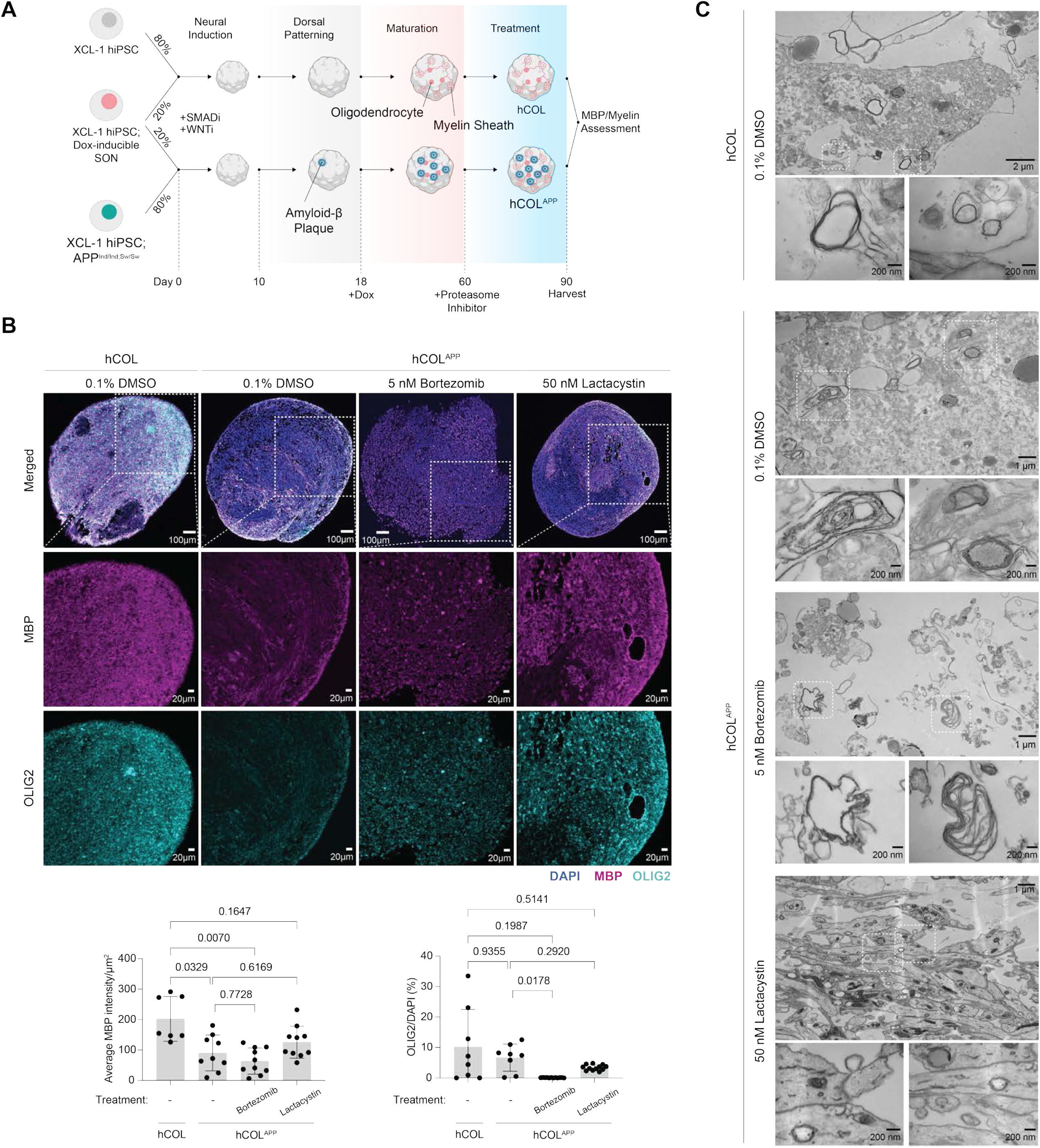
Proteasome inhibition rescues MBP protein levels and myelin ultrastructure in hCOL^APP^ organoids. (**A**) Experimental schematic of hCOL and hCOLAPP development followed by proteasome inhibition with bortezomib and lactacystin. hCOL and hCOL^APP^ organoids were generated from XCL-1 hiPSC lines carrying doxycycline-inducible SON and pathogenic APP mutations, respectively. Doxycycline was added at day 18 to induce oligodendrocyte differentiation and maturation. Beginning at day 60, organoids were treated with 0.1% DMSO vehicle, 5 nM bortezomib, or 50 nM lactacystin for 30 days. Organoids were harvested at day 90 for MBP and myelin ultrastructure assessment via immunofluoresncence and electron microscopy. (**B**) (Top) Representative immunofluorescence images of hCOL (0.1% DMS) and hCOL^APP^ (0.1% DMSO, 5 nM bortezomib, 50 nM lactacystin) organoids stained for MBP (magenta) and OLIG2 (cyan), with DAPI nuclear counterstain. Scale bars: 100 µm (merged), 20 µm (individual channels). (Bottom) Quantification of average MBP intensity per area (µm^2^) (left) and OLIG2+/DAPI ratio (%) (right) across treatment conditions. Each dot represents a region randomly selected. 3 organoids per condition. Borwn-Forsynthe and Welch ANOVA with Dunnett’s T3 multiple comparison test. (**C**) Representative electron microscopy images of myelin ultrastructure in hCOL and hCOL^APP^ organoids with 0.1% DMSO, 5nM bortezomib, or 50 nM lactacystin. Low magnification (top, scale var 1-2 µm) and high magnification overview (bottom, scale bar 200 nm) each condition.

**Figure S22.**
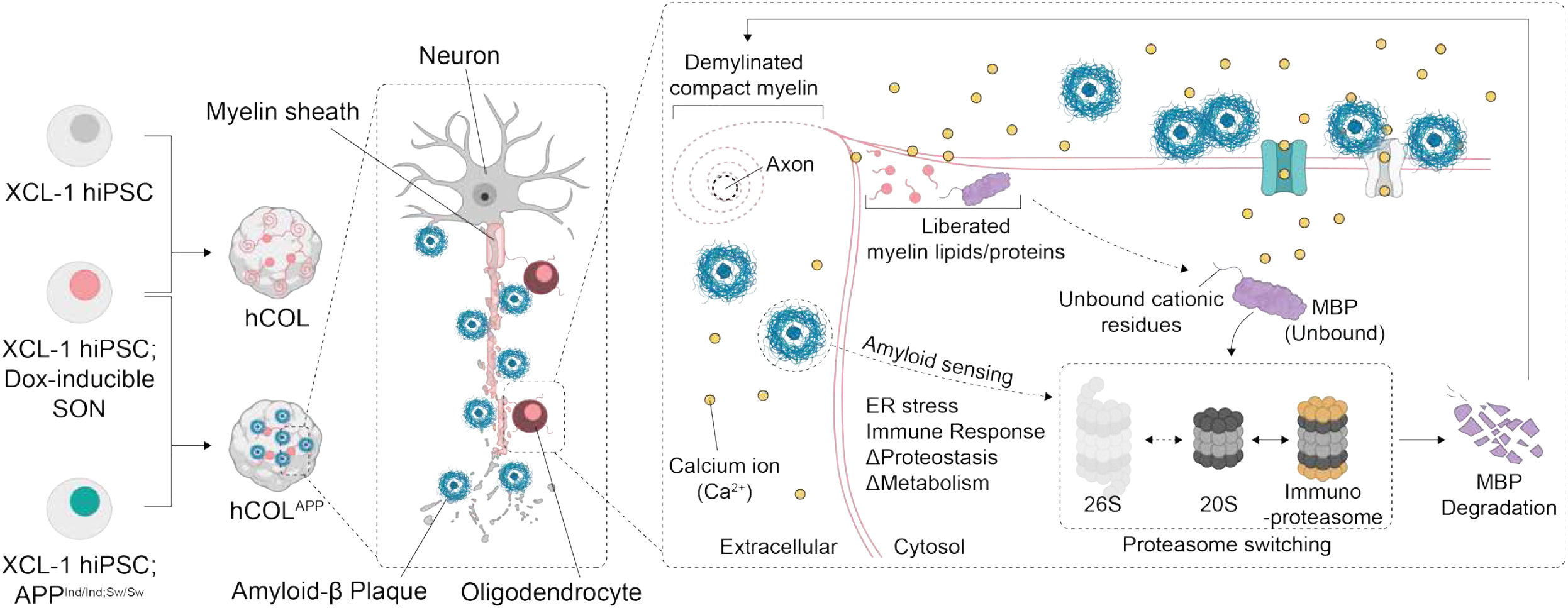
Summary and proposed mechanism of demyelination in Alzheimer’s Disease. Doxycycline-induced expression of SON factors (*SOX10*, *OLIG2*, and *NKX6-2*) enables maturation of oligodendrocytes with myelinating function. In forebrain organoids carrying pathogenic amyloidogenic APP mutations (Swedish/Indiana; hCOL^APP^), myelination is substantially reduced, consistent with myelin loss observed in AD and murine models. In hCOL^APP^, myelin loss is associated with aberrant oligodendrocyte maturation and altered expression of genes involved in lipid/myelin biosynthesis, ER stress responses, immune and inflammatory pathways, and protein catabolism, together with alteration in central metabolism (mitochondrial respiration and major energy metabolism), cytoskeletal maintenance, and autophagy. Based on findings conserved across organoids and human datasets in this study, we propose the following mechanistic model of myelin loss in AD oligodendrocytes: increasing amyloid-β plaque burden in the microenvironment activates intracellular oligodendrocyte stress responses, while impairing autophagy and cellular metabolism. In parallel, plaques may promote intracellular Ca²⁺ overload (for example, by forming membrane pores or activating Ca²⁺ channels), which can destabilize myelin by disrupting interactions between negatively charged myelin lipids and myelin proteins such as MBP. Released lipids may be degraded or remodeled and can contribute to inflammation. Destabilized MBP may become susceptible to ubiquitin-independent proteasomal degradation, potentially via the 20S proteasome. Immune activation can also induce antigen processing and immunoproteasome activity, which may further enhance MBP degradation. Although stressed oligodendrocytes upregulate myelination-related transcripts as a compensatory response, disrupted proteostasis and metabolism limit effective myelin maintenance in an AD-like environment.

**Table S1. Differentially expressed genes between hCOL and hCOL^APP^ oligodendrocytes at day 115**

Differentially expressed genes identified between hCOL and hCOL^APP^ oligodendrocytes at day115 by scRNA-seq using the Wilcoxon rank sum test. Columns: gene symbol; p_val, unadjusted p-value; avg_log2FC, average log2 fold change (positive values indicate higher expression in hCOL^APP^); pct.1, proportion of cells expressing the gene in hCOL^APP^; pct.2, proportion of cells expressing the gene in hCOL; p_val_adj, Bonferroni-adjusted p-value.

**Table S2. Differentially expressed genes between oligodendrocytes subtype 5 and 9 in hCOL and hCOL^APP^**

Differentially expressed genes identified between oligodendrocyte subtype 5 and subtype 9 in hCOL and hCOL^APP^ scRNA-seq using the Wilcoxon rank sum test. Columns: gene symbol; p_val, unadjusted p-value; avg_log2FC, average log2 fold change (positive values indicate higher expression in subtype 5); pct.1, proportion of cells expressing the gene in subtype 5; pct.2, proportion of cells expressing the gene in subtype 9; p_val_adj, Bonferroni-adjusted p-value.

**Table S3. Plaque-associated gene trajectory classification in hCOL and hCOL^APP^ oligodendrocytes**

Plaque-associated trajectory pattern assignments and z-score expression profiles for oligodendrocyte genes in hCOL and hCOL^APP^. Columns: gene symbol; hCOL_pattern, trajectory pattern assigned in hCOL; hCOLAPP_pattern, trajectory pattern assigned in hCOL^APP^; hCOL_low/mid/high, row z-scores for hCOL across plaque density tertiles (Low, Mid, High); hCOLAPP_low/mid/high, row z-scores for hCOL^APP^ across plaque density tertiles. Pattern categories: Up-Progressive, Up-Plateau, Up-Down, Down-Progressive, Down-Plateau, Down-Up.

**Table S4. Gene Ontology enrichment analysis of plaque-associated trajectory patterns in hCOL^APP^ oligodendrocytes**

GO biological process enrichment results for genes assigned to each of the six-plaque density-dependent trajectory patterns (Up-Progressive, Up-Plateau, Up-Down, Down-Progressive, Down-Plateau, Down-Up), stratified by condition specificity. Gene list sheets: Genes_hCOLAPP_only, genes assigned to each pattern exclusively in hCOL^APP^; Genes_hCOL_only, genes assigned exclusively in hCOL; Genes_Shared, genes assigned to the same pattern in both conditions. GO enrichment sheets provide results for each condition-specific and shared gene set per pattern. Columns: ID, GO term identifier; Description, GO term name; GeneRatio, ratio of pattern genes annotated to the GO term; BgRatio, background ratio of all genes annotated to the GO term; RichFactor, gene count ratio relative to background; FoldEnrichment, fold enrichment over background; zScore; pvalue, nominal p-value; p.adjust, BH-adjusted p-value; qvalue; geneID, gene symbols contributing to enrichment; Count, number of genes.

**Table S5. Detailed characterization of human postmortem brain donors and tissue quality metrics for samples used in this study.**

Donor group, study-assigned donor number, BEB ID, research diagnosis, gender, age (years), age at symptom onset (years), disease duration (years), and reported family history of Alzheimer’s disease are listed alongside a brief neuropathology report summary for all 13 donors (6 unaffected control, 7 AD). Alzheimer’s disease neuropathology (NP) is reported using NIA–AA criteria, including the “ABC” scores: A (Thal amyloid phase, 0–5), B (Braak NFT stage, 0–VI), and C (CERAD neuritic plaque score, 0–3), as well as the derived ADNC level (Not/Low/Intermediate/High). Comorbid synucleinopathy variables include PD Braak stage and the presence/absence of diffuse Lewy body disease/Lewy body dementia. Tissue handling metrics include postmortem interval (PMI, hours), brain weight (g), and fixation duration (days). RNA quality metrics include RIN (whole brain; UMBEB) and DV200 (percentage of RNA fragments >200 nt) measured from BA39 extractions (Merck & Co., Inc., Rahway, NJ, USA and GeneWiz). Self-reported race/ethnicity are provided when available; blank indicates not available.

**Table S6. Plaque-associated gene trajectory classification in human AD oligodendrocytes.** Plaque density-dependent trajectory pattern assignments and z-score expression profiles for oligodendrocyte genes in human AD spatial transcriptomics. Columns: gene symbol; Pattern, trajectory pattern assigned in human AD tissue; Low/Mid/High, row z-scores across plaque density tertiles (Low, Mid, High). Pattern categories: Up-Progressive, Up-Plateau, Up-Down, Down-Progressive, Down-Plateau, Down-Up.

**Table S7. Gene Ontology enrichment analysis of plaque-associated trajectory patterns in human AD oligodendrocytes.**

GO biological process enrichment results for genes assigned to trajectory patterns showing significant enrichment in human AD oligodendrocytes. Sheets correspond to patterns with significant GO enrichment: Up-Progressive, Down-Progressive, and Down-Up. Columns: ID, GO term identifier; Description, GO term name; GeneRatio, ratio of pattern genes annotated to the GO term; BgRatio, background ratio of all genes annotated to the GO term; RichFactor, gene count ratio relative to background; FoldEnrichment, fold enrichment over background; zScore; pvalue, nominal p-value; p.adjust, BH-adjusted p-value; qvalue; geneID, gene symbols contributing to enrichment; Count, number of genes.

**Table S8. Overlapping plaque-associated genes between organoid and human AD oligodendrocytes across trajectory pattern categories.**

Plaque density-dependent trajectory pattern assignments and z-score expression profiles for genes shared between organoid and human AD oligodendrocytes. Sheets Organoid_AD and Human_AD provide the full gene lists with pattern assignments and z-scores (Low, Mid, High) for each model separately. Pattern sheets (Up-Progressive, Up-Plateau, Up-Down, Down-Progressive, Down-Plateau, Down-Up) provide the overlapping genes assigned to the same pattern in both models. Columns in pattern sheets: gene symbol; pattern_organoid, trajectory pattern in organoid; Low/Mid/High_organoid, row z-scores in organoid across plaque density tertiles; pattern_human, trajectory pattern in human AD tissue; Low/Mid/High_human, row z-scores in human AD tissue across plaque density tertiles.

**Table S9. Gene Ontology enrichment analysis of plaque-associated genes conserved between organoid and human AD oligodendrocytes.**

GO biological process enrichment results for genes assigned to the same trajectory pattern in both organoid and human AD oligodendrocytes. Sheets correspond to the four pattern categories showing significant conservation: Up-Progressive, Up-Down, Down-Progressive, and Down-Up. Columns: ID, GO term identifier; Description, GO term name; GeneRatio, ratio of conserved pattern genes annotated to the GO term; BgRatio, background ratio of all genes annotated to the GO term; RichFactor, gene count ratio relative to background; FoldEnrichment, fold enrichment over background; zScore; pvalue, nominal p-value; p.adjust, BH-adjusted p-value; qvalue; geneID, gene symbols contributing to enrichment; Count, number of genes.

